# Cerebellar output neurons impair non-motor behaviors by altering development of extracerebellar connectivity

**DOI:** 10.1101/2024.07.08.602496

**Authors:** Andrew S. Lee, Tanzil M. Arefin, Alina Gubanova, Daniel N. Stephen, Yu Liu, Zhimin Lao, Anjana Krishnamurthy, Natalia V. De Marco García, Detlef H. Heck, Jiangyang Zhang, Anjali M. Rajadhyaksha, Alexandra L. Joyner

**Affiliations:** Developmental Biology Program, Sloan Kettering Institute, New York 10065, NY, USA; Neuroscience Program, Weill Cornell Graduate School of Medical Sciences, New York 10021, NY, USA; Bernard and Irene Schwartz Center for Biomedical Imaging, Department of Radiology, New York University Grossman School of Medicine, New York 10016, NY, USA; Department of Biomedical Sciences, University of Minnesota Medical School, Duluth, MN 55812, USA; Center for Cerebellar Network Structure and Function in Health and Disease, University of Minnesota, Duluth, MN 55812, USA; Center for Neurogenetics, Brain and Mind Research Institute, Weill Cornell Medicine, New York 10021, NY 10021, USA; Pediatric Neurology, Department of Pediatrics, Weill Cornell Medicine, New York 10021, NY, USA; Weill Cornell Autism Research Program, Weill Cornell Medicine, New York 10021, NY, USA; Biochemistry, Cell and Molecular Biology Program, Weill Cornell Graduate School of Medical Sciences, New York 10021, NY, USA; Center for Neurotechnology in Mental Health Research, Department of Biomedical Engineering, The Pennsylvania State University, University Park, PA 16801, USA; Center for Substance Abuse Research, Lewis Katz School of Medicine, Temple University, Philadelphia, PA 19140, USA and Department of Neural Sciences, Lewis Katz School of Medicine, Temple University, Philadelphia, PA 19140, USA

**Author notes:** Correspondence to: Alexandra L. Joyner, Developmental Biology Program Sloan Kettering Institute, 1275 York Avenue, Box 511 New York, NY 10065, USA.

## Abstract

The capacity of the brain to compensate for insults during development depends on the type of cell loss, whereas the consequences of genetic mutations in the same neurons are difficult to predict. We reveal powerful compensation from outside the cerebellum when the excitatory cerebellar output neurons are ablated embryonically and demonstrate that the minimum requirement for these neurons is for motor coordination and not learning and social behaviors. In contrast, loss of the homeobox transcription factors Engrailed1/2 (EN1/2) in the cerebellar excitatory lineage leads to additional deficits in adult learning and spatial working memory, despite half of the excitatory output neurons being intact. Diffusion MRI indicates increased thalamo-cortico-striatal connectivity in *En1/2* mutants, showing that the remaining excitatory neurons lacking *En1/2* exert adverse effects on extracerebellar circuits regulating motor learning and select non-motor behaviors. Thus, an absence of cerebellar output neurons is less disruptive than having cerebellar genetic mutations.

## Introduction

The brain has a large capacity to compensate for neuronal loss due to injury when it occurs during development but not in adulthood. In contrast, germline mutations in genes that regulate neural development that result in hypoplasia can have deficits that range from minor to devastating. It is particularly important to understand the causes of behavior deficits in the context of pediatric cerebellar defects, as the degree of recovery of cerebellar function seems to be differentially influenced by the location and extent of insult or the type of genetic mutation^1,2,3,4,5^. As the cerebellum is a complex folded structure housing the majority of the neurons in the brain^6,7^ and many lobules in the cerebellum share functions with another lobule that converges onto similar downstream forebrain targets^8^, the cerebellum is an important structure to study developmental compensation.

The communication between the cerebellum and the rest of the brain is through the downstream cerebellar nuclei (CN) and they contribute to a wide range of motor and non-motor functions^9,10,11,12^. The CN comprise three bilaterally symmetrical nuclei and form a topographic circuit between their presynaptic Purkinje cells (PCs) based on their spatial position within lobules and along the mediolateral axis of the cerebellar cortex^13,14,15,16,17,18,19^. The circuit functions of subregions of the CN have begun to be defined in adult animals by mapping their projections and transiently inhibiting neural activity using targeted viral injections or neuronal ablation^9,10,11,12^. In addition, many studies have manipulated specific lobules in the cerebellar cortex and inferred which CN are involved in regulating motor coordination, motor learning and/or non-motor behaviors based on circuit topography. An additional consideration for ablation studies is that during development, the excitatory neurons of the CN (eCN) play a pivotal role in supporting the survival of PCs which in turn ensure the proper expansion of other cell types in the cerebellar cortex through secretion of Sonic hedgehog^18,20,21^. Thus, growth of each lobule is dependent on the eCN targets of their PCs, and removing a subset of eCN embryonically will impact the development of both their downstream forebrain targets and presynaptic PCs and their local microcircuit. Given the crucial role eCN play in generation and circuit function of the cerebellum, it is important to develop tools to enable manipulations of the same eCN during development and in the adult to define the cerebellar-associated behaviors dependent on the neurons and uncover possible developmental compensation.

We used the medial eCN as a test case for comparing the necessity of a CN subregion during development versus in adulthood. We then determined the baseline requirement for having all eCN intact during development and compare the behavior deficits to mouse mutants lacking developmental genes in the eCN. We identified a *Cre* transgene that targets the medial eCN from embryogenesis onwards. Using this tool we demonstrate that while in adult mice medial eCN activity is required for reversal learning, the brain can compensate for embryonic loss of these neurons alleviating the behavioral deficit. Surprisingly, we find that when all eCN are killed in the embryo there is only a selective impairment in motor coordination behaviors. In contrast, genetic loss of the engrailed (EN1/2) developmental transcription factors in all eCN results in additional deficits in motor learning, acquisition/reversal learning, and spatial working memory, despite only half the eCN dying embryonically. Circuit mapping and diffusion MRI (dMRI) provide evidence for aberrant thalamo-cortical-striatal connectivity as a result of aberrant eCN development.

## Results

### *SepW1-Cre* targets the excitatory neurons in the medial cerebellar nucleus

We chose the medial CN as a region to compare adult inhibition and embryonic ablation of a CN subregion as it can be divided into 4-5 subregions based on transcriptomic profiling and circuit mapping^14,22^. We screened transgenic databases (Gene Expression Nervous System Atlas and Allen Brain Atlas Transgenic Characterization) for a *Cre* driver that within the CN selectively targets the medial eCN. *SepW1-*Cre, a Selenow BAC construct^23^, was found to mediate Cre recombination only in the medial eCN. Immunostaining of cerebellar sections from *SepW1-Cre* mice carrying a Cre-dependent nuclear tdTomato reporter^24^ (*SepW1-Cre; Ai75D*) revealed that within the eCN 92.6% of all tdTomato+ cells are in the medial CN (**Fig. 1a-f**). Moreover, within the medial CN, 100% of tdTomato+ neurons expressed the eCN marker MEIS2+, and 8% of MEIS2+ neurons were not tdTomato+ (**Fig. 1f**). Approximately 50% of NeuN+ GCs in the internal granule cell layer (IGL) also expressed tdTomato (**Supplementary Fig. 1a,b**), as well as all TBR2+ unipolar brush cells (UBCs) (**Supplementary Fig. 1a,c** and ref^25^). Outside the cerebellum, tdTomato+ labeling was restricted to the vestibular nucleus, cerebral cortex, hippocampus, a subpopulation of hypothalamic nuclei and the nucleus of Darkschewitsch (see also Allen Brain Atlas Experiment ID: 488246361). Thus, among the CN neurons, the *SepW1-Cre* transgene selectively labels the eCN of the medial CN. Importantly, Cre is expressed in adult medial eCN, allowing targeting of subpopulations of the neurons in adult mice using viral injections (see below).

**Fig. 1.**
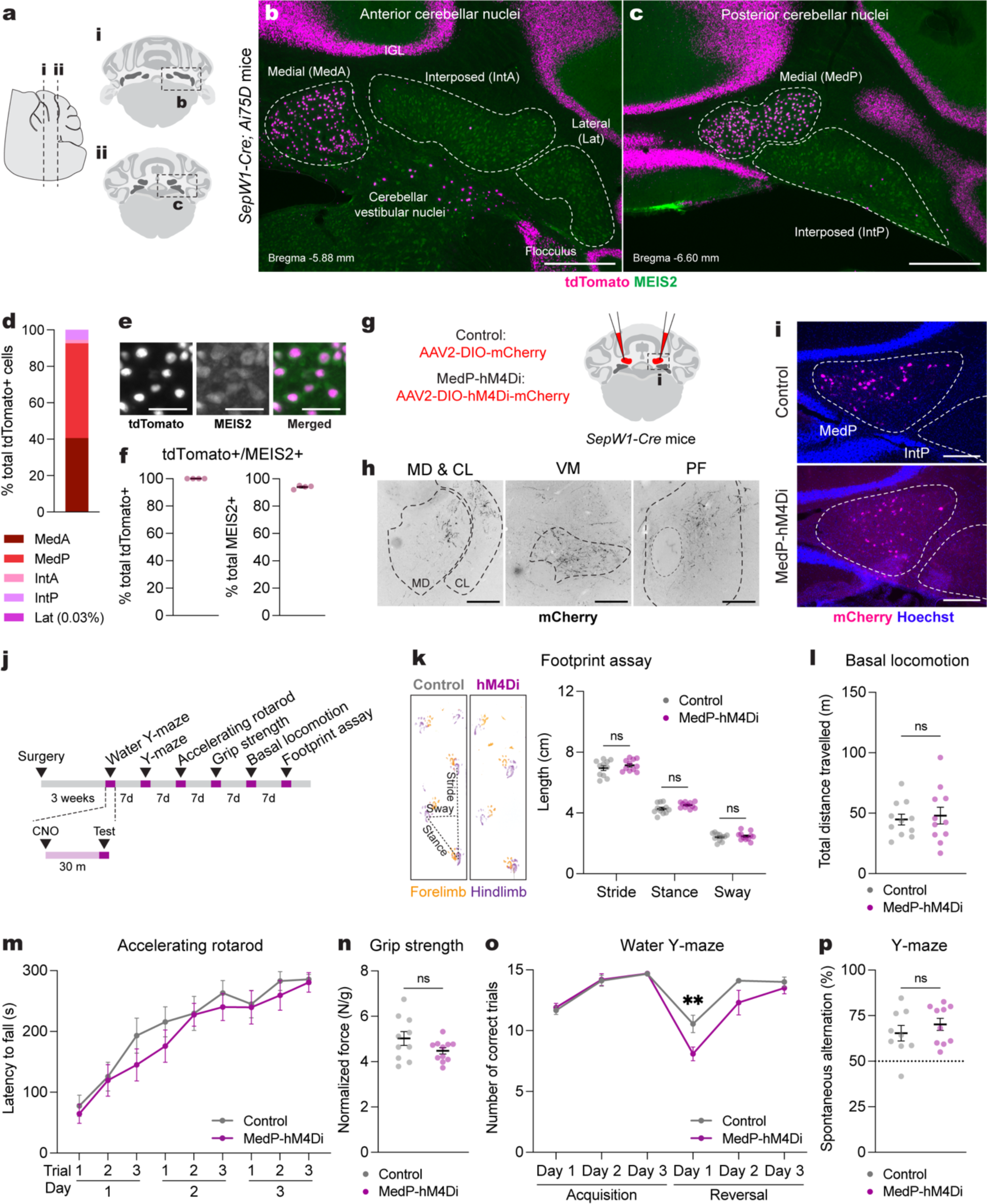
Acute adult chemogenetic inhibition of MedP eCN impairs reversal learning and not motor behaviors. **a**, Schematic representation of a lateral sagittal plane of the mouse cerebellum on the left with vertical lines (**i** and **ii**) indicating the location of the anterior and posterior coronal schematics shown to right. **b**,**c**, Representative coronal images of tdTomato expression in the anterior CN (**b**) and posterior (**c**) CN of *SepW1-Cre; Ai75D* mice. CN were subdivided into five subregions based on histological distinctions (Paxinos and Franklin, 2007) and MEIS2 immunostaining. Abbreviations: MedA=Anterior medial; MedP=Posterior medial; IntA=Anterior interposed; IntP=Posterior interposed; Lat=Lateral. Scale bars = 500 um. **d**, Quantification of tdTomato+ cells on every second coronal section of *SepW1-Cre; Ai75D* mice (n=4) in the lateral CN (Lat) and subregions of the intermediate (Int) and medial (Med) CN (n=4 mice). **e**, Representative image of tdTomato (magenta) and MEIS2 (green) co-expressing eCN in *SepW1-Cre; Ai75D* mice. Scale bars = 50 um. **f**, Quantification of tdTomato+ cells that co-express MEIS2 and the reverse in *SepW1-Cre; Ai75D* mice (n=4 mice). **g**, Schematic of viral injection to express mCherry (control) or hM4Di-mCherry (MedP-hM4Di) in adult MedP eCN. Dashed line indicates region shown in (**i**). **h,** Representative images of MedP eCN mCherry+ axon terminals (black) in four thalamic nuclei of control mice. Fluorescent images were inverted using the look up table in Fiji. Abbreviations: MD=mediodorsal; CL=centrolateral; VM=ventromedial; PF=parafascicular. Scale bars = 250 um. **i**, Representative images of viral mCherry expression in MedP eCN in control (top, AAV-DIO-mCherry) and MedP-hM4Di (bottom, AAV-DIO-hM4Di-mCherry) mice. Scale bars = 250 um. **j**, Experimental timeline of surgery, CNO injection and behavioral tests. **k**, (left) Representative images of footprints from control and MedP-hM4Di mice. (right) quantification of stride, stance, and sway (n=11 per group). Multiple Mann-Whitney *U* tests for effect of genotype on stride (*U* = 48, P = 0.4385), stance (*U* = 34, P = 0.0843) and sway (*U* = 58, P = 0.8851). **l**, Total distance travelled during basal locomotion (n=11 per group; t_20_ = 0.3910, P = 0.7000). **m**, Latency to fall during the accelerating rotarod test (MedP-hM4Di: n=11, control: n=10). Repeated measure two-way ANOVA: main effect of time (F_4.750,90.25_ = 27.51, P < 0.0001), but not of chemogenetics (F_1,19_ = 0.9367, P = 0.3453) or interaction (F_5,152_ = 0.3699, P = 0.9351). **n**, Forelimb grip strength normalized to body weight (MedP-hM4Di: n=11, control: n=10; two-tailed unpaired t-test: t_19_ = 1.677, P = 0.1099). **o**, Total number of correct trials during the water Y-maze test (MedP-hM4Di: n=10, control: n=9). Repeated measure two-way ANOVA: main effect of time (F_5,85_ = 27.65, P < 0.0001) and chemogenetics (F_1,17_ = 9.855, P = 0.006), but not of interaction (F_5,85_ = 2.257, P = 0.0559); with post hoc two-tailed t-tests with Šídák correction for effect of chemogenetics on Reversal Day 1 (t_102_ = 3.386, P = 0.006), and not other comparisons (P ≥ 0.05). **p**, Percentage spontaneous alternations in the Y-maze (MedP-hM4Di: n=10, control: n=9; two-tailed unpaired t-test: t_17_ = 0.9024, P = 0.3794). ns, not significant: P ≥ 0.05. Data are presented as mean values ± SEM.

### Acute inhibition of the adult posterior medial eCN impairs reversal learning and not motor behaviors

Leveraging *SepW1-Cre* mice, we tested the contribution of the posterior region of the medial eCN (MedP eCN) to adult cerebellar-associated motor coordination, learning and non-motor behaviors. Projections to the MedP CN^14,18,26^ that are preferentially from vermis lobules 6-8 have been shown to contribute to cognitive flexibility, anxiolytic and stereotyped/repetitive behaviors^16,27,28^, whereas projections to the anterior medial CN (MedA), which originate from lobules 1-5^17,18^, are associated with motor functions^17,29^. We therefore tested whether acute chemogenetic inhibition of the adult MedP eCN would impair cognitive flexibility without affecting motor functions. A Cre-dependent inhibitory (hM4Di) Gi-DREADD (*AAV2-hSyn-DIO-hM4Di-mCherry*; MedP-hM4Di mice) or control vector (*AAV2-hSyn-DIO-mCherry*; Control mice) was injected bilaterally into the MedP CN of *SepW1-Cre/+* littermates of both sexes (**Fig. 1g**). mCherry and hM4Di-mCherry expression was confirmed to be limited to the MedP (**Fig. 1i** and **Supplementary Fig. 2a,b**) and we observed the expected mCherry+ axon terminals in downstream motor and non-motor thalamic nuclei including mediodorsal (MD), centrolateral (CL), ventromedial (VM) and parafascicular (PF) thalamus (**Fig. 1h**).

The MedP eCN were acutely inhibited during a battery of motor and non-motor behaviors by injecting clozapine N-oxide (CNO, 5 mg/kg) 30 minutes before each behavioral test (**Fig. 1j**). Motor coordination and balance were tested using the footprint assay and revealed no differences in MedP-hM4Di mice compared to littermate controls injected with CNO (**Fig. 1k**). The open field assay also showed no differences in total distance travelled (**Fig. 1l** and **Supplementary Fig. 2c**) and average velocity (**Supplementary Fig. 2d**). Motor performance and learning using an accelerating rotarod test and forelimb grip strength also revealed no differences (**Fig. 1m,n** and **Supplementary Fig. 2e**). In contrast, when we tested acquisition and reversal learning as a readout of cognitive flexibility using a water Y-maze (WYM) test, we found that MedP-hM4Di mice showed normal acquisition learning for finding the submerged escape platform location, but had significantly impaired reversal learning for a new platform location compared to controls (**Fig. 1o**). As a test for spatial working memory, spontaneous alternations in a Y-maze revealed no differences between groups (**Fig. 1p**). Although the MedP-hM4Di mice showed lower total distance travelled (**Supplementary Fig. 2f**) in the Y-maze after CNO administration, the total number of entries (**Supplementary Fig. 2g**) were comparable to control mice. Altogether, these results demonstrated that acute inhibition of the MedP eCN in adult mice selectively impairs reversal learning without having a major effect on motor functions.

### Generation of mice lacking the MedP eCN by conditional knockout of *En1/2*

To examine the impact of embryonic loss of MedP eCN on reversal learning we deleted *En1/2* in the medial eCN using *SepW1-*Cre which initiates recombination at embryonic day 14.5 (E14.5; **Supplementary Fig. 3a-d**), since deletion of *En1/2* in all eCN leads to preferential death of the MedP and posterior interposed (IntP) eCN after E14.5^18^. We confirmed that 7-week-old *SepW1-En1/2* CKO mice of both sexes (*SepW1-Cre; En1^flox/flox^; En2^flox/flox^*) have a loss of medial eCN by quantifying large NeuN (100-600 um^2^) neurons as a proxy for eCN^18^. As predicted, there was a preferential loss of eCN in the posterior medial CN, with little loss in the interposed and lateral CN compared to littermate control mice (*En1^flox/flox^; En2^flox/flox^*)(**Fig. 2a,b**). As recently show, the loss of medial eCN (MEIS2+ cells) was observed by E17.5 and primarily in the MedP region^25^. This was confirmed in 7-week-old *SepW1-Cre; Ai75D* and *SepW1-En1/2 CKO; Ai75D* mice, with complete loss of tdTomato+ MedP eCN and approximately half of the tdTomato+ MedA eCN remaining in the mutants (**Supplementary Fig. 3e**). The CN interneurons were also lost in the MedP region as the only NeuN+ cells remaining in the region were displaced mutant UBCs (**Supplementary Fig. 3e** and ref^25^).

**Fig. 2.**
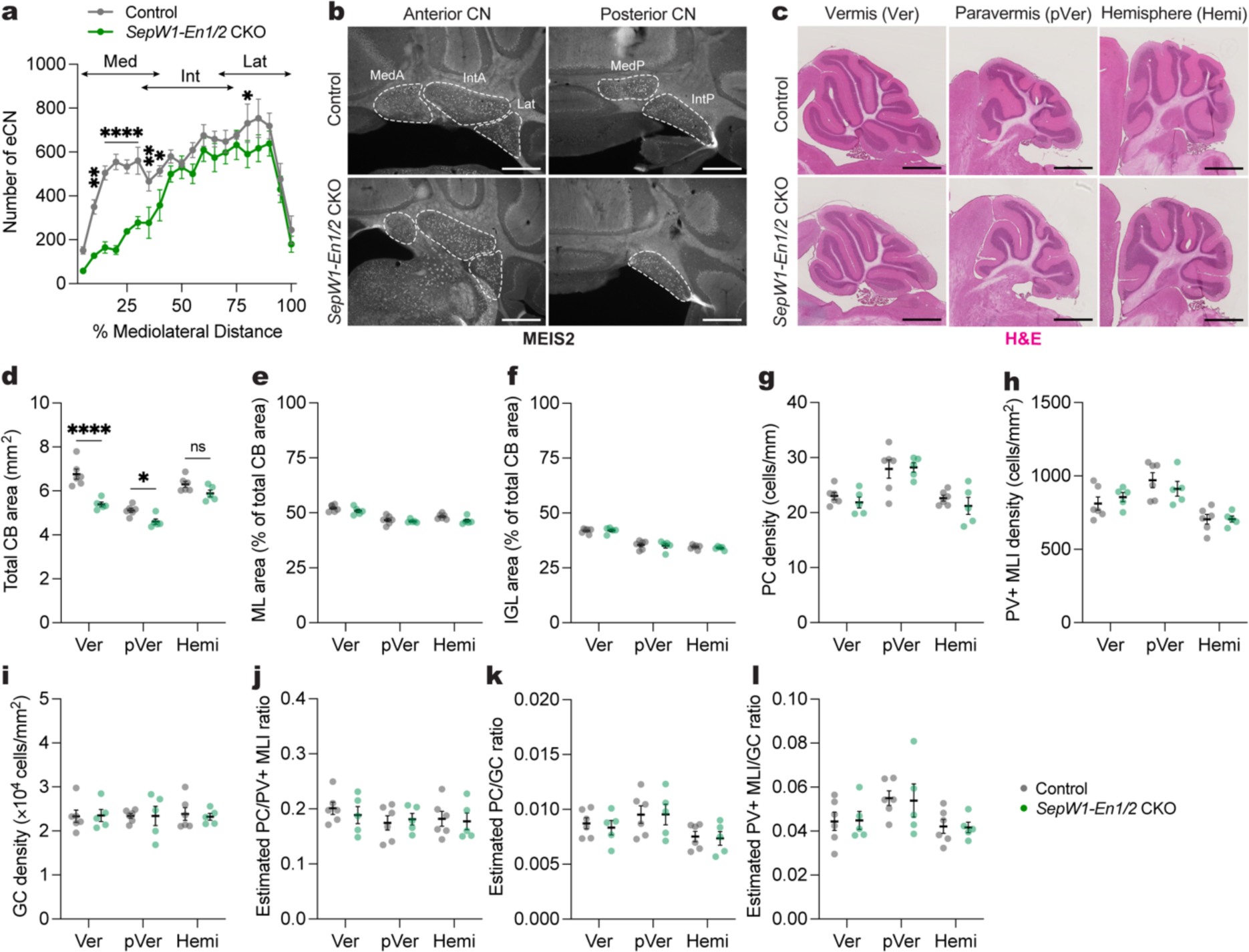
Generation of mice lacking the MedP eCN by conditional knockout of *En1/2*. **a**, Quantification of eCN number (large (100-600 um^2^) NeuN+ cells) along the medial-lateral axis in adult *SepW1-En1/2* CKOs (n=5) and littermate controls (n=6). Ordinary two-way ANOVA: main effect of mediolateral distance (F_19,180_ = 24.02, P < 0.0001), genotype (F_1,180_ = 86.54, P < 0.0001), and interaction (F_19,180_ = 2.449, P < 0.0001); with post hoc two-tailed t-tests with uncorrected Fisher’s LSD for effect of genotype for bin 5-10% (t_180_ = 3.180, P = 0.0017), bins 10-30% (list of t value for each bin: t_180_ = 4.858, 5.738, 4.218, 4.028; all P values: P < 0.0001), bin 30-35% (t_180_ = 2.703, P = 0.0075), bin 35-40% (t_180_ = 2.238, P = 0.0265), bin 75-80% (t_180_ = 2.002. P = 0.0468), but not other comparisons (P ý 0.05). Abbreviations: Med=medial; Int=interposed; Lat=lateral. **b**, Representative coronal images of MEIS2 labeling in the anterior and posterior CN of an adult *SepW1-En1/2* CKO and littermate control as indicated. Scale bars = 500 um. **c**, Representative images of H&E labeled sagittal sections of vermis (Ver), paravermis (pVer), and hemisphere (Hemi) from a *SepW1-En1/2* CKO and littermate control. Scale bars = 1 mm. **d**, Quantification of total cerebellar (CB) area of *SepW1-En1/2* CKOs (n=5) compared to littermate controls (n=6) in the vermis, paravermis and hemispheres. Ordinary two-way ANOVA: main effect of region (F_2,27_ = 42.06, P < 0.0001) and genotype (F_1,27_ = 37.13, P < 0.0001), and interaction (F_2,27_ = 5.825, P = 0.0079); with post hoc two-tailed t-tests with uncorrected Fisher’s LSD for effect of genotype for vermis (t_27_ = 6.292, P < 0.0001), paravermis (t_27_ = 2.359, P = 0.0258), and hemisphere (P = 0.0678). **e**, Quantification of molecular layer (ML) area as a percent of total CB area in *SepW1-En1/2* CKOs (n=5) compared to littermate controls (n=6). Ordinary two-way ANOVA: main effect of region (F_2,27_ = 32.11, P < 0.0001), genotype (F_1,27_ = 5.236, P = 0.0302), but not of interaction (P = 0.5866). **f**, Quantification of internal granule cell layer (IGL) area as a percent of total CB area in *SepW1-En1/2* CKOs (n=5) compared to littermate controls (n=6). Ordinary two-way ANOVA: main effect of region (F_2,27_ = 82.04, P < 0.0001), but not of genotype (P = 0.6943) or interaction (P = 0.8641). **g**, Quantification of Purkinje cell (PC) density in *SepW1-En1/2* CKOs (n=5) compared to littermate controls (n=6). Ordinary two-way ANOVA: main effect of region (F_2,18_ = 18.34, P < 0.0001), but not of genotype (P = 0.4167) or interaction (P = 0.7216). **h**, Quantification of PV+ MLI density in *SepW1-En1/2* CKOs (n=5) compared to littermate controls (n=6). Ordinary two-way ANOVA: main effect of region (F_2,27_ = 16.66, P < 0.0001), but not of genotype (P = 0.8926) or interaction (P = 0.4617). **i**, Quantification of granule cell (GC) density in the IGL of *SepW1-En1/2* CKOs (n=5) compared to littermate controls (n=6). Ordinary two-way ANOVA: no main effect of region (P = 0.9948), genotype (P = 0.8945) or interaction (P = 0.9502). **j**, Quantification of the estimated ratio of the number of PCs to PV+ MLIs in *SepW1-En1/2* CKOs (n=5) compared to littermate controls (n=6). Ordinary two-way ANOVA: no significant main effect of region (P = 0.3807), genotype (P = 0.7618) or interaction (P = 0.7701). **k**, Quantification of the estimated ratio of the number of PCs to GCs in *SepW1-En1/2* CKOs (n=5) compared to littermate controls (n=6). Ordinary two-way ANOVA: main effect of region (F_1,27_ = 4.769, P = 0.0168), but no main effect of genotype (P = 0.7368) or interaction (P = 0.2521). **l**, Quantification of the estimated ratio of the number of PV+ MLIs to GCs in *SepW1-En1/2* CKOs (n=5) compared to littermate controls (n=6). Ordinary two-way ANOVA: main effect of region (F_1,27_ = 4.638, P = 0.0186), but not of genotype (P = 0.9195) or interaction (P = 0.9841). ns, not significant: P ≥ 0.05. Data are presented as mean values ± SEM.

Given that the eCN support the survival of their presynaptic Purkinje cells (PCs), which in turn support production of GCs and interneurons^18^, we confirmed that growth of the cerebellum was reduced in the vermis of 6-week-old animals with proportional scaling down of GCs and interneurons to PCs. There was a significant reduction in the vermis (20.3%) and paravermis (10.1%) sagittal area, but not the hemispheres of *SepW1-En1/2* CKOs compared to littermate controls (**Fig. 2c,d**). In addition, the areas of the molecular layer (ML) and the IGL was primarily reduced in the vermis and their proportions were conserved relative to the total cerebellar area in mutants (**Fig. 2e,f**). The density of Calbindin+ PCs, parvalbumin+ molecular layer interneurons and NeuN+ GCs were also normal throughout the cerebellum (**Fig. 2g-l**). Thus, loss of MedP eCN leads to a preferential reduction of growth in the vermis maintaining the proportions of neurons.

### Mice lacking MedP eCN have normal reversal learning as well as motor behaviors

We next evaluated *SepW1-En1/2* CKOs and their littermate controls of both sexes in the same behavioral assays as those used in the chemogenetic manipulations of *SepW1-Cre* mice (**Fig. 1**). Adult *SepW1-En1/2* CKOs-showed a very small increase in sway length (**Fig. 3a**) with no difference in total distance travelled compared to littermate controls (**Fig. 3b** and **Supplementary Fig. 4a,b**). Additionally no deficits were observed in motor learning and performance between the genotypes (**Fig. 3c,d** and **Supplementary Fig. 4c**). Unlike MedP-hM4Di mice (**Fig. 1o**), *SepW1-En1/2* CKOs showed both normal acquisition and reversal learning compared to littermate controls (**Fig. 3e**). There were no genotype differences in spatial working memory (**Fig. 3f**), number of entries or distance travelled in the Y-maze (**Supplementary Fig. 4d,e**). Motor coordination was also normal in early postnatal pups, using a surface righting reflex assay at P7 (**Fig. 3g**) and negative geotaxis assay at P7 and P11 (**Fig. 3h**). Together, these results demonstrate that embryonic loss of the MedP eCN does not have a major impact on motor and non-motor functions.

**Fig. 3.**
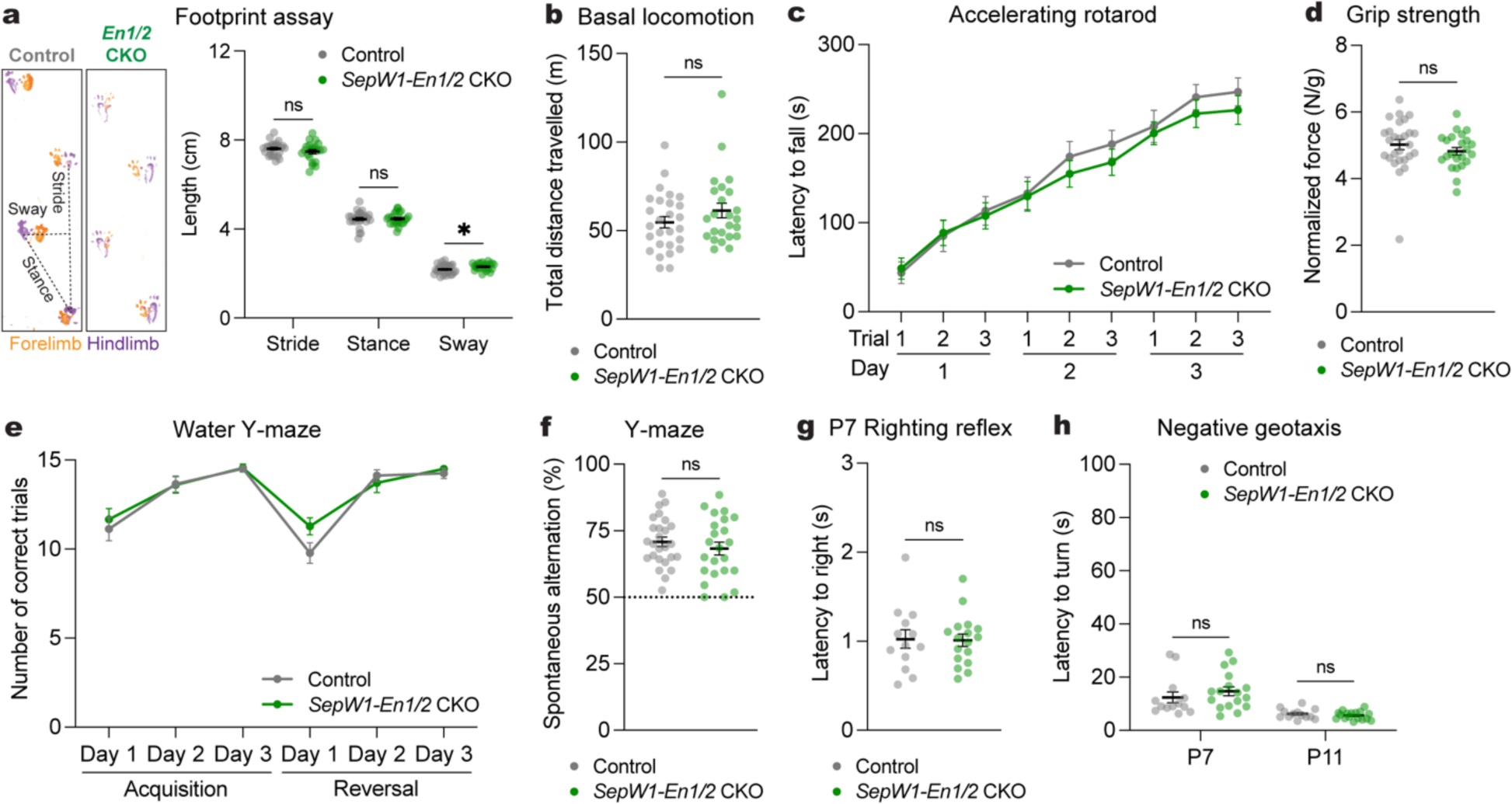
Mice lacking MedP eCN have normal reversal learning as well as motor behaviors. **a**, (left) Representative images of footprints from one *SepW1-En1/2* CKO and littermate control. (right) Quantification of stride, stance, and sway (*SepW1-En1/2* CKOs: n=27, littermate controls: n=22). Multiple Mann-Whitney *U* tests showing effect of genotype on sway (*U* = 188, P = 0.0276), but not stride (*U* = 234, P = 0.2087) or stance (*U* = 279, P = 0.7235). **b**, Total distance travelled during basal locomotion (*SepW1-En1/2* CKOs: n=24, littermate controls: n=27; Mann-Whitney *U* test: *U* = 264, P = 0.2640). **c**, Latency to fall during the accelerating rotarod test (*SepW1-En1/2* CKOs: n=23, littermate controls: n=27). Repeated measure two-way ANOVA: main effect of time (F_4.590,220.3_ = 69.92, P < 0.0001), but not of genotype (F_1,48_ = 0.3434, P = 0.5606) or interaction (F_8,384_ = 0.4280, P < 0.9041). **d**, Forelimb grip strength coronal normalized to body weight (*SepW1-En1/2* CKOs: n=23, littermate controls: n=27; two-tailed unpaired t-test: t_48_ = 1.1018, P = 0.6689). **e**, Total number of correct trials during the water Y-maze test (*SepW1-En1/2* CKOs: n=18, littermate controls: n=23). Repeated measure two-way ANOVA: main effect of time (F_3.132,122.1_ = 38.86, P < 0.0001), but not of genotype (F_1,39_ = 0.5638, P = 0.4572) or interaction (F_5,195_ = 1.492, P = 0.1941). **f**, Percentage of spontaneous alternations in the Y-maze (*SepW1-En1/2* CKOs: n=23, littermate controls: n=26); two-tailed unpaired t-test: t_47_ = 0.8600, P = 0.3942). **g**, Latency to right onto four paws at P7 (*SepW1-En1/2* CKOs: n=17, littermate controls: n=13; two-tailed unpaired t-test: t_28_ = 0.1171, P = 0.9076). **h**, Latency to turn upward on a negative slope at P7 and P11 (*SepW1-En1/2* CKOs: n=17, littermate controls: n=13). Multiple Mann-Whitney *U* tests with Holm-Šídák correction for effect of genotype at P7 (*U* = 79, P = 0.3562) and at P11 (*U* = 92.50, P = 0.4634). ns, not significant: P ≥ 0.05. Data are presented as mean values ± SEM.

### Generation of mice in which all embryonic eCN are ablated using Diphtheria toxin

One plausible explanation for the absence of a reversal learning deficit in mice with embryonic loss of the medial eCN compared to adult inhibition of the same neurons could be the sufficiency of the remaining interposed and lateral CN in the *SepW1-En1/2* CKOs. We therefore used an intersectional pharmacognetic approach to selectively kill all embryonic eCN soon after they are born^30^ as a means to determine the baseline requirement for eCN in a range of adult and neonatal behaviors. Mice were engineered to express Diphtheria toxin subunit A (DTA) in the developing eCN by combining an allele that expresses DTA only in cells that express both Cre and tTA (*lgs7^DRAGON-DTA/+^*; *Atoh1-tTA/+*; *En1^Cre/+^* mice) and administering doxycycline from E13.5 onwards (*eCN-DTA* mice) (**Fig. 4a**). Since the intersection of Cre and tTA expression is in immature eCN and granule cell precursors (GCPs), by administering doxycycline after E13.5 cell killing is limited to the eCN^18^. Immunostaining of sagittal sections at E17.5 revealed that 97.6% of the MEIS2+ eCN were missing in *eCN-DTA* mice compared to littermate controls (*lgs7^DRAGON-DTA^* mice with *Atoh1-tTA* or *En1^Cre^* or neither transgene and fed doxycycline)(36±6.8 cells per mutant vs 1529±95.3 per control) (**Fig. 4b,c**). Quantification of NeuN+ cells in the CN of 7-week-old mice demonstrated a major loss of large CN neurons (100-600 um^2^) in the *eCN-DTA* mice (**Fig. 4d**). RNA *in situ* hybridization analysis confirmed the loss was mainly due to glutamatergic (*Slc17a6*) eCN (**Fig. 4e**). RNAScope triple RNA hybridization analysis (n=3 per genotype) revealed that the rare *Slc17a6-*expressing cells in *eCN-DTA* mice also expressed *Slc32a1* (GABAergic) and *Slc6a5* (glycinergic) markers and were located preferentially in the MedA CN. Furthermore, similar numbers of such triple positive cells were present in controls (**Fig. 4f** and **Supplementary Fig. 5a,b**). In the medial CN both the eCN and interneurons were absent in *eCN-DTA* mice, similar to in *SepW1*-*En1/2* CKOs. We found that the remaining triple positive NeuN+ cells in the MedA CN were not targeted by the *Atoh1-tTA* transgene (**Supplementary Fig. 5c**), despite being targeted with *Atoh1-Cre* (**Supplementary Fig. 5d** and see refs^18,22^). Therefore, all remaining NeuN+ cells in the *eCN-DTA* mice are GABAergic (*Slc32a1+*) inhibitory CN neurons or triple glutamate-GABA-glycine+ neurons.

**Fig. 4.**
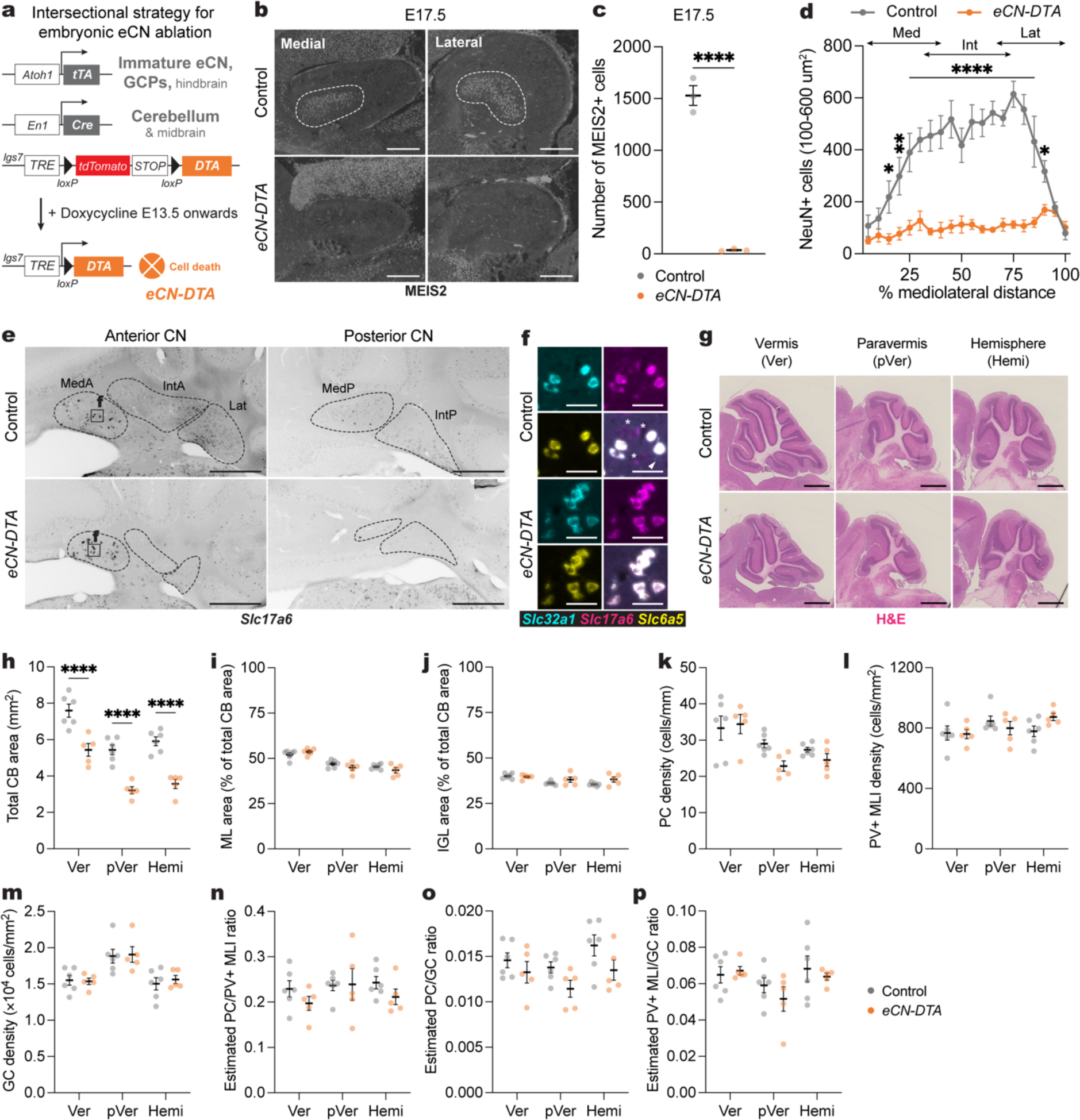
Generation of mice in which all embryonic eCN are ablated using Diphtheria toxin. **a**, Intersectional approach to pharmacogenetically ablate the embryonic eCN. A doxycycline (Dox)-controlled and recombinase activated gene overexpression allele (DRAGON) for attenuated diphtheria toxin fragment A (DTA) (*Igs7^DRAGON-DTA^*) combined with an *Atoh1-tTA* transgene and *En1^Cre^* knock-in allele results in embryonic killing of eCN when Dox is administered starting at E13.5 via expression of DTA. The genotypes of littermate controls are *Atoh1-tTA* or *En1^Cre^* along with the *lgs7^DRAGON-DTA^*allele. **b**, Representative images of sagittal sections stained for MEIS2 (white) in medial and lateral cerebellum of an E17.5 *eCN-DTA* and littermate control. Scale bars = 250 um. **c**, Quantification of total number of MEIS2+ cells on every 10^th^ sagittal section of E17.5 *eCN-DTA* mice (n=3) compared to littermate controls (n=3) (two-tailed unpaired t-test: t_4_ = 15.62, P < 0.0001). **d**, Mediolateral distribution of NeuN+ large cells (100-600 um^2^) in adult *eCN-DTA* mice (n=5) compared to littermate controls (n=6). Repeated measure two-way ANOVA: main effect of mediolateral distance (F_19,180_ = 6.669, P < 0.0001), genotype (F_1,180_ = 359.5, P < 0.0001), and interaction (F_19,180_ = 5.745, P < 0.0001); with post hoc two-tailed t-tests with uncorrected Fisher’s LSD for effect of genotype show significance for bin 10-15% (t_180_ = 3.327, P = 0.0163), bin 15-20% (t_180_ = 4.439, P = 0.0011), bins 20-85% (list of t value for each bin: t_180_ = 4.349, 4.684, 5.539, 5.676, 6.22, 4.587, 5.939, 6.153, 6.76, 6.181, 7.551, 6.744, 4.676; all P values: P < 0.0001), bin 85-90% (t_180_ = 2.229, P = 0.027), but not other comparisons (P ≥ 0.05). Abbreviations: Med=medial; Int=interposed; Lat=lateral. **e**, Representative images of RNA *in situ* analysis of coronal sections for *Slc17a6* expression in the CN of *eCN-DTA* mice and littermate controls. Dotted outlines indicate the CN subregions. Images are single channel inverted using lookup table in Fiji. Scale bars = 500 um. **f**, Representative images of triple RNA *in situ* of coronal sections in the medial CN showing some neurons co-express *Slc32a1*, *Slc17a6*, and *Slc6a5*. Arrowhead and asterisk indicate neurons expressing only *Slc32a1* or *Slc17a6* in controls, respectively. Scale bars = 50 um. **g**, Representative images of H&E labeled vermis (Ver), paravermis (pVer), and hemisphere (Hemi) sagittal sections from an *eCN-DTA* and littermate control. Scale bars = 1 mm. **h**, Quantification of total cerebellar (CB) area in *eCN-DTA* mice (n=5) and littermate controls (n=6) in the vermis, paravermis and hemispheres. Ordinary two-way ANOVA: main effect of region (F_2,27_ = 32.08, P < 0.0001) and genotype (F_1,27_ = 89.64, P < 0.0001), but not of interaction (P = 0.9488); with post hoc two-tailed t-tests with uncorrected Fisher’s LSD for effect of genotype for vermis (t_27_ = 5.260, P < 0.0001), paravermis (t_27_ = 5.425, P < 0.0001), and hemisphere (t_27_ = 5.713, P < 0.0001). **i**, Quantification of molecular layer (ML) area as a percent of total CB area in *eCN-DTA* mice (n=5) compared to littermate controls (n=6). Ordinary two-way ANOVA: main effect of region (F_2,27_ = 37.84, P < 0.0001), but not of genotype (P = 0.3540) or interaction (P = 0.1589). **j**, Quantification of IGL area as a percent of total CB area in *eCN-DTA* mice (n=5) compared to littermate controls (n=6). Ordinary two-way ANOVA: main effect of region (F_2,27_ = 7.249, P = 0.003), but not of genotype (P = 0.0545) or interaction (P = 0.1911); with post hoc two-tailed t-tests with uncorrected Fisher’s LSD for effect of genotype for hemisphere (t_27_ = 2.220, P = 0.0350), but not other comparisons (P ≥ 0.05). **k**, Quantification of PC density in *eCN-DTA* mice (n=5) compared to littermate controls (n=6). Ordinary two-way ANOVA: main effect of region (F_2,18_ = 10.23, P = 0.0011), but not of genotype (P = 0.1660) or interaction (P = 0.2277). **l**, Quantification of PV+ MLI density in *eCN-DTA* mice (n=5) compared to littermate controls (n=6). Ordinary two-way ANOVA: no main effect of region (P = 0.2153), genotype (P = 0.1660) or interaction (P = 0.2277). **m**, Quantification of GC density in *eCN-DTA* mice (n=5) compared to littermate controls (n=6). Ordinary two-way ANOVA: main effect of region (F_2,18_ = 10.08, P = 0.0012), but not of genotype (P = 0.6051) or interaction (P = 0.9220). **n**, Quantification of the estimated ratio of the number of PCs to PV+ MLIs in *eCN-DTA* mice (n=5) compared to littermate controls (n=6). Ordinary two-way ANOVA: no significant main effect of region (P = 0.4484), genotype (P = 0.2061) or interaction (P = 0.5916). **o**, Quantification of the estimated ratio of the number of PCs to GCs in *eCN-DTA* mice (n=5) compared to littermate controls (n=6). Ordinary two-way ANOVA: main effect of genotype (F_1,27_ = 7.013, P = 0.0134), but not of region (P = 0.0913) or interaction (P = 0.7645). **p**, Quantification of the estimated ratio of the number of PV+ MLIs to GCs in *eCN-DTA* mice (n=5) compared to littermate controls (n=6). Ordinary two-way ANOVA: no significant main effect of region (P = 0.0680), genotype (P = 0.4415) or interaction (P = 0.6184). ns, not significant: P ≥ 0.05. Data are presented as mean values ± SEM.

We next examined the size and number of neurons in the cerebellar lobules. As expected, cerebellar size and neuron numbers were reduced across the entire mediolateral axis in *eCN-DTA* mice compared to littermate controls while the proportions and densities of each cell type were maintained (**Fig. 4h-p**) As triple neurotransmitter positive eCN remain in the MedA, we examined the growth of the vermis lobules that preferentially target the MedA. The anterior vermis lobules (1-5) showed a significant reduction in area, but the magnitude was smaller than the reduction in the central vermis (lobules 6-8) (**Supplementary Fig. 6**). The posterior vermis (lobules 9 and 10) was not significantly reduced (**Supplementary Fig. 6**), consistent with the PCs projecting outside the cerebellum to the vestibular nuclei^31,32^. Thus, pharmacogenetic killing of the developing eCN in *eCN-DTA* mice leads to loss of all *Slc17a6* single neurotransmitter positive eCN and reduced cerebellar size with proportional scaling down of cell numbers throughout the cerebellum.

### Loss of all eCN impairs motor coordination, but not motor learning and non-motor behaviors

We repeated the same battery of behaviors in *eCN-DTA* mice of both sexes as in *SepW1-En1/2* CKOs (**Fig. 3**). *eCN-DTA* mice of both sexes had motor coordination deficits in negative geotaxis at P7 and P11 and righting reflex at P7 (**Fig. 5a,b**). Motor coordination continued to be abnormal in adult *eCN-DTA* mice as they had a significant decrease in stride and increase in sway length (**Fig. 5c**) and significantly reduced total distance travelled compared to littermate controls (**Fig. 5d** and **Supplementary Fig. 7a**), but with no difference in average velocity (**Supplementary Fig. 7b**). Interestingly, *eCN-DTA* mice showed normal motor learning (**Fig. 5e,f** and **Supplementary Fig. 7c**). *eCN-DTA* mice showed normal reversal learning, although there was a main effect of geneotype in the WYM (**Fig. 5g**) with no change in swim speed (**Supplementary Fig. 7d**). Moreover, there was no genotype difference in spatial working memory (**Fig. 5h,i**), although the total distance travelled and number of arm entries were decreased in the Y-maze (**Supplementary Fig. 7e-h**).

**Fig. 5.**
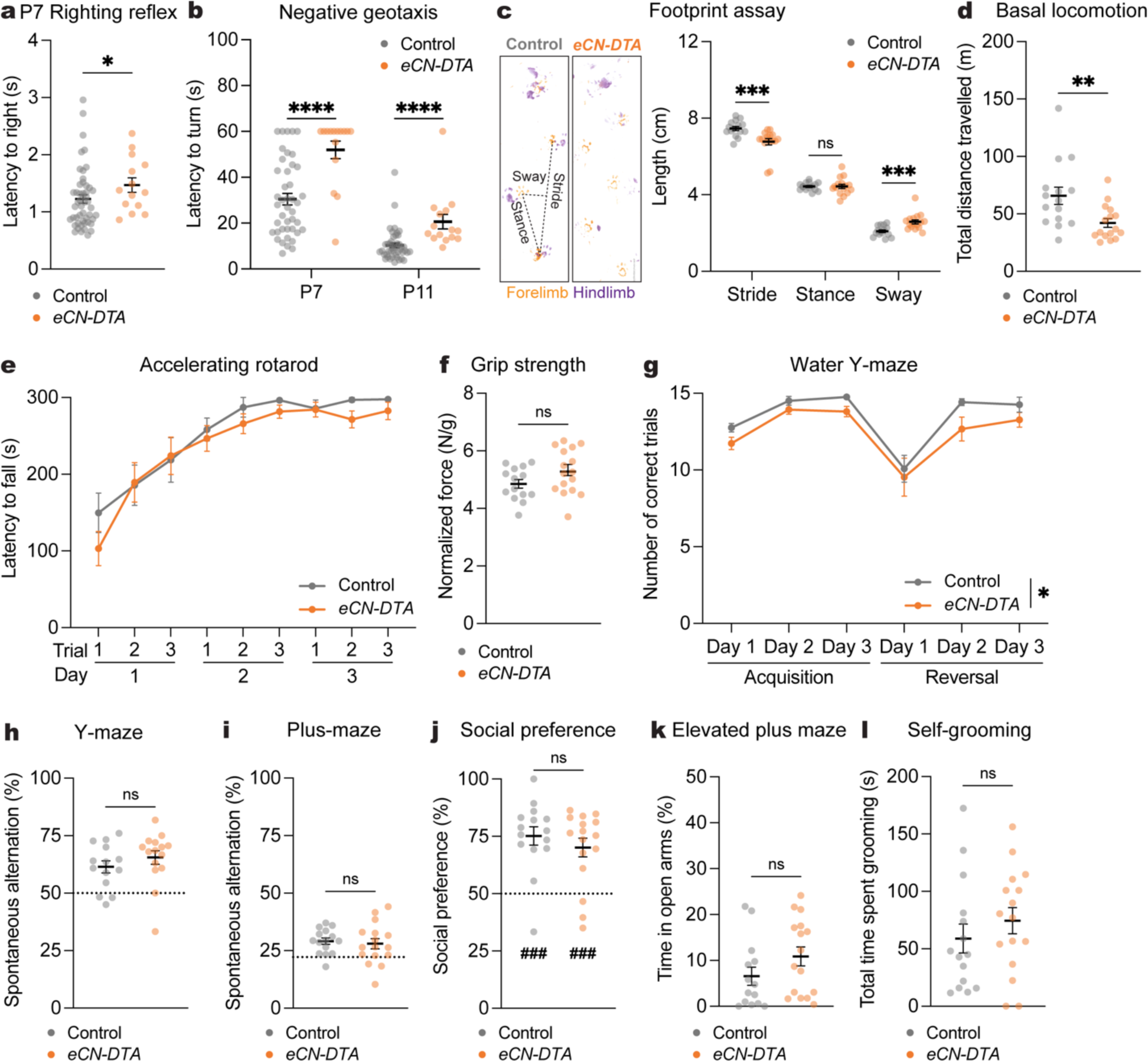
Loss of all eCN impairs motor coordination, but not motor learning and non-motor behaviors. **a**, Latency to right onto four paws at P7 (*eCN-DTA* mice: n=15, littermate controls: n=43; Mann-Whitney *U* test: *U* = 226, P = 0.0436). **b**, Latency to turn upward on a negative slope at P7 and P11 (*eCN-DTA* mice: n=15, littermate controls: n=43). Multiple Mann-Whitney *U* tests with Holm-Šídák correction for effect of genotype at P7 (*U* = 109, P < 0.0001) and at P11 (*U* = 89, P < 0.0001). **c**, (left) Representative images of footprints from an *eCN-DTA* and littermate control. (right) Quantification of stride, stance, and sway (*eCN-DTA* mice: n=16, littermate controls: n=15). Multiple Mann-Whitney *U* tests for effect of genotype on stride (*U* = 33, P = 0.00028) and sway (*U* = 32, P = 0.00023), but not stance (*U* = 113.5, P = 0.8073). **d**, Total distance travelled during basal locomotion (*eCN-DTA* mice: n=16, littermate controls: n=15; two-tailed unpaired t-test: t_29_ = 2.865, P = 0.0077). **e**, Latency to fall in the accelerating rotarod test (*eCN-DTA* mice: n=15; littermate controls: n=15). Repeated measure two-way ANOVA: main effect of time (F_3.426,95.92_ = 34.31, P < 0.0001), but not of genotype (P = 0.3873) or interaction (P = 0.6987). **f**, Forelimb grip strength normalized to body weight (*eCN-DTA* mice: n=16, littermate controls: n=15; two-wailed unpaired t-test: t_28_ = 1.684, P = 0.1033). **g**, Total number of correct trials during the water Y-maze test (*eCN-DTA* mice: n=15, littermate controls: n=12). Repeated measure two-way ANOVA: main effect of time (F_1.667,41.67_ = 17.92, P < 0.0001) and genotype (F_1,25_ = 4.898, P = 0.0362), but not of interaction (P = 0.9183); with post hoc two-tailed t-tests with Šídák correction for effect of genotype all being P ≥ 0.05. **h**, Percentage of spontaneous alternations in the Y-maze (*eCN-DTA* mice: n=15, littermate controls: n=14; Mann-Whitney *U* test: *U* = 76.50, P = 0.2209). Chance level performance is 50% (dotted line). **i**, Percentage of spontaneous alternations in the plus-maze (*eCN-DTA* mice: n=16, littermate control: n=15; two-tailed unpaired t-test: t_29_ = 0.4309, P = 0.6698). Chance level performance is 22.2% (dotted line). **j**, Social preference (percent time nose spent within novel mouse contact zone) during the three-chamber social approach test (*eCN-DTA* mice: n=16, littermate controls: n=15; Mann-Whitney *U* test: *U* = 102, P = 0.4945). Wilcoxon test against a null hypothesis (50%) in *eCN-DTA* mice (*W* = 122, P = 0.0006) and littermate controls (*W* = 116, P = 0.0002). **k**, Percentage of time spent in the open arms of an elevated plus maze (*eCN-DTA* mice: n=16, littermate controls: n=14; Mann-Whitney *U* test: *U* = 74, P = 0.1179). **l**, Total time spent self-grooming (*eCN-DTA* mice: n=16, littermate controls: n=15; Mann-Whitney *U* test: *U* = 92, P = 0.2770). ns, not significant: P ≥ 0.05. Data are presented as mean values ± SEM.

Since all eCN die in the embryo we tested additional cerebellar-associated non-motor behaviors^9,11,12^. In a more challenging spatial working memory test (plus-maze), *eCN-DTA* mice showed no deficits (**Fig. 5i**), although the total distance travelled and number of arm entries were decreased (**Supplementary Fig. 7g,h**). Compared to littermate controls *eCN-DTA* mice showed no difference in social preference after normalizing for hypolocomotion (three-chambered social approach assay^33,34^; **Fig. 5j** and **Supplementary Fig. 7i,j**). We also did not find a genotype difference in anxiety-like behavior (elevated plus maze) when normalizing for hypolocomotion (**Fig. 5k** and **Supplementary Fig. 7k,l**). Finally, we did not find a genotype difference in total time spent self-grooming (**Fig. 5l**). Altogether, the behavior analyses reveal that *eCN-DTA* mice, lacking nearly all eCN, exhibit early motor coordination deficits that persist into adulthood, but motor learning, cognitive, social, and anxiety-like behaviors are largely intact.

### Loss of *En1/2* in all eCN impairs motor coordination and learning, cognitive flexibility, and spatial working memory

Given a previous report that adult mice lacking excitatory activity in all eCN (*Atoh1-Slc17a6* CKO)^5^ have motor learning deficits that we did not observe following loss of all eCN (*eCN-DTA* mice), we were prompted to analyze in more detail the behavior of a previously generated *En1/2* conditional knockout mice (*Atoh1-En1/2* CKOs) that has motor learning deficits^18^. In *Atoh1-En1/2* CKOs all eCN and GCs lack *En1/2* and the mice have an overall ∼50% loss of eCN during embryogenesis that includes a complete loss in the MedP and IntP eCN (**Supplementary Fig. 8a-e** and ref^18^). Further characterization revealed that similar to *SepW1-En1/2* CKOs (**Supplementary Fig. 3e**) there was a significant loss of local interneurons, especially in the MedP and IntP where all neurons are missing (**Supplementary Fig. 8b**).

*Atoh1-En1/2* CKO mice of both sexes had motor coordination deficits in negative geotaxis at P7 and P11 compared to littermate controls (**Fig. 6a**), but no genotype difference in righting reflex at P7 (**Fig. 6b**). Replicating our previous findings^18^, adult *Atoh1-En1/2* CKOs of both sexes had motor coordination (**Fig. 6c**) and motor learning deficits (**Fig. 6e** and **Supplementary Fig. 9c**) and reduced total distance travelled (**Fig. 6d** and **Supplementary Fig. 9a**) and slower velocity (**Supplementary Fig. 9b**) compared to littermate controls. Forelimb grip strength was normal (**Fig. 6f**). Thus, *Atoh1-En1/2* CKOs have motor coordination deficits detected by P7 that persist into adulthood and adult motor learning deficits.

**Fig. 6.**
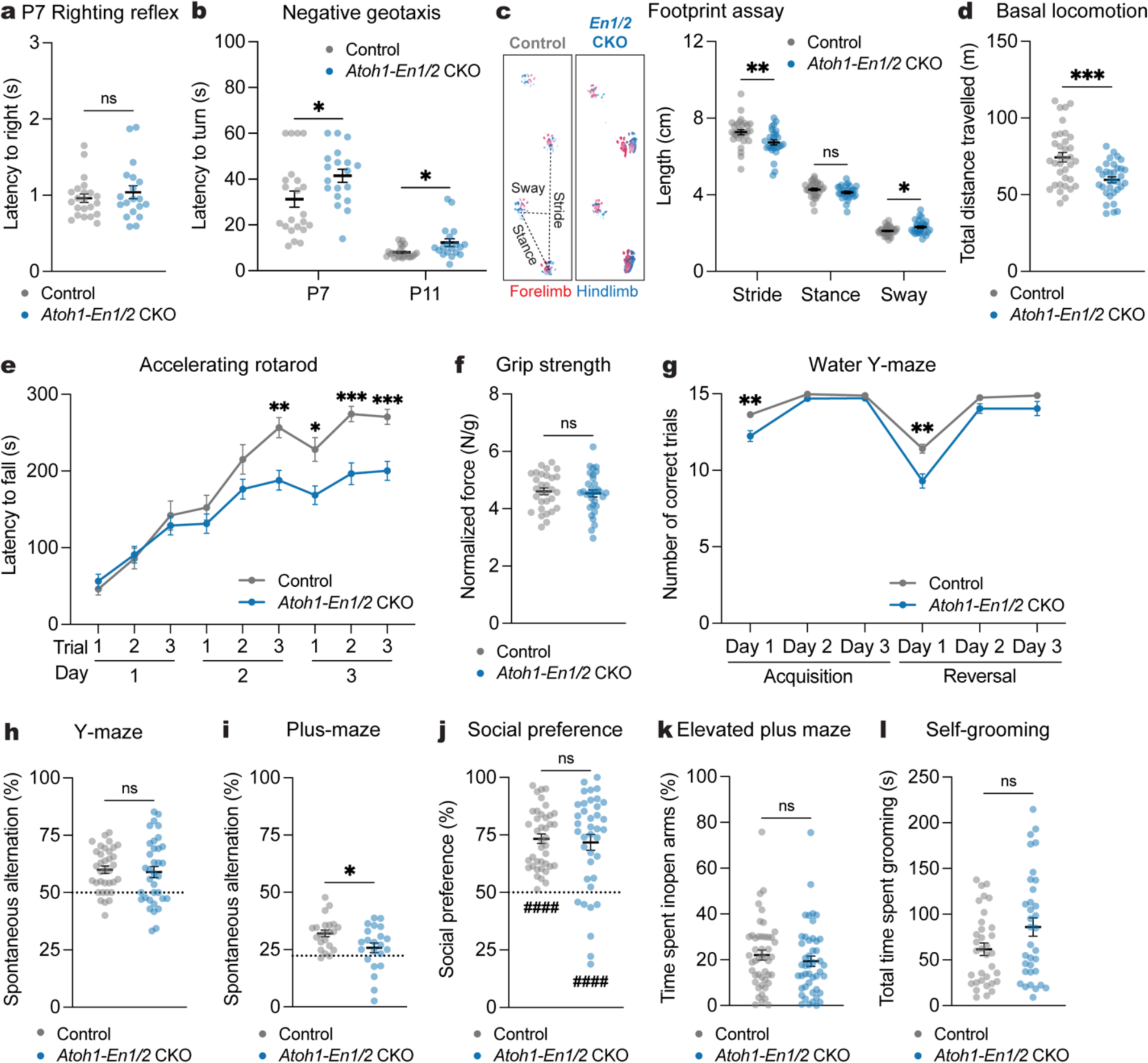
Loss of *En1/2* in all eCN impairs motor coordination and learning, cognitive flexibility, and spatial working memory. **a**, Latency to right onto four paws at P7 (*Atoh1-En1/2* CKOs: n=19, littermate controls: n=22; Mann-Whitney *U* test: *U* = 192.5, P = 0.6741). **b**, Latency to turn upward on a negative slope at P7 and P11 (*Atoh1-En1/2* CKOs: n=19, littermate controls: n=22). Multiple Mann-Whitney *U* tests with Holm-Šídák correction for effect of genotype at P7 (*U* = 128, P = 0.0245) and P11 (*U* = 114.5, P = 0.033). **c**, (left) Representative images of footprints from an *Atoh1-En1/2* CKO and littermate control. (right) Quantification of stride, stance, and sway (*Atoh1-En1/2* CKOs: n=28, littermate controls: n=30). Multiple Mann-Whitney *U* tests for effect of genotype on stride (*U* = 237, P = 0.0039) and sway (*U* = 292.5, P = 0.047), but not stance (*U* = 312, P = 0.0936). **d**, Total distance travelled during basal locomotion (*Atoh1-En1/2* CKOs: n=33, littermate controls: n=35; two-tailed unpaired t-test: t_66_ = 3.931, P = 0.0002). **e**, Latency to fall in the accelerating rotarod test (*Atoh1-En1/2* CKOs: n=32, littermate controls: n=30). Repeated measure two-way ANOVA: main effect of time (F_5.648,338.9_ = 90.56, P < 0.0001), genotype (F_1,60_ = 7.791, P = 0.0070), and interaction (F_8,480_ = 5.827, P < 0.0001); with post hoc two-tailed t-tests with Šídák correction for effect of genotype on day 2-trial 3 (t_59.83_ = 3.721, P = 0.0040), day 3-trial 1 (t_55.69_ = 3.019, P = 0.0338), day 3-trial 2 (t_55.46_ = 4.502, P = 0.0003), day 3-trial 3 (t_57.98_ = 4.416, P = 0.0004), and other comparisons (P ≥ 0.05). **f**, Forelimb grip strength normalized to body weight (*Atoh1-En1/2* CKOs: n=32, littermate controls: n=30; two-tailed unpaired t-test: t_60_ = 0.4298, P = 0.6689). **g**, Total number of correct trials during the water Y-maze test (*Atoh1-En1/2* CKOs: n=31, littermate controls: n=35). Repeated measure two-way ANOVA: main effect of time (F_3.003,192.2_ = 118.4, P < 0.0001), genotype (F_1,64_ = 21.47, P < 0.0001), and interaction (F_5,320_ = 5.101, P = 0.0002); with post hoc two-tailed t-tests with Šídák correction for effect of genotype on Acquisition Day 1 (t_42.50_ = 3.583, P = 0.0052), Reversal Day 1 (t_52.14_ = 3.821, P = 0.0021), and no other comparisons (P ≥ 0.05). **h**, Percentage of spontaneous alternations in the Y-maze (n=35 per genotype; two-tailed unpaired t-test: t_68_ = 0.3622, P = 0.7183). Chance level performance is 50% (dotted line). **i**, Percentage of spontaneous alternations in the plus-maze (n=22 per genotype; two-tailed unpaired t-test: t_42_ = 2.486, P = 0.0170). Chance level performance is 22.2% (dotted line). **j**, Social preference (percent time nose spent within novel mouse contact zone) during the three-chamber social approach test (*Atoh1-En1/2* CKOs: n=38, littermate controls: n=40; Mann-Whitney *U* test: *U* = 723, P = 0.7167). Wilcoxon test against a null hypothesis (50%) in *Atoh1-En1/2* CKOs (*W* = 605, P < 0.0001) and littermate controls (*W* = 820, P < 0.0001). **k**, Percentage of time spent in the open arms of an elevated plus maze (*Atoh1-En1/2* CKOs: n=46, littermate control: n=47; Mann-Whitney *U* test: *U* = 923, P = 0.2266). **j**, Total time spent self-grooming (n=34 per genotype; Mann-Whitney *U* test: *U* = 455, P = 0.1336). ns, not significant: P ≥ 0.05. Data are presented as mean values ± SEM.

In terms of non-motor behaviors, *Atoh1-En1/2* CKOs showed both acquisition (day 1) and reversal learning deficits in the WYM compared to littermate controls (**Fig. 6g**), which was not due to a difference in swim speed (**Supplementary Fig. 9d**). There was no genotype difference in spatial working memory (**Fig. 6h**) despite reduced total distance travelled and arm entries (**Supplementary Fig. 9e,f**). However *Atoh1-En1/2* CKOs showed impaired spatial working memory in the plus-maze (**Fig. 6i**) with no difference in total distance travelled or arm entries (**Supplementary Fig. 9g,h**). *Atoh1-En1/2* CKOs showed no difference in social preference when normalizing for hypolocomotion compared to littermate control mice (**Fig. 6j** and **Supplementary Fig. 9i,j**). Also, genotype differences were not found in anxiety-like behavior (**Fig. 6k** and **Supplementary Fig. 9k,l**) and self-grooming (**Fig. 6l**). Thus, *Atoh1-En1/2* CKOs have impaired adult motor learning, acquisition/reversal learning and spatial working memory, in addition to the motor coordination deficits as seen in *eCN-DTA* mice.

### *Atoh1-En1/2* CKOs have reduced but not ectopic cerebellothalamic projections

One possible explanation for the differing behavioral outcomes in *Atoh1-En1/2* CKOs compared to *eCN-DTA* mice is that the remaining eCN in *Atoh1-En1/2* CKOs exhibit aberrant projections to the thalamus, a primary target region involved in behaviors studied. To address this question, we mapped the projections to the thalamus of the remaining eCN in adult *Atoh1-En1/2* CKO and littermate controls. We first examined whether the remaining MedA and IntA CN make ectopic projections by injecting an *AAV2-hSyn-mCherry* virus or biotinylated dextran amine (BDA) into the MedA or IntA, respectively (n=3 per genotype; **Fig. 7a,d**). Given the reduced neuron number in the MedA and IntA CN, *Atoh1-En1/2* CKOs showed reduced mCherry+ or BDA+ axon terminals in all thalamic nuclei compared to littermate controls (**Fig. 7b,e**), but importantly no obvious ectopic projections were observed compared to littermate controls (**Fig. 7c,f**). We next performed retrograde tracing from the intralaminar thalamic nuclei (ILM), which regulate motor learning and cognitive flexibility^35,36,37,38^. ILM receive strong projections from the MedP, interposed and lateral CN, but not MedA^22^, allowing us to examine whether inputs to ILM from the remaining MedA (and other nuclei) in *Atoh1-En1/2* CKOs are altered. We injected 10% Fluoro-Ruby preferentially into the centrolateral (CL) or parafascicular (PF) thalamus, two nuclei within the ILM, of adult *Atoh1-En1/2* CKOs and littermate controls (**Fig. 7g-i**). As expected, *Atoh1-En1/2* CKOs showed no Fluoro-Ruby+ cells in the region of the MedP and IntP CN. Furthermore, no significant differences in the total number of Fluoro-Ruby+ cells in the MedA, IntA, and lateral CN were detected compared to littermate controls for CL injections (**Fig. 7j**) and PF injections (**Fig. 7k**). Thus, *Atoh1-En1/2* CKOs have reduced cerebellothalamic projections, but the remaining CN do not make ectopic projections.

**Fig. 7.**
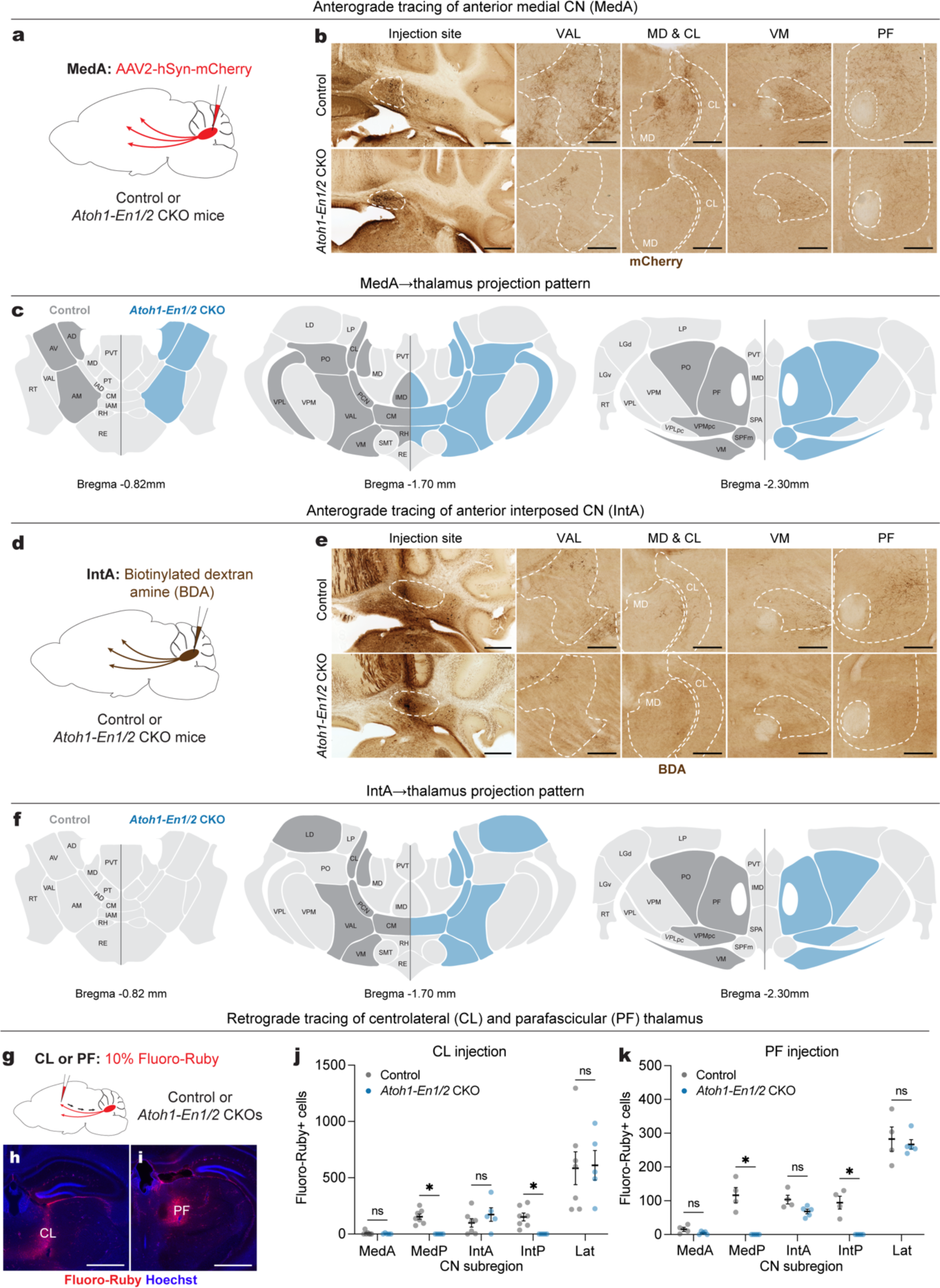
*Atoh1-En1/2* CKOs have reduced cerebellothalamic projections, but no ectopic cerebellothalamic projections. **a**, Schematic of anterograde tracing of MedA CN cells in adult *Atoh1-En1/2* CKOs and littermate controls. **b**, Representative images of coronal sections showing injection site and mCherry+ axon terminals (brown) in various thalamic regions from an *Atoh1-En1/2* CKO and littermate control. Scale bar: injection site = 1 mm and mCherry images = 250 um. **c**, Summary of mCherry+ axon terminals observed in thalamic nuclei of *Atoh1-En1/2* CKOs (blue) versus littermate controls (dark grey) on three representative coronal planes adapted from Allen Brain Atlas. Blue indicates reduced density. **d**, Schematic of anterograde tracing of anterior interposed CN (IntA) cells in adult *Atoh1-En1/2* CKOs and littermate controls. **e**, Representative images of coronal sections showing injection site and biotinylated dextran amine (BDA)+ axon terminals (brown) in various thalamic regions from an *Atoh1-En1/2* CKO and littermate control. Scale bar: injection site = 1 mm and BDA images = 250 um. **f**, Summary of BDA+ axon terminals observed in thalamic nuclei of *Atoh1-En1/2* CKOs (blue) and littermate controls (dark grey) versus on three representative coronal planes adapted from Allen Brain Atlas. Blue indicates reduced density. **g**, (top) Schematic of retrograde tracing in adult *Atoh1-En1/2* CKOs and littermate controls. **h**,**i**, Representative images of the injection site of Fluoro-Ruby (red) and Hoechst (blue) in centrolateral thalamus (CL, **h**) and parafascicular thalamus (PF, **i**). Scale bars = 1 mm. **j**, Quantification of Fluoro-Ruby+ cells in CN subregions that are retrogradely labeled from CL injection. Multiple Mann-Whitney *U* tests with Holm-Šídák correction for effect of genotype on MedP (*U* = 0, P = 0.0126) and IntP (*U* = 0, P = 0.0126), but not on MedA (*U* = 15.5, P = 0.9400), IntA (*U* = 11, P = 0.6882), and Lat (*U* = 15, P = 0.9400). **k**, Quantification of Fluoro-Ruby+ cells in CN subregions that are retrogradely labeled from PF injection. Multiple Mann-Whitney *U* tests with Holm-Šídák correction for effect of genotype on MedP (*U* = 0, P = 0.0390) and IntP (*U* = 0, P = 0.0390), but not on MedA (*U* = 3.5, P = 0.2516), IntA (*U* = 1, P = 0.0922), and Lat (*U* = 10, P > 0.9999). Abbreviations: AD=Anterodorsal nucleus; AM=Anteromedial nucleus; AV=Anteroventral nucleus of thalamus; CL=Central lateral nucleus; CM=Central medial nucleus; IAD=Interanterodorsal nucleus; IAM=Interanteromedial nucleus; IMD=Intermediodorsal nucleus; LD=Lateral dorsal nucleus of thalamus; LGv=Ventral part of the lateral geniculate complex; LP=Lateral posterior nucleus; MD=Mediodorsal nucleus of thalamus; PCN=Paracentral nucleus; PF=Parafascicular nucleus; PO=Posterior complex; PT=Parataenial nucleus; PVT=Paraventricular nucleus; RE=Nucleus of reuniens; RH=Rhomboid nucleus; RT=Reticular nucleus; SMT=Submedial nucleus; SPFm=Subparafascicular nucleus, magnocellular part; VAL=Ventral anterior-lateral complex; VM=Ventral medial nucleus; LGd=Dorsal part of the lateral geniculate complex; VPM=Ventral posteromedial nucleus; VPL=Ventral posterolateral nucleus; SPA=Subparafascicular area; VPMpc=Ventral posteromedial nucleus, parvicellular part; VPLpc=Ventral posterolateral nucleus, parvicellular part. ns, not significant: P ≥ 0.05. Data are presented as mean values ± SEM.

### Diffusion MRI shows *Atoh1-En1/2* CKOs have connectivity changes outside the cerebellum that are distinct from *eCN-DTA* mice

Since the remaining eCN circuits in *Atoh1-En1/2* CKOs appear intact, we used high resolution *ex vivo* dMRI^39,40^ to examine whether there are alterations in regional volume, connectivity, and network properties outside of the cerebellum in adult *Atoh1-En1/2* CKO of both sexes as well as *eCN-DTA* mice for comparison. We first examined regional volume in 10 distinct brain regions (**Supplementary Fig. 10a**) and as expected a large reduction in the volume of the cerebellum was detected in *Atoh1-En1/2* CKO compared to littermate controls (**Fig. 8a**). No changes in regional volumes were seen except for a small but significant reduction in the midbrain (**Fig. 8b**), when normalizing for an overall smaller brain in mutants (**Supplementary Fig. 10b,c**). Network analysis using eight brain regions as nodes revealed a significant reduction in global efficiency and small-worldness in *Atoh1-En1/2* CKOs compared to controls (**Fig. 8c-e** and **Supplementary Fig. 10d-f**). Examining the number of streamlines between the thalamus and CN revealed the expected significant reduction in *Atoh1-En1/2* CKOs compared to controls (**Supplementary Fig. 10g**). We further examined the number of streamlines between the ILM and three of its primary downstream targets; the primary somatosensory cortex (SS), primary motor cortex (MO), and dorsal striatum (DS). Interestingly, we observed a significant increase in the number of streamlines in ILM-SS, ILM-MO, and ILM-DS circuits in *Atoh1-En1/2* CKOs compared to littermate controls (**Fig. 8f-h,k-m** and **Supplementary Fig. 10h-j**). Furthermore, the number of streamlines of the SS-DS and MO-DS circuits were significantly increased in *Atoh1-En1/2* CKOs compared to littermate controls (**Fig. 8i,j,n,o** and **Supplementary Fig. 10k,l**).

**Fig 8.**
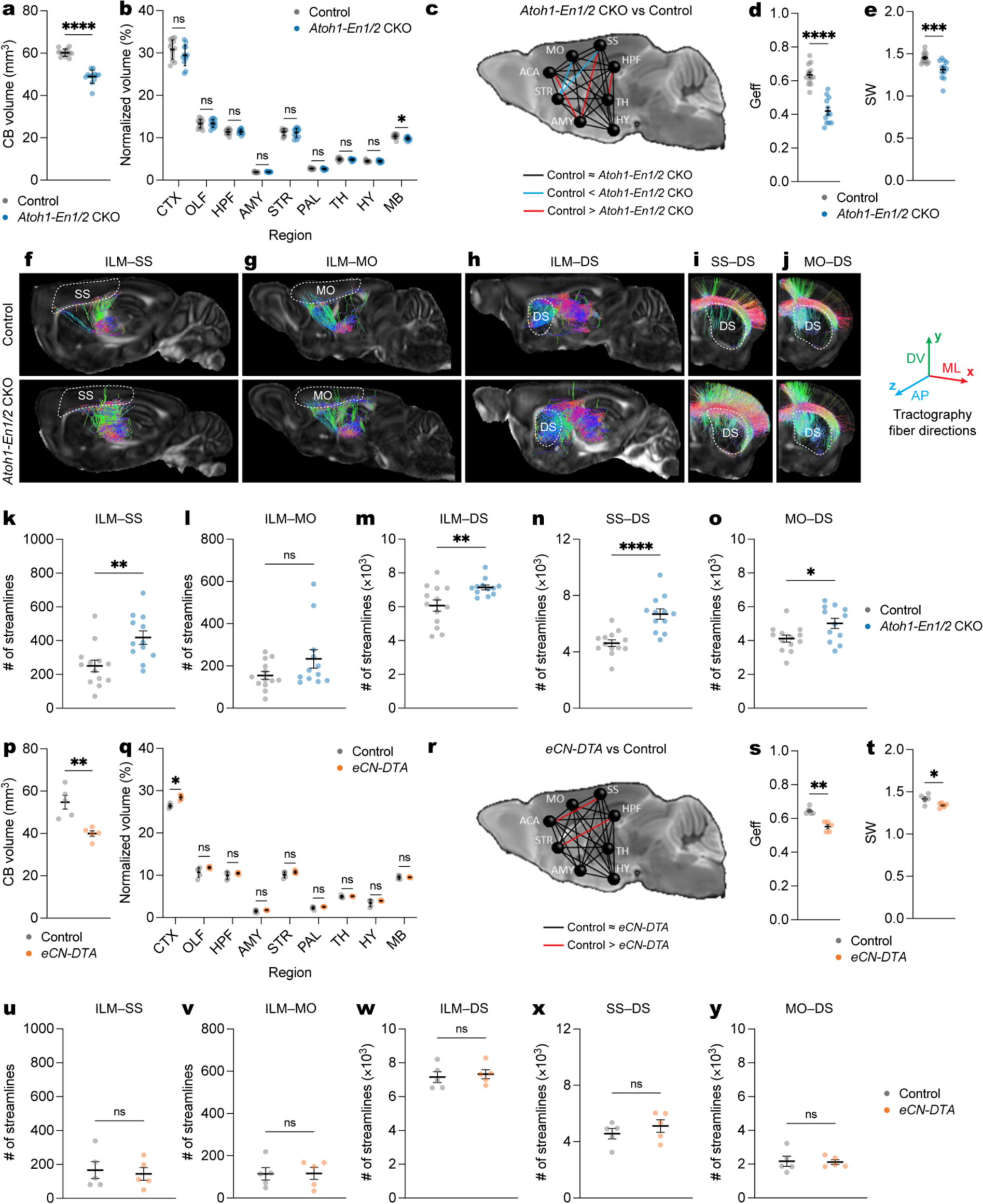
Diffusion MRI shows *Atoh1-En1/2* CKOs have connectivity changes outside the cerebellum that are distinct from *eCN-DTA* mice. **a**, Quantification of cerebellar (CB) volume in *Atoh1-En1/2* CKOs (n=12) compared to littermate controls (n=13) (two-tailed unpaired t-test: t_23_ = 11.15, P < 0.0001). **b**, Quantification of regional volumes normalized to forebrain plus midbrain combined volume in *Atoh1-En1/2* CKOs (n=12) compared to littermate controls (n=13). Two-tailed unpaired t-tests to test for effect of genotype on MB (t_23_ = 2.834, P = 0.0094) and other comparisons P ≥ 0.05. **c**, Schematic representation of global connectivity in *Atoh1-En1/2* CKOs compared to littermate controls. Black lines indicate no significant difference, red lines indicate reduced connectivity in *Atoh1-En1/2* CKOs and blue lines indicate increased connectivity in *Atoh1-En1/2* CKOs compared to littermate controls (two-tailed unpaired t-tests with Welch’s correction). **d**, Quantification of global efficiency (Geff; *Atoh1-En1/2* CKOs: n=12, littermate controls: n=13; two-tailed unpaired t-test: t_23_ = 7.876, P < 0.0001). **e**, Quantification of small worldness (SW; *Atoh1-En1/2* CKOs: n=12, littermate controls: n=13; two-tailed unpaired t-test: t_23_ = 3.913, P = 0.0007). **f-j**, Representative images of ILM-SS (**f**), ILM-MO (**g**), ILM-DS (**h**), SS-DS (**i**), MO-DS (**j**) tractographies in right hemisphere of one *Atoh1-En1/2* CKO and littermate control. Target regions are outlined in dotted lines. The color of streamlines indicates their orientations, as indicated by the colored arrows on the right. Abbreviations: AP=anteroposterior; ML=mediolateral; DV=dorsoventral. **k-o**, Quantification of average (left plus right hemispheres) of ILM-SS tractography (**k**, two-tailed unpaired t-test: t_23_ = 3.225, P = 0.0038), ILM-MO tractography (**I**, two-tailed unpaired t-test: t_23_ = 1.701, P = 0.1024), ILM-DS tractography (**m**, two-tailed unpaired t-test: t_23_ = 2.902, P = 0.0080), SS-DS tractography (**n**, two-tailed unpaired t-test: t_23_ = 4.813, P < 0.0001), and MO-DS tractography (**o**, two-tailed unpaired t-test: t_23_ = 2.515, P = 0.0194) in *Atoh1-En1/2* CKO compared to littermate controls (*Atoh1-En1/2* CKOs: n=12, littermate controls: n=13). **p**, Quantification of cerebellar (CB) volume in *eCN-DTA* mice compared to littermate controls (n=5 per genotype; two-tailed unpaired t-test: t_8_ = 4.295, P = 0.0026). **q**, Quantification of regional volume normalized to forebrain plus midbrain combined volume in *eCN-DTA* mice (n=5) compared to littermate controls (n=5). Two-tailed unpaired t-tests to test for effect of genotype on CTX (t_8_ = 5.876, P = 0.0004) and other comparisons P ≥ 0.05. **r**, Schematic representation of global connectivity for *eCN-DTA* mice compared to littermate controls. Black lines indicate no significant difference and red lines indicate reduced connectivity in *eCN-DTA* mice compared to littermate controls (two-tailed unpaired t-tests with Welch’s correction). **s**, Quantification of global efficiency (Geff; n=5 per genotype; two-tailed unpaired t-test: t_8_ = 2.535, P = 0.035). **t**, Quantification of small worldness (SW; n=5 per genotype; two-tailed unpaired t-test: t_8_ = 1.591, P = 0.1503). **u-y**, Quantification of average (left and right hemispheres) ILM-SS tractography (**u**, two-tailed unpaired t-test: t_8_ = 0.3742, P = 0.7180), ILM-SS tractography (**v**, two-tailed unpaired t-test: t_8_ = 0.0516, P = 0.9601), ILM-SS tractography (**w**, two-tailed unpaired t-test: t_8_ = 0.4177, P = 0.6871), ILM-SS tractography (**x**, two-tailed unpaired t-test: t_8_ = 0.9447, P = 0.3725), and ILM-SS tractography (**y**, two-tailed unpaired t-test: t_8_ = 0.1414, P = 0.8911) in *eCN-DTA* mice compared to littermate controls (n=5 per genotype). Abbreviations: CTX=cerebral cortex; OLF=olfactory bulb; HPF=hippocampal formation; AMY=amygdala; STR=striatum; PAL=pallidum; TH=thalamus; HY=hypothalamus; MB=midbrain; HB=hindbrain; CB=cerebellum; ILM=intralaminar nuclei; SS=primary somatosensory cortex; MO=primary motor cortex; DS=dorsal striatum. ns, not significant: P ≥ 0.05. Data are presented as mean values ± SD for **a**, **j** and mean value ± SEM for **d**,**e**,**k**-**o**,**s-y**.

In *eCN-DTA* mice, in addition to the expected reduction in cerebellum volume (**Fig. 8p**), there was a small but significant increase in the cerebral cortex but no other changes in the rest of the brain (**Fig. 8q**) when normalized to a smaller overall brain compared to littermate controls (**Supplementary Fig. 10m-o**). There was a significant decrease in global efficiency, but no changes in small-worldness in mutants (**Fig. 8r-t** and **Supplementary Fig. 10p,q**). Examining the number of streamlines between the thalamus and CN confirmed a significant reduction in *eCN-DTA* mice (**Supplementary Fig. 10r**). There were no genotype differences in the thalamo-cortico-striatal connectivity that were detected in *Atoh1-En1/2* CKOs (**Fig. 8u-y** and **Supplementary Fig. 10s-w**). Thus, unlike *eCN-DTA* mice, *Atoh1-En1/2* CKOs show extracerebellar changes involving an aberrant increase in connectivity of ILM-cortico-striatal circuits.

## Discussion

Our study applied a novel combination of cell-type-specific genetic manipulations, chemogenetics, dMRI and mouse behavior tests to uncover the requirements during development of the eCN as a whole or the MedP in regulating motor and non-motor functions. We revealed that the main requirement for the eCN if they are all removed in the embryo is for postnatal and adult motor coordination. Furthermore, leveraging an *En1/2* CKO model we demonstrate that seemingly intact eCN that lack critical developmental transcription factors can have major adverse effects on cerebellar and extracerebellar circuits regulating adult motor learning and non-motor behaviors.

We identified a genetic tool (*SepW1-Cre*) to manipulate the medial eCN from E14.5 onwards, adding to existing tools to manipulate all eCN or several subregions^38,41,42,43,44,45,46,47^. By comparing developmental loss (*SepW1-En1/2* CKOs) to adult inhibition of MedP eCN (MedP-hM4Di) we show the neurons are critical for adult reversal learning (WYM) but dispensable if they die embryonically. The MedP-hM4Di results are in line with impaired reversal learning after indirectly inhibiting^27^ or directly activating^16,28^ PCs that preferentially target the MedP eCN (lobule 6-8). Also, inhibition of the adult MedP eCN to ventrolateral periaqueductal gray^48^ or MD^49^ circuits impairs fear extinction, another form of cognitive flexibility. Interestingly, manipulation of other regions like Crus I neurons that target the lateral CN also show reversal learning deficits^27,28^. Furthermore, inhibiting lobule 6 versus Crus I PCs differentially alters c-Fos staining of recruited forebrain regions during reversal learning^28^, likely reflecting distinct downstream pathways^16,22,50^. Therefore, in *SepW1-En1/2* CKOs it is possible that the intact lateral eCN modify their activity to carry out normal reversal learning.

On the contrary, we discovered that when all eCN are ablated embryonically, adult *eCN-DTA* mice show normal reversal learning, indicating that reversal learning must be regulated by extracerebellar brain regions in these mutants. Indeed, regional specific manipulation of the intralaminar thalamus^36^, dorsal striatum^51^, and cerebral cortex^52^ are sufficient to impair reversal learning. Therefore, one possibility is that during development the corticostriatal or thalamostriatal circuits are altered to confer reversal learning without cerebellar input. Similarly, extracerebellar circuits can modulate social preference^53^, spatial working memory^54^, anxiety-like^12,55^, and stereotyped/repetitive^56^ behaviors independent of the CN, which may explain why *eCN-DTA* mice can perform these behaviors.

As expected for a cerebellar-specific developmental perturbation, *eCN-DTA* mice show impaired neonatal and adult motor coordination^5,57,58,59,60^. However, unlike when all eCN activity is inhibited^5^ motor learning is not impaired in *eCN-DTA* mice. In addition to the eCN^58^, the GABAergic CN interneurons play a role in motor learning through their projections to the inferior olive^61^. Therefore, the remaining GABAergic CN interneurons in *eCN-DTA* mutants might contribute to motor learning, as well as extracerebellar circuits^62^ providing compensation. Curiously, these possible sources for compensation in *eCN-DTA* mutants also are expected to apply to mice lacking neurotransmission in all adult eCN^5^. Since *Atoh1^Cre^* was used to generate *Atoh1-Slc17a6* CKO^5^ mice rather than our intersectional approach that targets eCN, one possibility is that one or more of the cell types outside the cerebellum that express *Atoh1^Cre^*^63^ are responsible for the motor leaning deficits in adult *Atoh1-Slc17a6* CKO.

We found that although only ∼50% of eCN are lost in *Atoh1-En1/2* CKOs, adult mutants exhibit impaired motor learning, acquisition/reversal learning, and spatial working memory, which is not seen in *eCN-DTA* mice. Of likely relevance, the remaining eCN in *Atoh1-En1/2* CKOs lack the EN1/2 transcription factors, key regulators of cerebellar development^18,64,65,66,67,68,69,70,71,72,73^. The behavioral deficits in *Atoh1-En1/2* CKOs compared to *eCN-DTA* mice provides direct evidence that dysfunctional eCN circuits due to genetic mutations can have worse outcomes than losing the neurons embryonically. We propose that in *Atoh1-En1/2* CKOs, the remaining eCN while targeting the correct thalamic nuclei are dysfunctional due to altered gene expression. In line with these conclusions, removal of the cerebellum or vermis neonatally in genetically dystonic rats^74,75^ or *weaver* mutants^76,77^ significantly rescues the adult motor coordination deficits seen in both mutants. Circuits that comprise the ILM-cortico-striatal circuitry have been implicated in motor learning^36,37,78^, cognitive flexibility^37,79,80^, and spatial working memory^81^. The excessive connectivity seen in *Atoh1-En1/2* CKOs using dMRI might thus be caused by the remaining *En1/2*-lacking eCN and contribute to their behavioral deficits.

In conclusion, our study highlights the importance of developing relevant models for directly comparing developmental versus adult loss of neurons and the contribution of dysfunctional neurons to understanding behavioral defects and possible compensation (**Supplementary Fig. 11**). Moreover, our findings offer the potential to be leveraged for the development of therapeutic avenues for patients with pediatric cerebellar injuries.

## Methods

### Animals

All animal care and procedures were performed according to the Memorial Sloan Kettering Cancer Center and Weill Cornell Medicine Institutional Animal Care and Use Committee guidelines. Mice were kept in a 12-h/12-h light/dark cycle and in temperature- and humidity-controlled rooms and had *ad libitum* access to standard laboratory mouse chow and water. All transgenic mouse lines were maintained on a mixed genetic background containing 129, C57BL/6J, and Swiss Webster. For behavior analysis, males and females were analyzed separately, but as there were no sex differences all final analyses combined the two sexes. Estrous cycle was not evaluated for females. The following mouse lines were used in the study: *Atoh1-Cre* (JAX #011104)^82^, *tetO-Cre* (JAX #006234)^83^, *R26^LSL-nls-tdTomato^* (Ai75D, JAX #025106)^24^, 129S1/SvImJ (JAX #002448), Swiss Webster mice (Taconic Biosciences catalog #SW), *En1^flox^* (JAX #007918)^71^, *En2^flox^* (JAX #008872)^64^, *Atoh1-tTA*^18^, *En1^Cre^* (JAX #007916)^84^, *lgs7^TRE-lox-tdTomato-STOP-lox-DTA*G^*^128^*^D^* (or *lgs7^DRAGON-DTA^*, JAX #034778)^30^, and *SepW1-Cre* (MMRRC #036190-UCD)^23^. Details of all mouse strains used in the study are listed in **Table 1** and primers used for genotyping are listed in **Table 2**.

**Table. 1.**
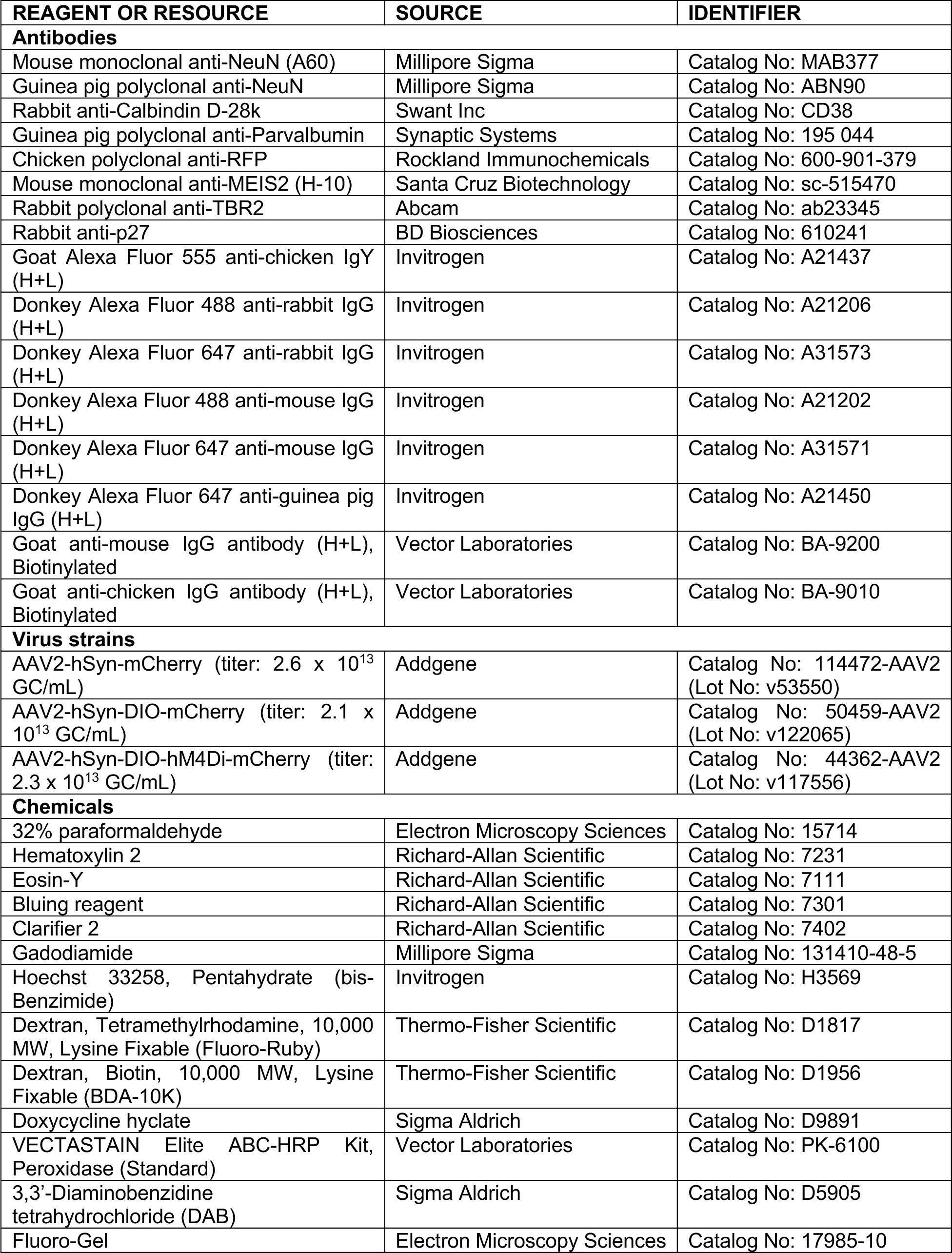
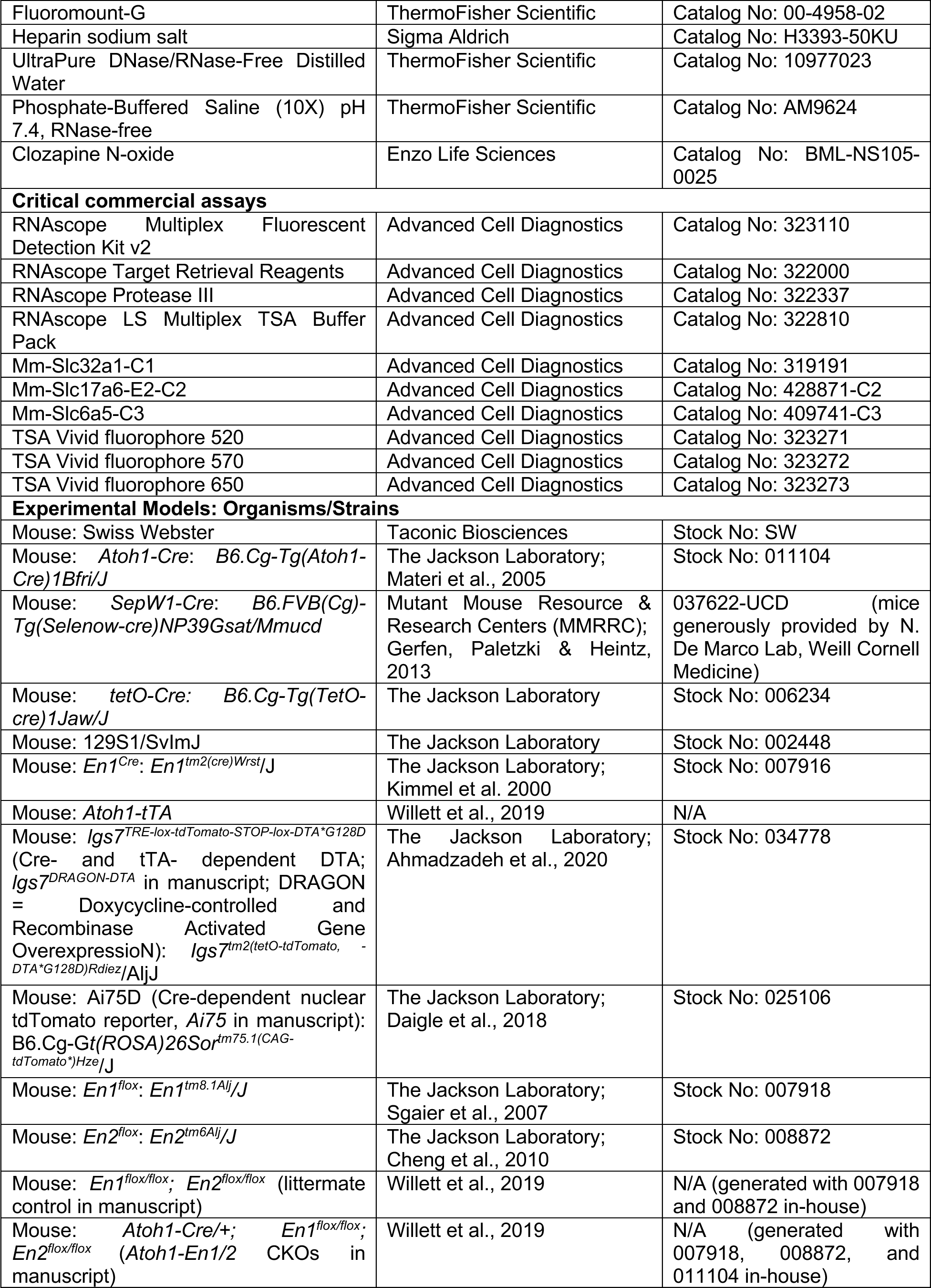
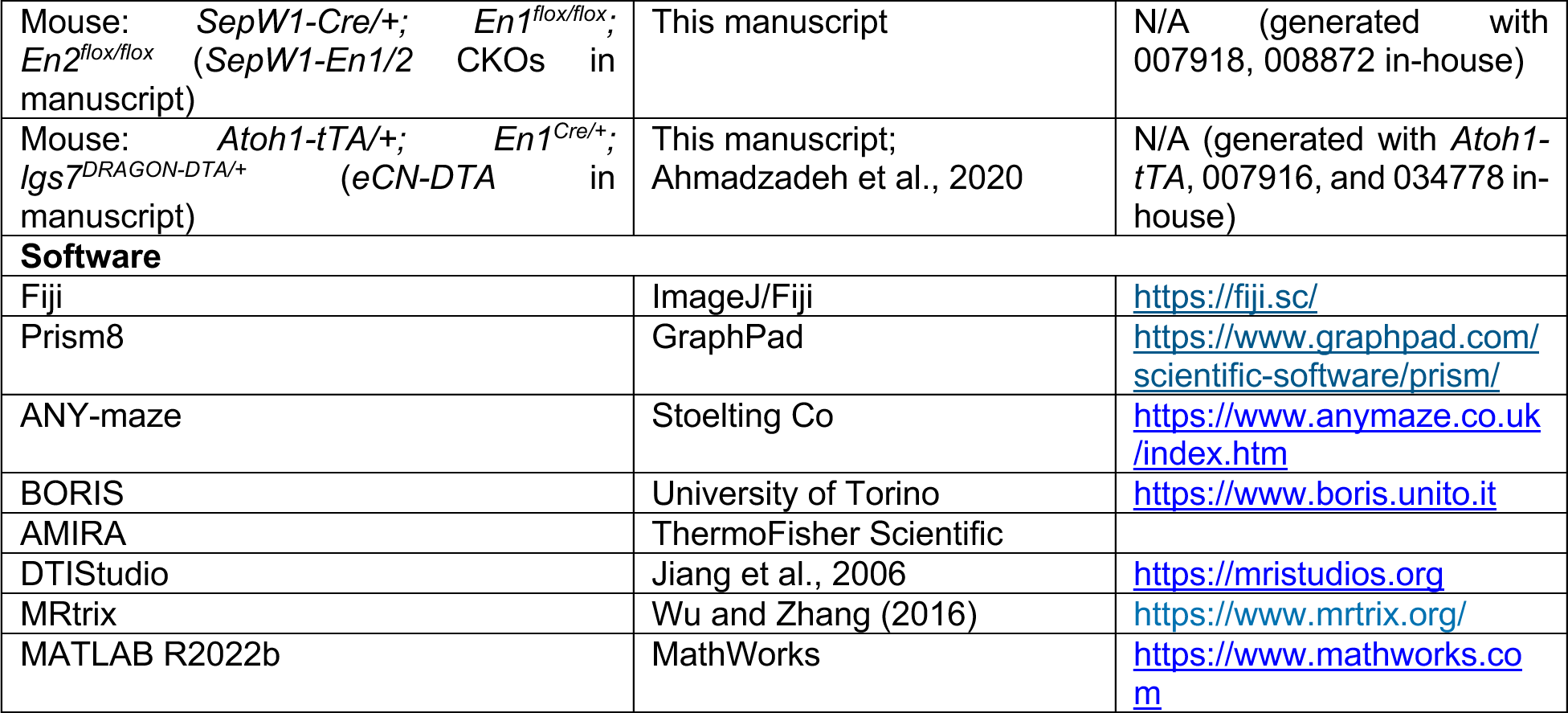
Key resources and sources.

**Table. 2.**
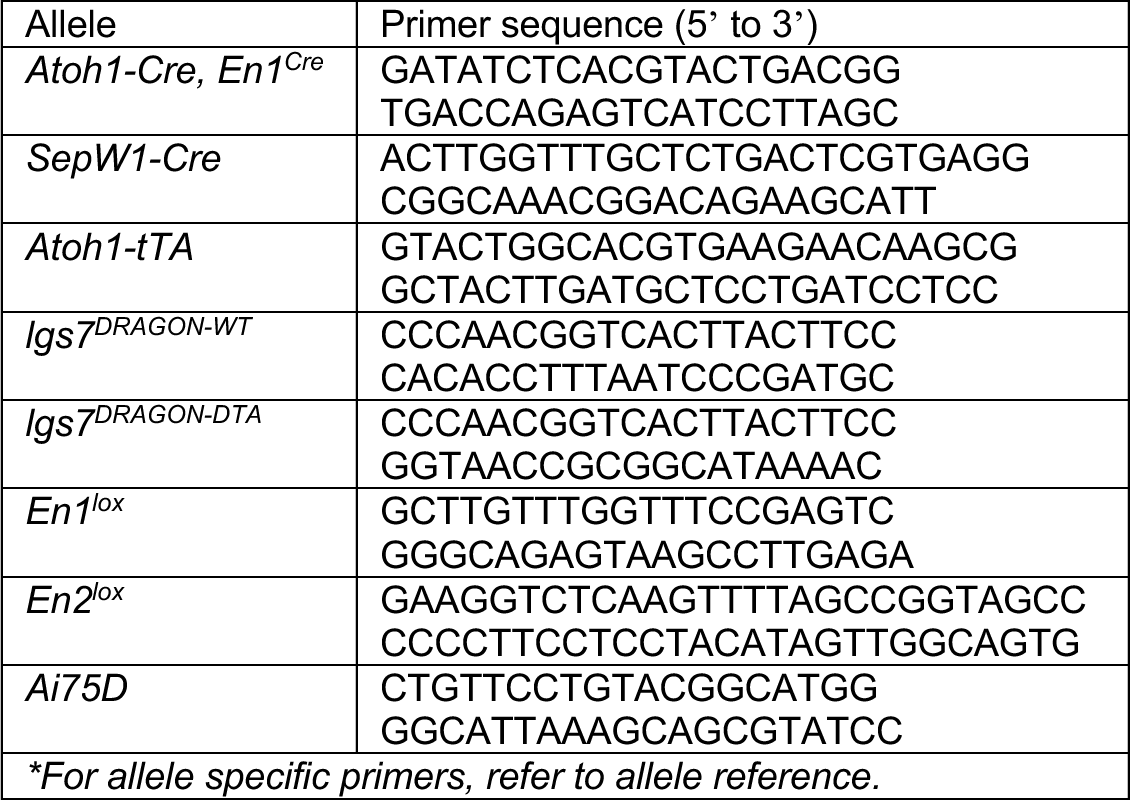
Primers and PCR conditions used for genotyping.

Embryonic ablation of excitatory cerebellar nuclei (eCN) was achieved by crossing *Atoh1-tTA/+; En1^Cre/+^* mice with *lgs7^DRAGON-DTA/DRAGON-DTA^* mice to generate *Atoh1-tTA/+; En1^Cre/+^*; *lgs7^DRAGON-DTA/+^* mice (*eCN-DTA*). Noon of the day that a vaginal plug was discovered was designated as embryonic day 0.5 (E0.5). Doxycycline hyclate (Sigma D9891) was diluted in drinking water (0.02 mg/mL) and provided at E13.5 (when neurogenesis of eCN is complete) until postnatal day 28 (P28) for behavioral studies or otherwise until the end of the experiment and replaced with fresh doxycycline every 3-4 days. To determine if the *Atoh1-tTA* transgene targets the *Slc6a5*-expressing anterior medial eCN, *Atoh1-tTA/+; tetO-Cre/+; R26^LSL-nls-tdTomato/+^* mice were generated and administered doxycycline from E13.5 onwards. *Atoh1-En1/2* CKOs and *Atoh1-En1/2* CKO; *Ai75D* mice were generated by crossing *Atoh1-Cre/+; En1^flox/flox^; En2^flox/flox^* males with *En1^flox/flox^; En2^flox/flox^* or *En1^flox/flox^; En2^flox/flox^; R26^LSL-nls-tdTomato/LSL-nls-tdTomato^* females, respectively. Non-littermate control *Atoh1-Cre; Ai75D* mice were generated by crossing *Atoh1-Cre/+* males with *R26^LSL-nls-tdTomato/LSL-nls-tdTomato^*females. *SepW1-En1/2* CKO and *SepW1-En1/2* CKO; *Ai75D* mice were similarly generated. *SepW1-Cre; Ai75D* mice were generated by crossing *SepW1-Cre/+* or *SepW1-Cre/Cre* males with *R26^LSL-nls-tdTomato/LSL-nls-tdTomato^*females. For all experiments and analyses, investigators were blinded to the genotypes and experimental conditions.

### Behavioral assays

All adult behavioral tests were conducted in a behavioral suite at Weill Cornell Medicine (WCM) and early postnatal behavioral tests were conducted in an animal room at Memorial Sloan Kettering Cancer Center (MSKCC). Six-week-old mice were transferred from MSKCC to WCM two weeks prior to the start of the first behavioral test. Adult mice were acclimated to the animal suite for 1 hour the day before testing. Before conducting each behavioral test, mice were brought to the behavioral suite and left undisturbed for at least 1 hour before testing. The order of each behavior test and age of animals (in brackets) for *eCN-DTA* and *Atoh1-En1/2* CKO mice (and their littermate controls) was as follows: basal locomotor activity^33^ (6 weeks), three-chambered social approach^33,34^ (7 weeks), elevated plus maze^33^ (8 weeks), self-grooming^27^ (9 weeks), accelerating rotarod^18^, grip strength^18^ and footprint assay^18^ (11 weeks), spontaneous alternation in dry Y-maze^33,85^ (12 weeks), water Y-maze^16,59,86^ (13 weeks). Plus-maze^87^ was tested with different groups of mice at 15 weeks (see details below). MedP-hM4Di mice (and control) and *SepW1-En1/2* CKOs (and their littermate controls) followed the same order and age of animals as *eCN-DTA* and *Atoh1-En1/2* CKOs, but were only tested for basal locomotor activity, accelerating rotarod, grip strength, footprint assay, spontaneous alternation, and water Y-maze. For *SepW1-En1/2* CKO, *Atoh1-En1/2* CKO, and *eCN-DTA* mice, negative geotaxis^88^ (at P7 and P11) and surface righting reflex^88^ (P7) were tested. For MedP-hM4Di and control mice, clozapine N-oxide dissolved in 0.9% saline (5 mg/kg, CNO, Enzo Life Sciences) was injected intraperitoneally 30 minutes before the start of each behavioral testing. For accelerating rotarod and water Y-maze, CNO was injected every day of behavioral testing. All behavioral experiments were recorded with a Logitech C920 HD Pro Webcam (30 fps) and analyzed with ANY-maze (Stoelting Co) or hand scored using BORIS software^89^.

#### Negative geotaxis

Mice were tested for negative geotaxis reflex at P7 and P11 as previously described^88^. Mice were placed head down on a negative incline (−35°) platform that was covered with a sterile poly-lined drape. Time until mice turned 180° in either direction was measured using a stopwatch. The test was suspended if mice did not turn within 60 seconds (s) or fell down the slope (considered failed trials). Failed trials were assigned a 60 s latency. All mice were tested three trials per test day.

#### Surface righting reflex

Mice were tested for surface righting reflex at P7 as previously described^88^. Mice were placed in the supine position on a flat surface with a sterile poly-lined drape. The time taken for each mouse to turn onto their four paws was measured. All mice were tested for three trials per test day.

#### Basal locomotor activity

Mice were placed in a polycarbonate test chamber (27.3 cm x 27.3 cm) equipped with three infrared beam arrays. Horizontal locomotor activity was monitored by computer-assisted activity monitoring software (Med Associates). For each test session, animals were placed in the chamber and recorded for 1 hour without interruption with no incandescent lighting^33^. Locomotor activity was measured as total distance traveled in centimeters.

#### Three-chamber social approach

Mice were tested with a modified version of the three-chamber social approach and social novelty assays as described previously^33,34^. All testing was conducted in the three-chamber apparatus (Ugo Basile sociability apparatus, Stoelting Co) in a room with 30 lux lighting at the center of each chamber (1∼2 lux difference across chambers) and a ceiling-mounted camera for ANY-maze tracking. Two days before testing, age- and sex-matched 129S1/SvImJ mice were habituated to the wire cup (see ref^34^ for details). After 1-hour habituation to the testing room, two 129S1/SvImJ mice were placed individually under a wire cup (3.8 cm bottom diameter, rust-proof/rust-resistant, noncorrosive, steel wire pencil cups) in the left or right chamber of the apparatus. The entrance to each chamber was blocked. 129S1/SvImJ mice were observed for 10 min for behaviors that are potentially disruptive, such as bar-biting, excessive self-grooming, circling, or clinging to the side bars with all four paws. Only 129S1/SvImJ mice showing docile behavior were used as novel mice in the social approach and novelty testing.

On the day of testing, mutant or control mice were placed in the center of an empty apparatus to freely explore all three chambers for 10 min. During the 10 min habituation, empty wire cups were placed in both left and right chamber. Mice were briefly taken out of the apparatus and a novel object (orange rectangle block) and a novel mouse were placed under each wire cup. The location of the novel mouse was randomly assigned across each subject mouse. Test mice were placed back into the center chamber and allowed to freely explore for 5 min. Test mice were kept in a separate cage until all animals from its original home cage were tested. Time spent in each chamber and time spent in the contact zone (nose within a 2 cm radius around the wire cups) were calculated by automated detection.

#### Elevated plus maze

Mice were placed in the center of an elevated plus maze (L x W x H = 50 cm x 5 cm x 50 cm and 38 cm above the floor) for 10 min with 15 lux lighting in the open arms and 5 lux lighting in the closed arms^33^. A ceiling-mounted camera for ANY-maze tracking was used to measure time spent in and entries into each arm.

#### Self-grooming

Mice were placed in a clean cage for 10 min to habituate to the test arena with 25-30 lux lighting^27^. After habituation, mouse activity was recorded for 10 min. Time spent grooming was hand scored by an experimenter blind to the genotype.

#### Accelerating rotarod and grip strength

Mice were tested using an accelerating rotarod protocol as previously described^18^ with 30 lux lighting. Mice were put on a rotarod (47650, Ugo Basile) rotating at 10 rpm until all mice (3∼5 mice tested simultaneously) were facing forward for at least 5 s. The rod was then accelerated from 10 rpm to 40 rpm over 5 min. Mice were tested for 3 trials per day across 3 consecutive days. Animals rested for 10 min in their home cage between each trial. Latency to fall (seconds) was recorded for each animal. If a subject mouse held onto the rod and rotated 3 consecutive times, the latency at which the mouse first started to rotate with the rod was recorded as the latency to fall and the mouse was returned to their home cage. If a subject mouse repeatedly fell within 10-15 s after the start of each trial, the mouse was excluded from the final analysis. If mice did not fall throughout the entire trial, 300 s was assigned as latency to fall. The following day after the accelerating rotarod test, forelimb grip strength was measured using a horizontal grip bar (1027SM Grip Strength meter with single sensor, Columbus Instruments). Mice were allowed to hold the triangle grip bar while being gently pulled away by the base of their tail with their body parallel to the bench. The average of 5 measurements was normalized to the mouse’s body weight (force/gram). Animals rested 5 min in their home cage between each measurement.

#### Footprint assay

Mice were tested using a footprint assay as previously described with 30 lux lighting^18^. After forepaws and hindpaws were painted with different nontoxic acrylic paint colors (red/blue or orange/purple, Crayola), mice were allowed to walk through a plexiglass tunnel (L x W x H = 50 cm x 10 cm x 15 cm) lined with a strip of paper. A dark box was placed at the end of the plexiglass tunnel. Each mouse was tested three times, and three gait parameters (stride, sway, and stance) were measured from each run and averaged.

#### Y-maze

Mice were tested using a Y-shaped maze with equal length arms (L x W x H = 33.0 cm x 7.6 cm x 38.1 cm) as described previously^33,85^ with 20 lux lighting at the bottom of the center of the arena. Mice were placed in the center of the arena and allowed to freely explore the arena for 5 min. A ceiling-mounted camera for ANY-maze tracking was used to record the distance travelled in the maze. The sequence of arm entries and number of entries to each arm was manually scored. The percentage of spontaneous alternation was calculated as previous described^33,85^.

#### Water Y-maze

Mice were tested using an adapted version of the water Y-maze test as described previously^16,59,86^ with no incandescent lighting (red light for experimenter). The same Y-shape maze used in the dry Y-maze test above was filled with room temperature water colored with non-toxic white paint to be opaque. After dry Y-maze testing and ∼1 week prior to testing, mice were placed at the base of one of the Y-maze arms (filled with paint but no hidden platform) and allowed to freely swim for 3 minutes. Average velocity was measured using ANY-maze tracking software. On day 1, mice were placed in the base of one arm of the Y-maze and allowed to swim for 60 s without a platform present to further habituate the mice to the arena. On the next three days (2-4), a white platform was submerged about 1 cm below the surface of the water at the end of one of the arms of the Y-maze. The location of the platform was randomly assigned to either the left or right arm relative to the starting arm (base arm). Mice were placed in the base of the Y-maze and latency to find the platform was recorded. The number of correct trials were recorded only when the first arm entered was the arm with the platform and the mouse found the platform without entering another arm. Once the mouse found the platform, the mouse was allowed to sit on it for 15 s and removed from the Y-maze and placed under a heat lamp in an empty cage. If the mouse did not find the platform within 35 s, it was placed onto the platform for 20 s. All mice were run for the same trial before repeating the process for all 15 trials. On average 10-15 mice were run during one session. This procedure was repeated with each animal for 15 trials per day for 3 consecutive days. For days 5-7, the location of the platform was switched to the opposite arm from which the mouse was initially trained on days 2-4. The same procedure as days 2-4 were repeated. Each day, at the end of the set of 15 trials, the mouse was toweled dry and placed under a heat lamp for 5 min before returning to its home cage. The correct choices (when the first arm entered was the correct arm and the mouse found the platform) and latency to find platform were hand scored using a stopwatch during each trial.

#### Plus-maze

Mice were tested using a plus-shape maze with equal length arms (L x W x H = 30.48 x 5.08 x 15.24 cm) as described previously^87^ with incandescent lighting. Mice were placed in one arm of the maze designated the base of the arena and allowed to freely explore the arena for 12 min each day for a total of 4 days (day 1 was considered habituation to the arena). A ceiling-mounted camera for ANY-maze tracking was used to record the distance travelled in the maze. The sequence of arm entries and number of entries into each arm was scored using a custom MATLAB script. The percentage spontaneous alternation was calculated as previous described^87^ and the last 3 days of the behavioral testing was averaged for final analysis. As previously described, the estimated chance level performance for the plus maze test is 22.2% spontaneous alternations^87^. For *Atoh1-En1/2* CKOs, mice were tested in two different locations: Weill Cornell Medicine (New York, NY) and University of Tennessee Health Science Center (Memphis, TN). Data from both institutions were pooled as the results were consistent. For *eCN-DTA* mice, all mice were tested at Weill Cornell Medicine (New York, NY).

### Stereotaxic Injections

Mice were anesthetized with 2.5% isoflurane and 80% O_2_ and head-fixed in a stereotaxic frame (David Kopf Instruments, Tujunga, CA) equipped with digital manipulator arms (Stoelting Co, Wood Dale, IL). A nose cone was used to deliver isoflurane to maintain anesthesia. Mice were given a subcutaneous injection of meloxicam (2 mg/kg) and 0.1 mL of 0.25% Marcaine around the incision site. After exposing the skill, craniotomies were made with an electric drill (Stoelting CO, Wood Dale, IL) with a ball bur attached. A Neuros 7000 series 1 µL Hamilton syringe with a 33-gauge needle (Reno, NV) connected to a remote automated microinfusion pump (KD Scientific, Holliston, MA) was used for delivery of tracers or viral constructs at a rate of 100 or 50 nL/min, respectively. Details of dye, viruses, and stereotaxic coordinates are provided in **Table 3**. Following infusion, the needle was left in place for 5 min and then slowly manually retracted. Mice were placed on a heat pad for at least 30 min post-surgery before being returned to their home cage. Mice were monitored for three days post-surgery to ensure recovery. Brains were collected and imaged (see below) to confirm viral expression and injection placements and no misplacements were found.

**Table. 3.** Details of dye and viruses used in stereotaxic surgeries.

| Tracer/virus | Serotype | Company | Catalog no. | Concentration/titer | Stereotaxic coordinate (distance from Bregma) | Injection volume |
| --- | --- | --- | --- | --- | --- | --- |
| Dextran, Tetramethylrhodamine, 10,000 MW, Lysine Fixable (Fluoro-Ruby) | n/a | Thermo-Fisher Scientific | D1817 | 10% | CL: AP, -1.20 mm; ML, 0.69 mm; DV, -3.8 mm<br>PF: AP, -2.1 mm; ML, 0.6 mm; DV, -3.5 mm | CL: 300 nL<br>PF: 100 nL |
| Dextran, Biotin, 10,000 MW, Lysine Fixable (BDA-10K) | n/a | Thermo-Fisher Scientific | D1956 | 10% | IntA: AP, -6.24 mm; ML, 1.10 mm; DV, -3.80 mm | 300 nL |
| pAAV-hSyn-mCherry | AAV2 | Addgene | 114472-AAV2 (Lot No: v53550) | $2.6 \times 10^{13}$ GC/mL | MedA: AP, -6.24 mm; ML, 0.85 mm; DV, -3.70 | 200 nL |
| pAAV-hSyn-DIO-mCherry | AAV2 | Addgene | 50459-AAV2 (Lot No: v122065) | $2.1 \times 10^{13}$ GC/mL | MedP: AP, -6.7 mm; ML, $\pm 0.9$ mm; DV, -3.20 | 300 nl |
| pAAV-hSyn-DIO-hM4Di-mCherry | AAV2 | Addgene | 44362-AAV2 (Lot No: v117556) | $2.3 \times 10^{13}$ GC/mL | MedP: AP, -6.7 mm; ML, $\pm 0.9$ mm; DV, -3.20 | 300 nl |

### Tissue preparation

For all histological analyses, mice were anesthetized with isoflurane and then perfused transcardially with room temperature (RT) phosphate buffered saline (PBS) with heparin (0.02 mg/mL) and then ice-cold 4% paraformaldehyde (PFA, Electron Microscopy Sciences, catalog no: 15714). Brains were dissected and post-fixed in 4% PFA overnight at 4°C and cryopreserved in 30% sucrose in PBS for ∼2 days at 4°C. Brains were embedded in Tissue-Tek OCT compound (Sakura Finetek). Coronal serial cryosectioning was performed at either 40 um (6 series) or 100 um (3 series) and collected in 0.01M phosphate buffer (PB) with 0.02% sodium azide and kept at 4°C until further processing. For long-term storage, sections were transferred to 24-well plates with cryoprotectant (30% sucrose and 30% ethylene glycol in 0.1M PB) and stored at −20°C. Sagittal serial cryosectioning was performed at either 14 um (10 series) or 30 um (6 series) and collected on charged glass slides (Fisherbrand ColorFrost Plus) and stored at −20°C until further processing. Details of reagents are listed in **Table 1**.

For single molecular fluorescence *in situ* hybridization, tissue was prepared as previously described^90^. Mice were anesthetized with isoflurane and then perfused transcardially with RT 0.9% saline with 10 units/mL heparin (Sigma Aldrich, H3393-50KU) followed by RT 4% PFA in 0.2M phosphate buffer (PB). Brains were dissected and post-fixed in the same fixative overnight at RT. Brains were cryopreserved by incubating in 15% sucrose in 0.2M PB overnight followed by an overnight incubation in 30% sucrose in 0.2M PB. Brains were embedded in Tissue-Tek OCT compound and stored at −80°C until further processing. Coronal serial cryosectioning was performed at 30 um and sections were stored in cryoprotectant solution at −20°C until further processing. Details of reagents are listed in **Table 1**.

### Hematoxylin and eosin (H&E) staining

All reagents for H&E staining were obtained from Richard-Allan Scientific: hematoxylin 2 solution (catalog no: 7231), eosin-Y (catalog no: 7111), bluing reagent (catalog no: 7301), clarifier 2 (catalog no: 7402). Slides were first washed in PBS for 5 min and incubated in hematoxylin 2 solution for 3 min. Slides were rinsed in deionized water (diH_2_O) and immersed in staining reagents for 1 min each (diH_2_O, bluing reagent, diH_2_O, clarifier 2, diH_2_O). After dehydration and defatting (dH_2_O-70% ethanol-95%–95%-100%-100%-xylene-xylene-xylene, 1 min each) slides were mounted with a coverslip and DPX mounting medium (Electron Microscopy Sciences). Details of reagents are listed in **Table 1**.

### Immunofluorescence

For slide-mounted sections, slides were washed three times in PBS for 5 min. If necessary, antigen retrieval was then performed by immersing slides in sodium citrate buffer (10 mM sodium citrate with 0.05% Tween-20, pH 6.0) at 95°C for 20 min followed by washing with PBS. Slides were then incubated in blocking buffer (5% BSA in 0.4% Triton-X100 in PBS (PBST)) for 1 hour at 4°C. Primary antibody solution in blocking buffer was then applied overnight at 4°C. Primary antibody information and dilutions are listed in **Tables 1 and 4**. Slides were then washed with RT 0.1% PBST three times and incubated in secondary antibody solution in blocking buffer (1:500) for 1 hour at RT. Slides then were incubated in Hoechst (Invitrogen, catalog no: H3569 diluted 1:1000 in PBS) for 10 min and washed with PBS three times and cover-slipped with Fluoro-Gel (Electron Microscopy Sciences, catalog no: 17985-10).

**Table. 4.** Details of antibodies used for immunofluorescence.

| Antibody (clone) | Species | Company | Catalog no. | IHC | Antigen retrieval | Temperature | Time |
| --- | --- | --- | --- | --- | --- | --- | --- |
| Anti-NeuN (A60) | Mouse mono | Millipore Sigma | MAB377 | 1:1000 | 20 m | 4°C | o/n |
| Anti-NeuN | Guinea pig poly | Millipore Sigma | ABN90 | 1:1000 | 20 m | 4°C | o/n |
| Anti-Calbindin D-28k | Rabbit | Swant Inc | CD38 | 1:1000 | 20 m | 4°C | o/n |
| Anti-Parvalbumin | Guinea pig poly | Synaptic Systems | 195 044 | 1:1000 | 20 m | 4°C | o/n |
| Anti-RFP | Chicken poly | Rockland Immunochemicals | 600-901-379 | 1:1000 (no antigen retrieval)<br>1:500 (with antigen retrieval) | No<br>20 m | 4°C | o/n or 48 h* |
| Anti-MEIS2 (H-10) | Mouse mono | Santa Cruz Biotechnology | sc-515470 | 1:1000 | 20 m | 4°C | o/n or 48 h* |
| Anti-TBR2 | Rabbit poly | Abcam | ab23345 | 1:500 | 20 m | 4°C | o/n |
| Anti-p27 | Mouse | BD Biosciences | 610241 | 1:500 | 20 m | 4°C | o/n |
| Alexa Fluor 555 anti-chicken IgY (H+L) | Goat | Invitrogen | A21437 | 1:500 | n/a | RT | 1 h |
| Alexa Fluor 488 anti-rabbit IgG (H+L) | Donkey | Invitrogen | A21206 | 1:500 | n/a | RT | 1 h |
| Alexa Fluor 647 anti-rabbit IgG (H+L) | Donkey | Invitrogen | A31573 | 1:500 | n/a | RT | 1 h or 2 h* |
| Alexa Fluor 488 anti-mouse IgG (H+L) | Donkey | Invitrogen | A21202 | 1:500 | n/a | RT | 1 h |
| Alexa Fluor 647 anti-mouse IgG (H+L) | Donkey | Invitrogen | A31571 | 1:500 | n/a | RT | 1 h or 2 h* |
| Alexa Fluor 647 anti-guinea pig IgG (H+L) | Goat | Invitrogen | A21450 | 1:500 | n/a | RT | 1 h |
| Anti-mouse IgG antibody (H+L), Biotinylated | Goat | Vector Laboratories | BA-9200 | 1:500 | n/a | RT | 1 h |
| Anti-chicken IgG antibody (H+L), Biotinylated | Goat | Vector Laboratories | BA-9010 | 1:500 | n/a | RT | 1 h |
| *Applicable for free-floating immunofluorescence or immunohistochemistry |  |  |  |  |  |  |  |

For free-floating sections, sections were first washed three times in PBS for 10 min. Sections then were incubated in blocking buffer (10% normal donkey serum in 0.5% PBST, Sigma-Aldrich, catalog no: D9663-10ML) for 1 hour at RT before incubating in primary antibody solution (2% normal donkey serum in 0.4% PBST) for 24-48 hours at 4°C. Slides were washed in PBS three times and incubated in secondary antibody solution (1:500 in 2% normal donkey serum in 0.4% PBST) for 2 hours at RT. Slides then were incubated in Hoechst (1:1000 in PBS) for 10 min and washed with PBS three times. Sections were mounted on glass slides and after sections had fully dried, cover-slipped with Fluoro-Gel (Electron Microscopy Sciences).

### Immunohistochemistry

All steps were performed at RT, unless specified. Cryosectioned tissue mounted on slides were washed three times in PBS for 5 min (hereafter PBS washes). Antigen retrieval was achieved by immersing sections in sodium citrate buffer at 95°C for 40 min and washing in PBS. Slides were then treated with 50% methanol (in deionized H_2_O) with 0.03% H_2_O_2_ for 1 hour and washed with PBS. Primary antibody solution in blocking buffer (5% BSA in 0.4% PBST) was applied overnight at 4°C. After washing in PBS, a biotinylated secondary antibody in blocking buffer (1:500) was applied for 1 hour, followed by PBS washes. Vectastain Elite ABC HRP solution (Vector Labs, Burlingame, CA, USA; PK-6100) in blocking buffer (1:500 for A and B) was applied for 1 hour, followed by PBS washes. Sections were washed in 0.175 M sodium acetate (in ddH_2_O) and incubated in either standard DAB (0.02% 3,3’-Diaminobenzidine tetrahydrochloride (Sigma-Aldrich, D5905), 2.5% H_2_O_2_ in 0.175 M sodium acetate) or Nickel-enhanced DAB (Ni-DAB) solution (addition of 2.5% NiSO_4_ in standard DAB solution) for 20 min. DAB or Ni-DAB reaction was stopped by washing sections in 0.175 sodium acetated followed by PBS washes. Sections were mounted on glass slides and dried overnight. After dehydration and defatting (dH_2_O-70% ethanol-95%–95%-100%-100%-xylene-xylene-xylene, 1 min each) slides were mounted with a coverslip using DPX mounting medium (Electron Microscopy Sciences).

### Single molecule fluorescence *in situ* hybridization

Tissue was treated according to the RNAscope Multiplex Fluorescent Assay v2 manufacturer’s instructions and reagents from Advanced Cell Diagnostics (Hayward, CA). Details of reagents are listed in **Table 1**. Sections were rinsed twice (5 min each) in RNAase and DNAase free PBS before mounting onto Superfrost Plus coated slides (VWR, catalog no: 48311-703) in RNase- and DNase-free PBS. Slides were dried overnight and heated for 1 hour at 60°C. Slides were pretreated with target retrieval buffer for 10 min at 95-100°C, then treated with Protease III for 30 min at 40°C, followed by a probe incubation for two hours at 40°C. The probes used were: Mm-*Slc32a1*-C1, Mm-*Slc17a6*-C2 and Mm-*Slc6a5*-C3. After probe incubation, slides underwent three amplification steps, followed by developing horseradish peroxidase signal fluorescence with TSA-based fluorophores. Hoechst 33258 was used as a counterstain and slides were dried for 15 min before cover-slipping with Fluoro-Gel.

### Double labeling by RNA *in situ* hybridization and protein immunofluorescence

Sagittal sections (14 um) were first processed for *in situ* hybridization of *Slc6a5* as described previously^91^. Briefly, probes were *in vitro* transcribed from PCR-amplified templates prepared from cDNA synthesized from an adult cerebellum lysate. The forward (5’-GTATCCCACGAGATGGATTGTT-3’) and reverse (5’-CCATACAGAACACCCTCACTCA-3’) primers used for PCR amplification were based on the Allen Brain Atlas. Primers were flanked in the 5’ with SP6 (antisense) and 3’ with T7 (sense) promoters. After visualizing the probe, slides were incubated in 4% PFA overnight at 4°C. Following PBS washes, slides were incubated in sodium citrate buffer (10 mM sodium citrate with 0.05% Tween-20, pH 6.0) at 95°C for 40 min. Slides were placed in blocking buffer for one hour and then incubated in primary antibody solution (anti-RFP at 1:500 in blocking buffer) overnight at 4°C. Slides were washed with 0.1% PBST three times and incubated in secondary antibody solution in blocking buffer (1:500) for 1 hour at RT. Slides were then incubated in Hoechst (1:1000 in PBS, Invitrogen, catalog no: H3569) for 10 min and washed with PBS three times and cover-slipped with Fluoromount-G (ThermoFisher Scientific, catalog no: 00-4958-02).

### Microscopy, image processing, and analyses

All images were acquired with a DM6000 Leica fluorescent microscope (Leica Camera, Wetzlar, Germany) or NanoZoomer Digital Pathology (Hamamatsu Photonics, Shizuoka, Japan). Images were processed or analyzed using Fiji^92^ or Photoshop (Adobe Inc., San Jose, CA, USA). Cell counts for littermate controls, *SepW1-En1/2* CKO, *Atoh1-En1/2* CKO; *Ai75D*, *Atoh1-Cre; Ai75D*, and *eCN-DTA* mice were manually obtained using the Cell Counter plugin for Fiji or semi-automated using a custom script using the Analyze Particle plugin for Fiji as previously described^18^.

The CN subregions were delineated for quantifying the recombination efficiency in *SepW1-Cre; Ai75D* mice using MEIS2 staining and the mouse brain atlas^93^. 10-14 brain slices that were 40-um apart were analyzed. The percentage of tdTomato+ cells across the CN were calculated for each animal (total n=4 mice) and averaged. The same slides were used to calculate the percentage of tdTomato+ cells in the medial CN that were co-labeled for MEIS2 (total n=4 mice) and averaged.

### Brain sample preparation for *ex vivo* dMRI

Mice were transcardially perfused with RT PBS containing heparin and then 4°C 4% PFA. Brains were kept inside the skull, but skin, eyeballs and muscle surrounding was removed and then kept overnight in 4% PFA at 4°C. Samples were then stored in PBS with 0.02% sodium azide until imaging. One week before imaging, samples were equilibrated with Gadodiamide (0.2 mM) in PBS at 4°C and scanned by an experimenter blind to the sex and genotype.

### High resolution *ex vivo* dMRI

Imaging of the brains was conducted as previously described^39,40^. High resolution *ex vivo* dMRI datasets were acquired on a horizontal 7T MR scanner (Bruker Biospin, Billerica, MA, USA) using a 72-mm conventional circularly polarized birdcage radiofrequency resonator (Bruker Biospin, Ettlingen, Germany) for homogeneous transmission in conjunction with a four-channel receive-only phased array CryoProbe (CRP, Bruker Biospin, Ettlingen, Germany) and a modified 3D diffusion-weighted gradient-and-spin-echo (DW-GRASE) sequence^94^ (TE/TR: 35 ms/ 400 ms, b-value: 5000 s/mm^2^, no. of diffusion directions: 60, resolution: 0.1 mm^3^ isotropic, acquisition time: 10.6h).

### dMRI data processing

dMRI rawdata was processed using a pipeline demonstrated in our earlier studies^39,40^. In brief, the following steps were followed for each subject: 1) removal of signals from non-brain tissue in the dMRI datasets using AMIRA (ThermoFisher Scientific, https://www.thermofisher.com), 2) Correction for specimen displacements using rigid alignment implemented in DTIStudio^95^, 3) calculation of diffusion tensor, fractional anisotropy, and fiber orientation and distribution maps Using MRtrix^96^ (https://www.mrtrix.org/), 4) regional volumetric analysis, and 5) streamline tractography between the selected pairs of brain regions or nodes.

### Volumetric analysis

For regional brain volumetric analysis, each subject was mapped to a high-resolution mouse brain atlas^39^. Whole brain volume or 10 regional volumes (cortex – CTX, olfactory area – OLF, hippocampus – HPF, amygdala – AMY, striatum – STR, pallidum – PAL, thalamus – TH, hypothalamus – HY, midbrain – MB, and cerebellum – CB) were calculated.

### Streamline tractography and structural connectome generation

Structural connectivity between nodes and the structural connectome were constructed as previously reported^39,40,97^. In short, eight nodes (anterior cingulate cortex – ACA, motor cortex – MO, somatosensory cortex – SS, HPF, STR, AMY, TH, and HY) were included to construct the node-to-node SC maps using streamline tractography as implemented in MRtrix by seeding (n = 50,000) within the seed region with minimum streamline length of 3 mm, FOD cut-off 0.05, step size 0.025, and angle 45°. Excluding the intra-regional connectivity, in total 28 connections per subject were generated, which were subsequently used to construct the subject-specific structural connectome^40^.

### Global network analysis

Brain global network properties, global efficiency (Geff) and small-worldness (SW), were evaluated as previously described^97^ using principles of graph theory^98^ via GRETNA software^99^. Geff represents the efficiency of distant information transfer in a network and was defined as the inverse of the average characteristic path length between all nodes in the network. Brain network with short average path length between nodes and high degree of interconnectedness in local networks is considered to have SW properties. SW was computed by normalizing the network with respect to 1000 simulated random networks with equal distribution of edge weight and node strength as reported previously^100^.

### Statistics

No statistical methods were used to predetermine sample sizes, but our sample sizes are similar to those reported previously^18,33,59^. At least three different litters were used for all experiments (cell counting and behavior). No litter-related effects were seen in behavioral studies. Experimenters were blind to mouse genotypes during behavior analysis and quantifications. All data were subject to the Shapiro-Wilk test and Q-Q plots were generated to evaluate normality. If one or more dependent variable(s) failed the normality test, a non-parametric test was used. For behavioral experiments, outliers were defined if at least one dependent variable was more than mean+2×SD or less than mean-2×SD. Some additional criteria for outliers are indicated for each behavioral test (see above). A given outlier was not consistent across experiments. All statistical analyses were carried out with GraphPad Prism 8 software. Comparisons between two groups were analyzed using parametric test (unpaired *t*-test) or non-parametric test (Mann-Whitney test). The Holm-Šídák method was used for multiple unpaired t-test comparisons. Wilcoxon signed-rank test (null hypothesis = 50%) was used to test normal social preference in the three-chambered social approach test. Ordinary two-way ANOVA was used to compare means across two or more dependent variables. Repeated measures two-way ANOVA with Šídák’s multiple comparisons test were used for basal locomotion, accelerating rotarod, and water Y-maze. A three-way ANOVA was used to examine the effects of sex and genotype on behavior, but as there were no sex differences (data not shown), final analyses included both males and females. Data are presented as mean ± SEM or mean ± SD as indicated in each figure legend. Mean differences were considered significant if P < 0.05.

## Acknowledgements

We thank all members of the Joyner, Rajadhyaksha, Heck, and Zhang laboratories over the past five years for insightful discussions and technical support. We thank Alexander Walsh, Narmin Mekawy, Diego Scala Chavez, and Dr. Arlene Martinez-Rivera for their excellent support from the Rajadhyaksha lab. We thank Drs. Kristen Pleil (Weill Cornell Medicine), Peter Tsai (UT Southwestern), and Jacqueline Crawley (UC Davis) for their advice and input on data interpretation and technical support for DREADDs and behavioral testing. We thank Dr. Choong Heon Lee for his help with diffusion MRI scanning and post-processing. This work was supported by grants from the NIH to Alexandra L. Joyner (R37MH085726 and R01NS092096), Andrew S. Lee (NICHD T32HD060600), Anjali M. Rajadhyaksha (R01MH118934, R01MH125006, R01DA029122, R01DA050454), Natalia V. De Marco Garcia (2R01MH110553, 1R01MH125006, 1R01NS116137), Detlef H. Heck (R01MH112143), and Jiangyang Zhang (R01NS102904). Alexandra L. Joyner is also supported by an NCI Cancer Center Support Grant (CCSG, P30 CA08748) and the Cycle for Survival. Andrew S. Lee was supported by a Weill Cornell Medicine Clinical & Translational Science Center Predoctoral Training Award (TL1TR002386) from the National Center for Advancing Translational Sciences. Alina Gubanova was supported by a Fordham College at Lincoln Center Dean’s Undergraduate Research and Creative Practice Grant. Anjana Krishnamurthy was supported by a Dorris J. Hutchison Predoctoral Fellowship. Anjali M Rajadhyaksha was supported by the Weill Cornell Autism Research Program.

## Contributions

A.S.L., A.M.R., T.M.A., D.H.H., J.Z., A.L.J. formulated experiments and analysis. A.S.L., A.G., D.N.S., Y.L., Z.L., A.K. performed experiments and anlaysis. T.M.A. and J.Z. carried out the mouse diffusion MRI experiments and anlaysis. N.V.D.M.G. provided the *SepW1-Cre* mice. A.S.L., T.M.A., A.M.R., and A.L.J. prepared the manuscript.

## Data availability

Any additional information and requests for reagents and resources should be directed to and will be satisfied by the lead contact ALJ.

## Code Availability

All data reported in this study and the analysis codes used are available upon request.

---

**Supplementary Fig. 1.**
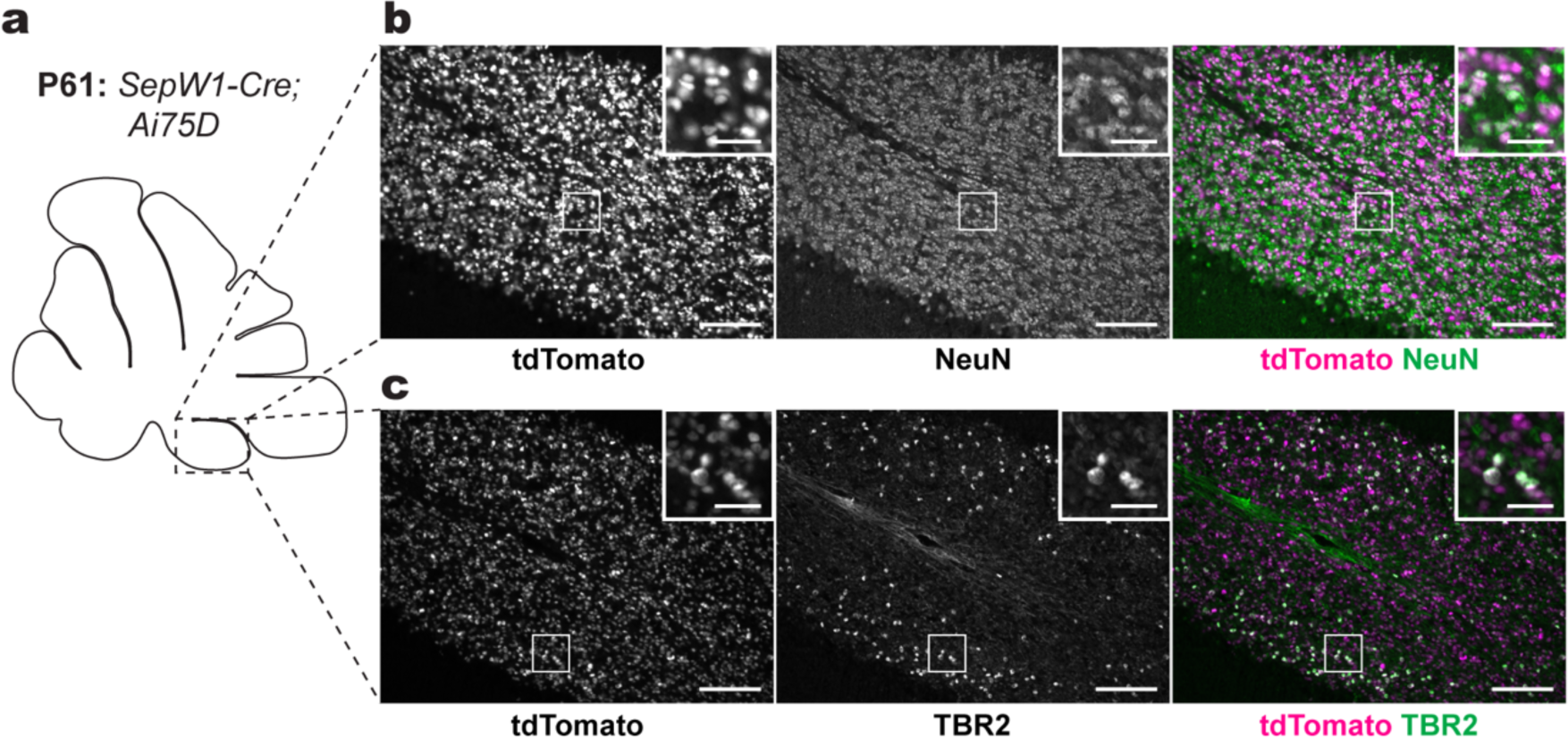
*SepW1-Cre* recombines in granule cells and unipolar brush cells. **a**, Schematic representation of sagittal plane of an adult mouse showing where **b** and **c** images were acquired. **b**, Immunofluorescence images of tdTomato (magenta) and NeuN (green) co-expressing granule cells in adult *SepW1-Cre; Ai75D* mice. Scale bars = 100 um; inset scale bars = 25 um. **c,** Immunofluorescence images of tdTomato (magenta) and TBR2 (green) co-expressing unipolar brush cells in adult *SepW1-Cre; Ai75D* mice. Scale bars = 100 um.

**Supplementary Fig. 2.**
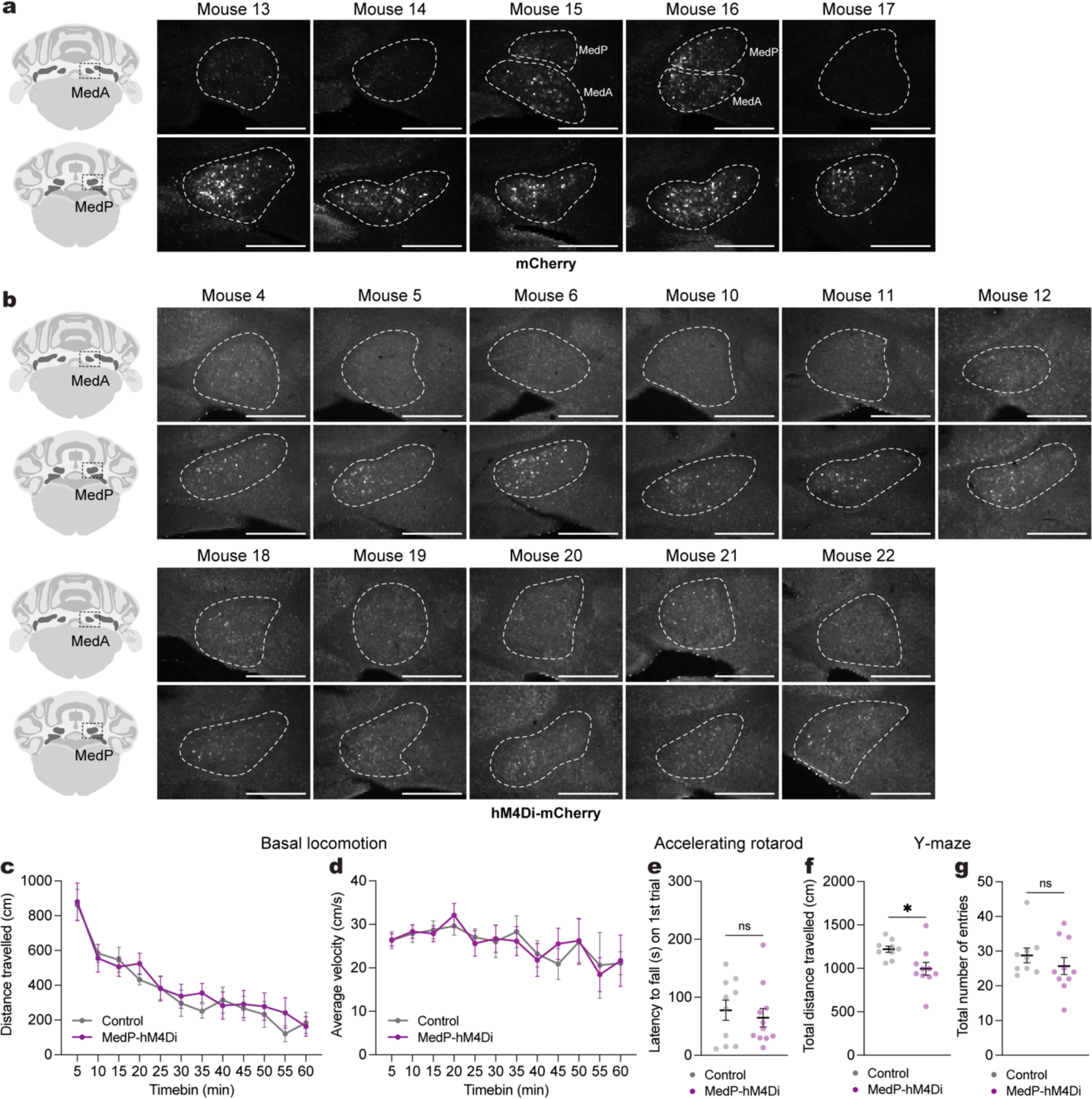
Motor coordination during behavior is largely intact after acute chemogenetic inhibition of adult MedP eCN. **a**, Schematic (left) and representative images of mCherry expression (right) in MedA (upper row) and MedP (lower row) CN in the five control mice. The other six control mice were Cre-negative (treated with CNO). Scale bars = 500 um. **b**, Schematic (left) and representative images of hM4Di-mCherry expression (right) in MedA and MedP CN in the eleven MedP-hM4Di mice. Scale bars = 500 um. **c**, Distance travelled during basal locomotion by 5 min time bins (n=11 per group). Repeated measure two-way ANOVA: no main effect of time (P = 0.0609), chemogenetics (P = 0.9730) or interaction (P = 0.9975). **d**, Average velocity during basal locomotion by 5 min time bins (n=11 per group). Repeated measure two-way ANOVA: main effect of time (F_4.870,97.41_ = 28.43, P < 0.0001), but not of chemogenetics (P = 0.7000) or interaction (P = 0.8668). **e**, Latency to fall on the first trial of the accelerating rotarod test (MedP-hM4Di: n=11, control: n=10; Mann Whitney *U* test: *U* = 48, P = 0.6412). **f**, Total distance travelled in the Y-maze (MedP-hM4Di: n=10, control: n=9; two-tailed unpaired t-test: t_17_ = 2.648, P = 0.0169). **g**, Total number of arm entries in the Y-maze (MedP-hM4Di: n=10, control: n=9; Mann Whitney *U* test: *U* = 30, P = 0.2326). ns, not significant: P ≥ 0.05. Data are presented as mean values ± SEM.

**Supplementary Fig. 3.**
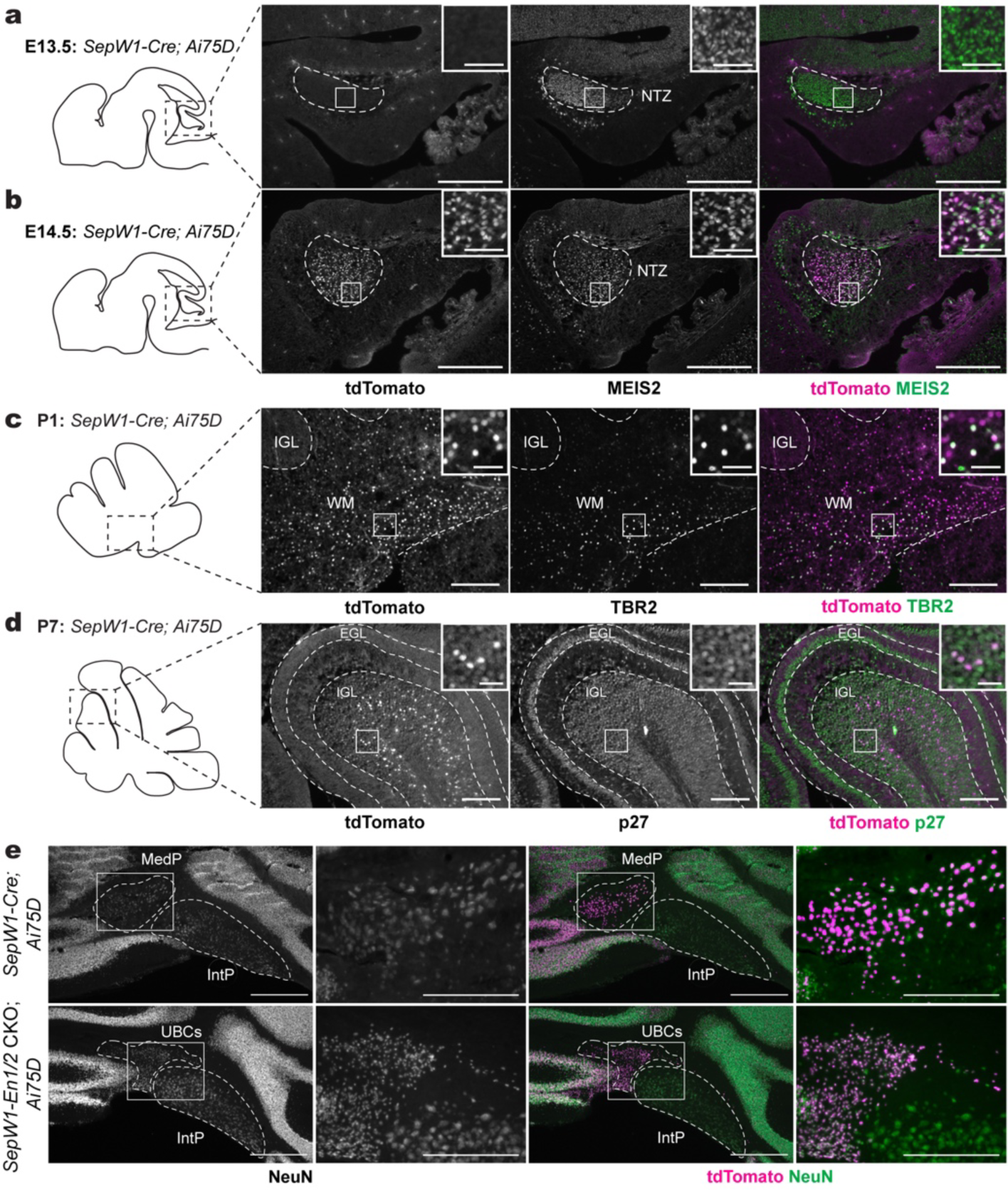
*SepW1-Cre* recombines in the developing excitatory cerebellar neurons and *SepW1-En1/2* CKOs have preferential loss of MedP CN. **a,b,** Schematic (left) and representative images (right) of sagittal sections stained for tdTomato (magenta) in *SepW1-Cre; Ai75D* mice showing recombination at E14.5 (**b**) but not at E13.5 (**a**) in eCN in the NTZ marked by MEIS2 (green). NTZ = nuclear transitory zone. **c**, Schematic (left) and representative sagittal image (right) of tdTomato (magenta) expression in *SepW1-Cre; Ai75D* mice showing recombination at postnatal day (P1) in TBR2+ (green) unipolar brush cells. **d**, Schematic (left) and representative sagittal images (right) of tdTomato (magenta) expression in *SepW1-Cre; Ai75D* mice showing recombination at P7 in p27+ (green) differentiated granule cells in the internal granule cell layer (IGL), but not in proliferating granule cell precursors in the external granule cell layer (EGL). Scale bars = 100 um; inset scale bars = 20 um. **e**, Representative images of coronal sections stained for NeuN (single channel), NeuN (green) and tdTomato (magenta) co-labeling in the posterior CN of *SepW1-Cre; Ai75D* and *SepW1-En1/2* CKO; *Ai75D* mice (*SepW1-Cre/+; En1^flox/flox^; En2^flox/flox^; R26^LSL-nls-tdTomato/+^*). NeuN labeling near the MedP of mutants are ectopic unipolar brush cells that are not TBR2+ or MEIS2+ (confirmed in Krishnamurthy et al., 2024). Abbreviations: MedP=Posterior medial; IntP=Posterior interposed. Scale bars for low magnification = 500 um; scale bars for high magnification = 100 um. Scale bars in **a**, **b**, **c** = 250 um; inset scale bars = 50 um.

**Supplementary Fig. 4.**
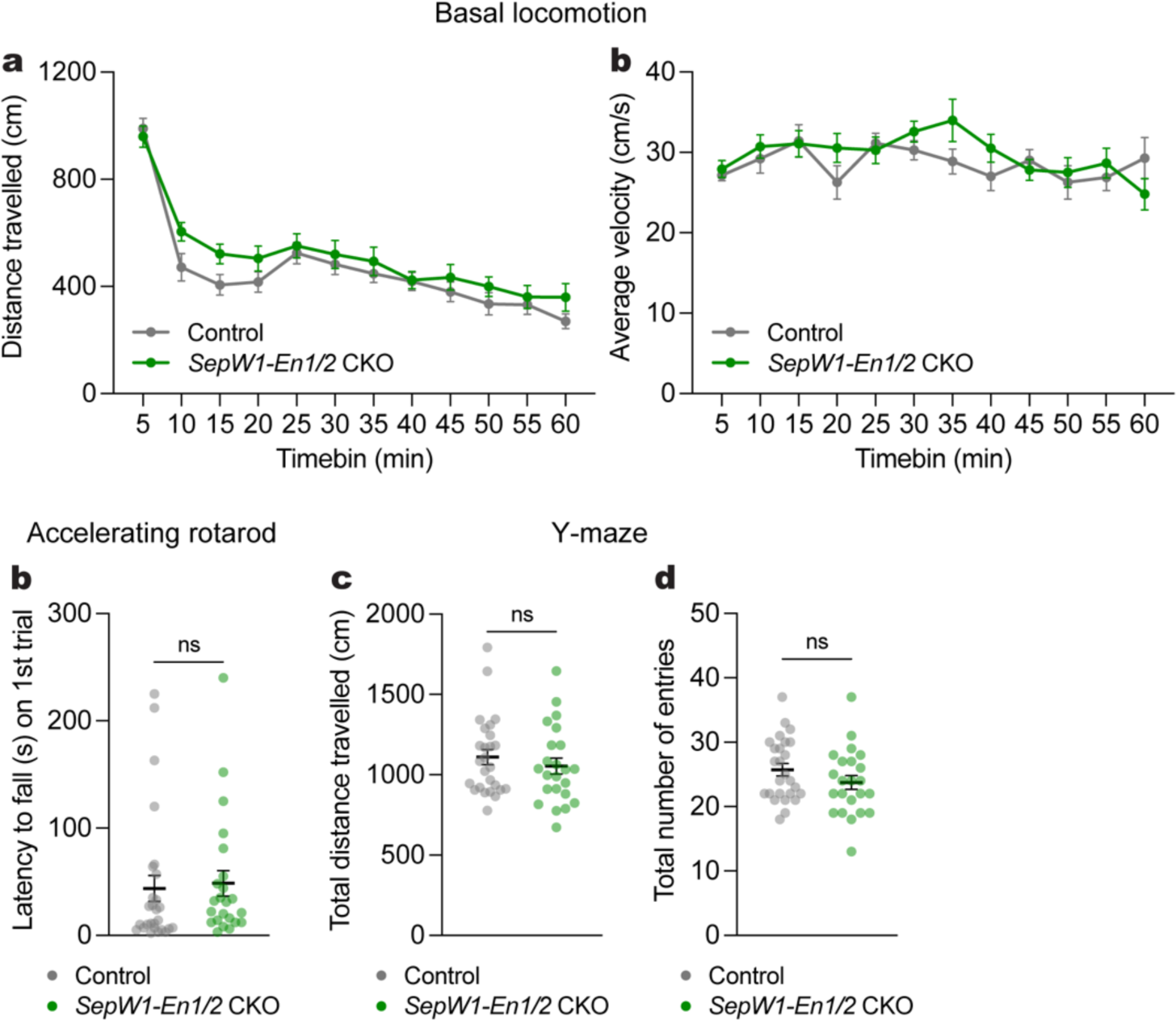
Motor coordination during behaviors is not altered in *SepW1-En1/2* CKOs. **a**, Distance travelled during basal locomotion by 5 min time bins (*SepW1-En1/2* CKOs: n=24, littermate controls: n=27). Repeated measure two-way ANOVA: main effect of time (F_5.410,265.1_ = 71.20, P < 0.0001), but not of genotype (P = 0.2081) or interaction (P = 0.2042). **b**, Average velocity during basal locomotion by 5 min time bins (*SepW1-En1/2* CKOs: n=24, littermate controls: n=27). Repeated measure two-way ANOVA: main effect of time (F_7.996,390.3_ = 2.386, P = 0.0162), but not of genotype (P = 0.3429) or interaction (P = 0.1806). **c**, Latency to fall on the first trial of the accelerating rotarod test (*SepW1-En1/2* CKOs: n=23, littermate controls: n=27; Mann-Whitney *U* test: *U* = 240.5, P = 0.1759). **d**, Total distance travelled in the Y-maze (*SepW1-En1/2* CKOs: n=23, littermate controls: n=26; Mann-Whitney *U* test: *U* = 266, P = 0.5184). **e**, Total number of arm entries in the Y-maze (*SepW1-En1/2* CKOs: n=23, littermate controls: n=26; two-tailed unpaired t-test: t_47_ = 1.397, P = 0.1689). ns, not significant: P ≥ 0.05. Data are presented as mean values ± SEM.

**Supplementary Fig. 5.**
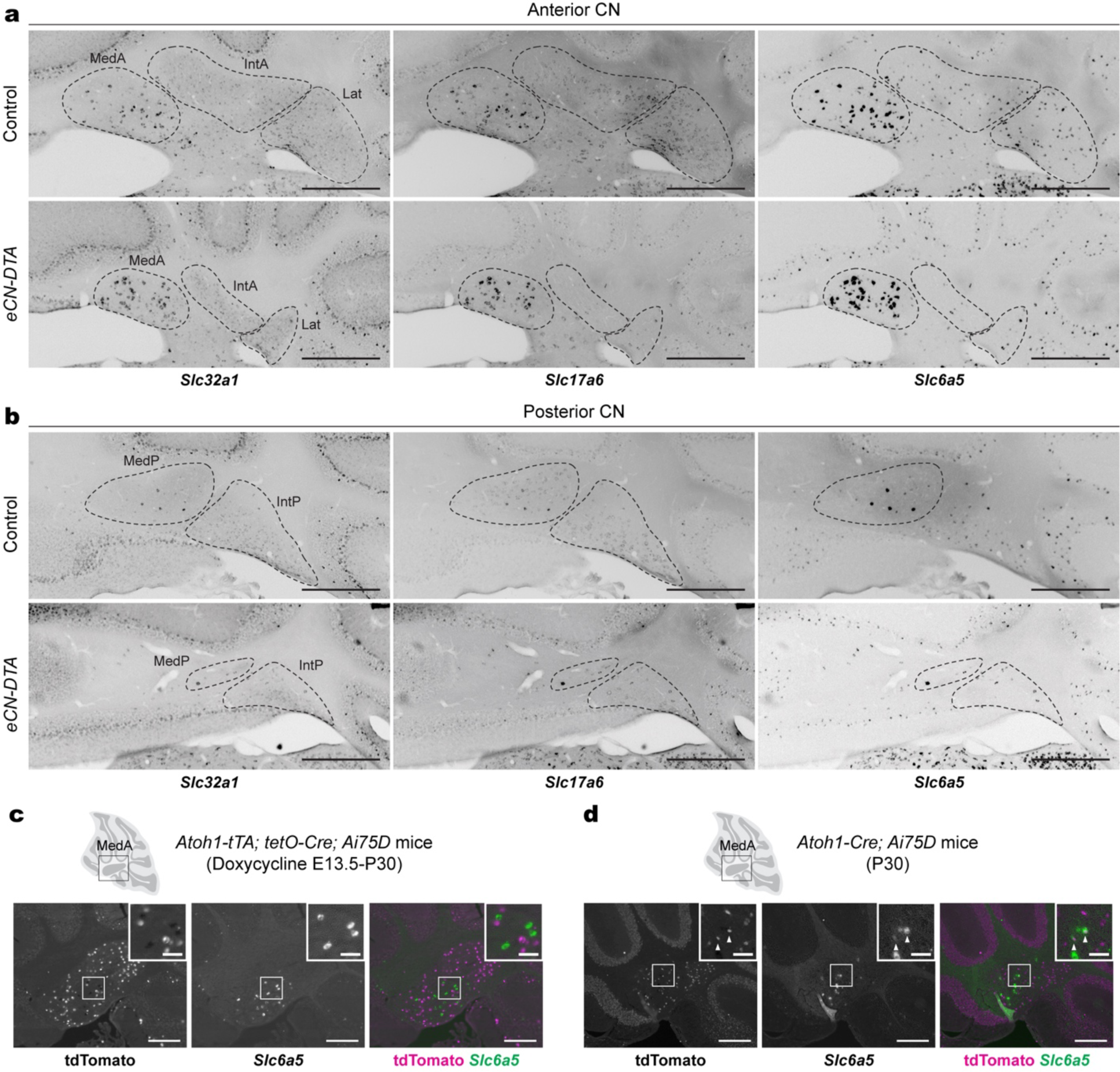
Remaining CN neurons in *eCN-DTA* mice are inhibitory neurons. **a,b**, Representative images of coronal sections of triple RNA *in situ* analysis of *Slc32a1*, *Slc17a6* and *Slc6a5* with single channel expression in anterior (**a**) and posterior (**b**) CN of *eCN-DTA* mice and littermate controls. Dotted outlines indicate the CN subregions. Abbreviations: MedA=Anterior medial; MedP=Posterior medial; IntA=Anterior interposed; IntP=Posterior interposed; Lat=Lateral. Scale bars = 500 um. **c**, Representative images from the MedA region of double RNA *in situ* hybridization and immunofluorescence for *Slc6a5* and tdTomato in P30 *Atoh1-tTA; tetO-Cre; Ai75D* (*Atoh1-tTA/+; tetO-Cre; R26^LSL-nls-tdTomato/+^*) mice treated with doxycycline from E13.5 until P30. *Slc6a5*+ CN neurons are not labeled by the *Atoh1-tTA* transgene (tdTomato as a readout). Scale bars = 250 um; inset scale bars = 50 um. **d**, Representative images from the MedA region of double RNA *in situ* hybridization and immunofluorescence for *Slc6a5* and tdTomato in P30 *Atoh1-Cre; Ai75D* mice. Subset of *Slc6a5*+ CN neurons are labeled by the *Atoh1-Cre* transgene (tdTomato as a readout). Scale bars = 250 um; inset scale bars = 50 um.

**Supplemental Fig. 6.**
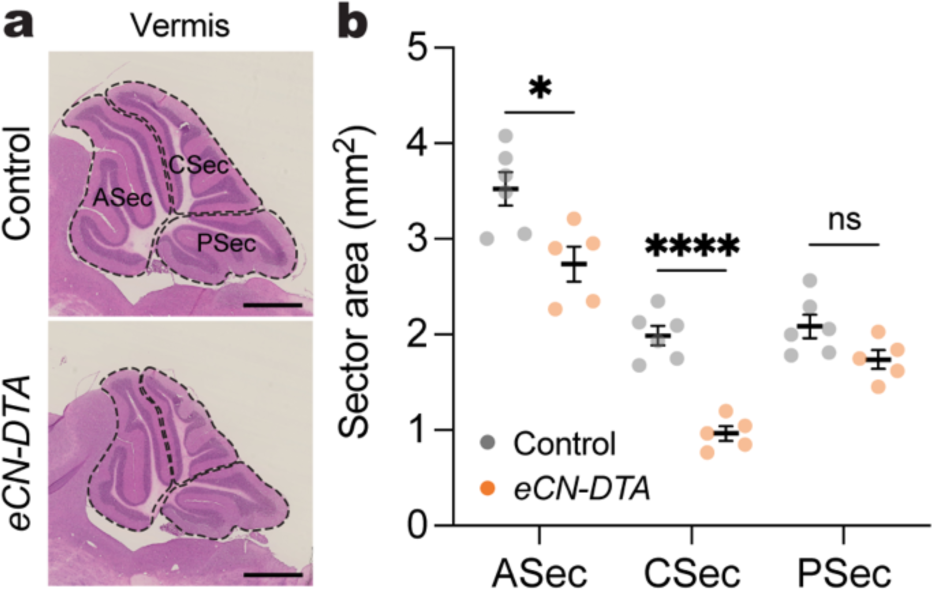
*eCN-DTA* mice show reduced growth in the anterior and central vermis. **a**, Representative images of H&E labeled vermis in *eCN-DTA* and littermate control mice. Anterior, central and posterior sectors (ASec, CSec, and PSec, respectively) are outlined in dotted lines. Scale bars = 1 mm. **b**, Quantification of sector area in eCN-DTA mice (n=5) compared to littermate controls (n=6). Ordinary two-way ANOVA: main effect of genotype (F_1,9_ = 17.96, P = 0.0022), main effect of sector (F_1.693,15.23_ = 264.6, P < 0.0001), and interaction (F_2,18_ = 10.65, P = 0.0009); with post hoc two-tailed t-tests with uncorrected Fisher’s LSD for effect of genotype for ASec (t_8.779_ = 3.100, P = 0.0131), CSec (t_8.706_ = 8.001, P < 0.0001), and PSec (P = 0.0540). ns, not significant: P ≥ 0.05. Data are presented as mean ± SEM.

**Supplementary Fig. 7.**
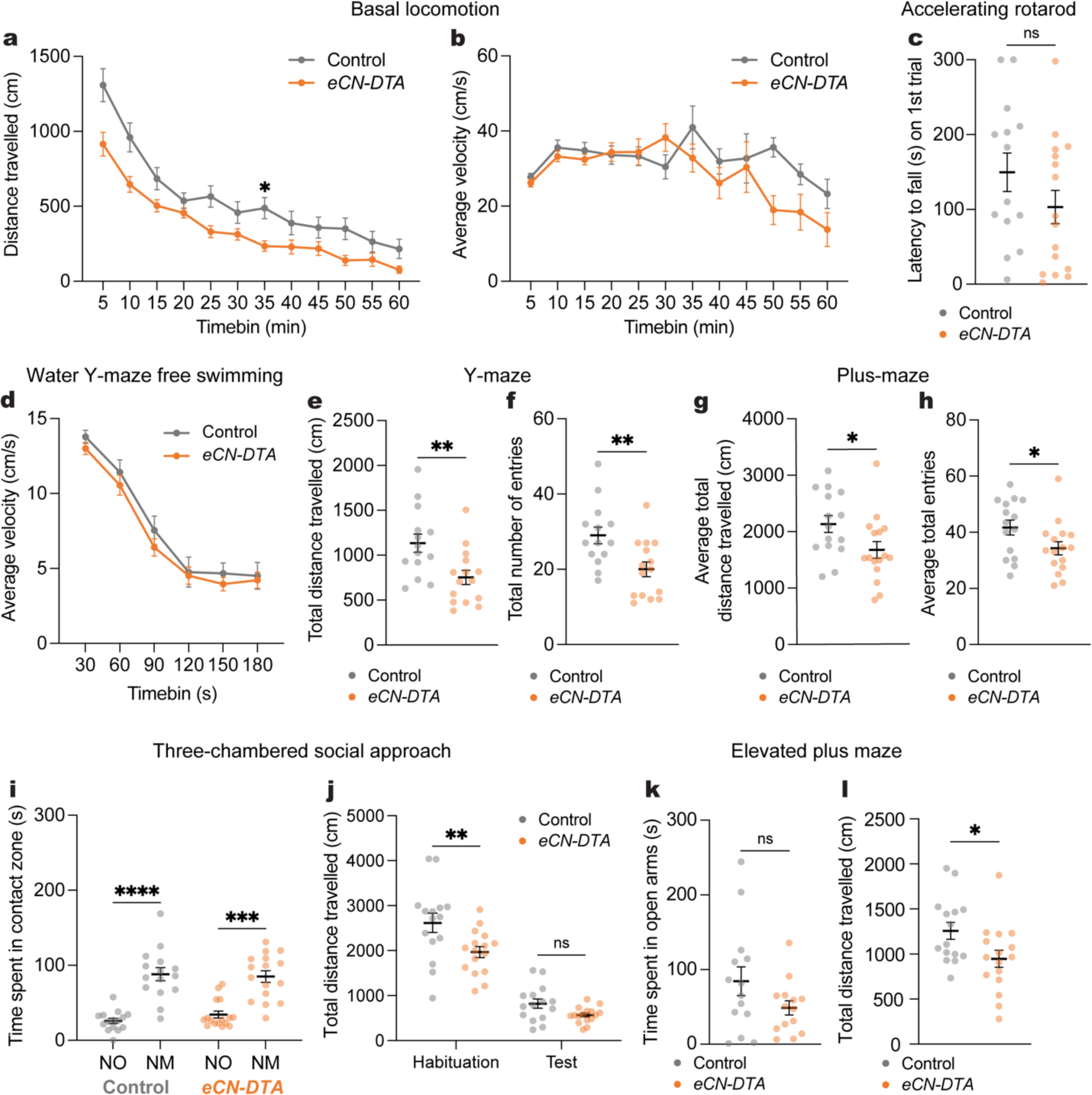
Motor coordination during behavior is task-dependent in *eCN-DTA* mice. **a**, Distance travelled during basal locomotion by 5 min time bins (*eCN-DTA* mice: n=16, littermate controls: n=15). Repeated measure two-way ANOVA: main effect of time (F_5.410,156.9_ = 100.4, P < 0.0001), main effect of genotype (F_1,29_ = 8.210, P = 0.0077), and interaction (F_11,319_ = 2.629, P = 0.0032); with post hoc two-tailed t-tests with Šídák correction for effect of genotype for 30-35 min (t_20.48_ = 3.245, P = 0.0466) but no other comparisons (P ≥ 0.05). **b**, Average velocity during basal locomotion by 5 min time bins (*eCN-DTA* mice: n=16, littermate controls: n=15). Repeated measure two-way ANOVA: main effect of time (F_5.283,153.2_ = 5.057, P = 0.0002), but not of genotype (P = 0.0883) or interaction (P = 0.0589). **c**, Latency to fall on the first trial of the accelerating rotarod test (*eCN-DTA* mice: n=16, littermate controls: n=14; Mann-Whitney *U* test: *U* = 72.50, P = 0.1034). **d**, Average swimming velocity during a three-minute swim (*eCN-DTA* mice: n=16, littermate controls: n=15). Repeated measure two-way ANOVA: main effect of time (F_5,145_ = 137.8, P < 0.0001), but not of genotype (P = 0.3829) or interaction (P = 0.9235). **e**, Total distance travelled during the Y-maze test (*eCN-DTA* mice: n=15, littermate controls: n=14; two-tailed unpaired t-test: t_27_ = 3.027, P = 0.0054). **f**, Total number of arm entries in the Y-maze (*eCN-DTA* mice: n=15, littermate controls: n=14; two-tailed unpaired t-test: t_27_ = 2.969, P = 0.0062). **g,** Average total distance travelled during two days of testing in the plus-maze (*eCN-DTA* mice: n=16, littermate controls: n=15: two-tailed unpaired t-test: t_29_ = 2.211, P = 0.0350). **h,** Average total number of entries during two days of testing in the plus-maze (*eCN-DTA* mice: n=16, littermate controls: n=15: two-tailed unpaired t-test: t_29_ = 2.114, P = 0.0432). **i,** Total time in which the animal’s nose was within the contact zone of a novel mouse (NM) and novel object (NO) during the three-chambered social approach test (*eCN-DTA* mice: n=16, littermate controls: n=15). Repeated measure two-way ANOVA: main effect of location (F_1,29_ = 53.64, P < 0.0001), but not of genotype (P = 0.5828) or interaction (P=0.4639); with post hoc two-tailed t-tests with Šídák correction for effect of location for littermate controls (t_29_ = 5.614, P < 0.0001) and *eCN-DTA* mice (t_29_ = 4.731, P = 0.0001) mice. **j**, Total distance travelled during habituation and test phases of the three-chambered social approach test (*eCN-DTA* mice: n=16, littermate controls: n=15). Repeated measure two-way ANOVA: main effect of phase (F_1,29_ = 358.9, P < 0.0001), genotype (F_1,29_ = 7.130, P = 0.0123), and interaction (F_1,29_ = 5.461, P = 0.0266); with post hoc two-tailed t-tests with Šídák correction for effect of genotype for the habituation phase (t_58_ = 3.433, P = 0.0022) but not test phase (t_58_ = 1.344, P = 0.3343). **k**, Total time spent in the open arms of an elevated plus maze (*eCN-DTA* mice: n=14, littermate controls: n=14; two-tailed unpaired t-test: t_26_ = 1.662, P = 0.1086). **l**, Total distance travelled in an elevated plus maze (*eCN-DTA* mice: n=15, littermate controls: n=16; two-tailed unpaired t-test: t_29_ = 2.320, P = 0.0276). ns, not significant: P ≥ 0.05. Data are presented as mean values ± SEM.

**Supplementary Fig. 8.**
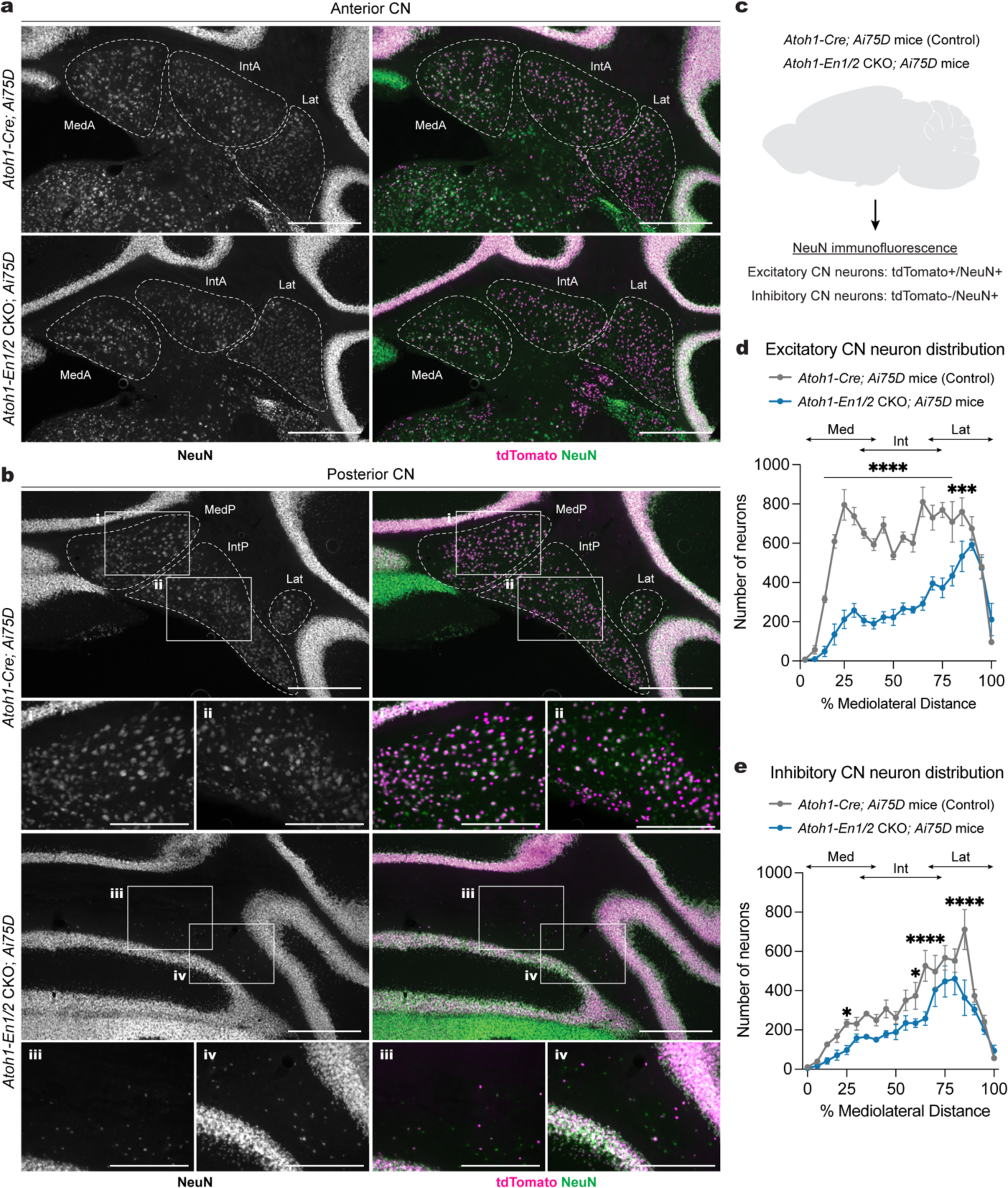
Inhibitory CN neurons are significantly reduced across the mediolateral axis in *Atoh1-En1/2* CKOs. **a**,**b**, Representative images of immunofluorescent staining NeuN (singe channel) and NeuN (green), tdTomato (magenta) co-labeling in of coronal sections of the anterior (**a**) and posterior (**b**) CN of *Atoh1-Cre; Ai75D* and *Atoh1-En1/2* CKO; *Ai75D* mice. Abbreviations: MedA=Anterior medial; MedP=Posterior medial; IntA=Anterior interposed; IntP=Posterior interposed; Lat=Lateral. Scale bars = 500 um, scale bars for **i**-**iv** = 100 um. **c**, Experimental design for quantifying excitatory and inhibitory CN neurons in *Atoh1-Cre; Ai75D* (*Atoh1-Cre/+; R26^LSL-nls-tdTomato/+^*) and *Atoh1-En1/2* CKO; *Ai75D* (*Atoh1-Cre/+; En1^flox/flox^; En2^flox/flox^; R26^LSL-nls-tdTomato/+^*) mice. **d**, Quantification and distribution of excitatory CN neurons in half of the cerebellum (every second section). Two-way ANOVA: main effect of % mediolateral distance (F_19,120_ = 31.38, P < 0.0001), genotype (F_1,120_ = 400.1, P < 0.0001), and interaction (F_19,120_ = 8.830, P < 0.0001); with post hoc two-tailed t-tests with uncorrected Fisher’s LSD for effect of genotype for bin 10-80% (list of t value for each bin: t_120_ = 4.029, 7.169, 8.828, 7.267, 6.726, 6.087, 7.103, 4.759, 5.516, 5.122, 7.85, 5.06, 6, 4.169; all P values: P < 0.0001), for bins 80-85% (t_120_ = 3.435, P = 0.0008), and no other comparisons (P ≥ 0.05). Abbreviations: Med=medial; Int=interposed; Lat=lateral. **e**, Quantification and distribution of inhibitory CN neurons in half of the cerebellum (every second section). Two-way ANOVA: main effect of % mediolateral distance (F_19,120_ = 23.97, P < 0.0001), and genotype (F_1,120_ = 50.81, P < 0.0001), but not interaction (F_19,120_ = 1.659, P = 0.0531); with post hoc two-tailed t-tests with uncorrected Fisher’s LSD for effect of genotype for bin 20-25% (t_120_ = 2.094, P = 0.0384), bin 55-60% (t_120_ = 2.141, P = 0.0343), bin 60-65% (t_120_ = 4.114, P < 0.0001), bin 80-85% (t_120_ = 5.316, P <0.0001), and no other comparisons (P ≥ 0.05). Abbreviations: Med=medial; Int=interposed; Lat=lateral. Data are presented as mean values ± SEM.

**Supplementary Fig. 9.**
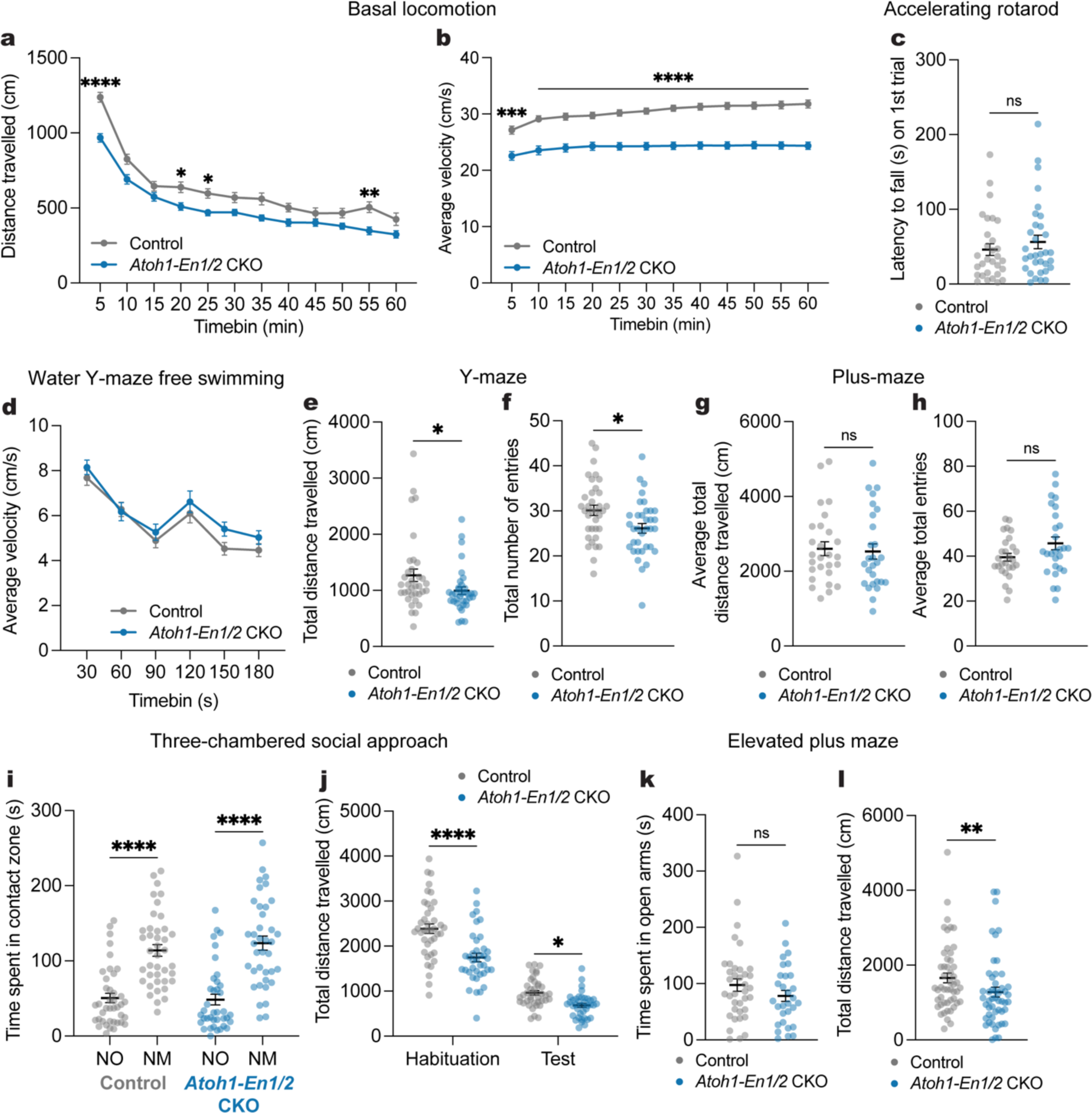
Motor coordination during behavior is task-dependent in *Atoh1-En1/2* CKOs. **a**, Distance travelled during basal locomotion by 5 min time bins (*Atoh1-En1/2* CKOs: n=33, littermate controls: n=35). Repeated measure two-way ANOVA: main effect of time (F_7.310,482.5_ = 171.0, P < 0.0001), genotype (F_1,66_ = 15.45, P = 0.0002), and interaction (F_11,726_ = 3.120, P = 0.0004); with post hoc two-tailed t-tests with Šídák correction for effect of genotype on 0-5 min (t_65.21_ = 6.250, P < 0.0001), 15-20 min (t_61.20_ = 3.012, P = 0.0443), 20-25 min (t_59.87_ = 3.358, P = 0.0163), 50-55 min (t_60.88_ = 3.556, P = 0.0088), and no other comparisons (P ≥ 0.05). **b**, Average velocity during basal locomotion by 5 min time bins (*Atoh1-En1/2* CKOs: n=33, littermate controls: n=35). Repeated measure two-way ANOVA: main effect of time (F_2.435,160.7_ = 171.0, P < 0.0001), genotype (F_1,66_ = 57.43, P = 0.0002), and interaction (F_11,726_ = 4.381, P < 0.0001); with post hoc two-tailed t-tests with Šídák correction for effect of genotype on 0-5 min (t_65.45_ = 4.355, P = 0.0006) and 5-60 min (t value for each bin: t_65.45_ = 4.355, t_61_ = 5.693, t_63.58_ = 5.927, t_62.8_ = 5.916, t_63.54_ = 6.755, t_61.95_ = 7.332, t_62.74_ = 7.867, t_65.1_ = 7.819, t_65.56_ = 7.863, t_65.83_ = 7.695, t_66_ = 7.698, t_65.53_ = 7.589, all P values: P < 0.0001). **c**, Latency to fall on the first trial of the accelerating rotarod test (*Atoh1-En1/2* CKOs: n=32, littermate controls: n=30; Mann-Whitney *U* test: *U* = 417, P = 0.3792). **d**, Average swimming velocity during a three-minute swim (*Atoh1-En1/2* CKOs: n=35, littermate controls: n=31). Repeated measure two-way ANOVA: main effect of time (F_5,320_ = 0.5563, P < 0.0001), but not of genotype (P = 0.1350) and interaction (P = 0.7335). **e**, Total distance travelled during the Y-maze (n=35 per genotype; Mann-Whitney *U* test: *U* = 405, P = 0.0144). **f**, Total number of arm entries in the Y-maze (n=35 per genotype; two-tailed unpaired t-test: t_68_ = 2.548, P = 0.0131). **g**, Average total distance travelled during two days of testing in the plus-maze (n=27 per genotype; Mann-Whitney *U* test: *U* = 332.5, P = 0.5857). **h**, Average total number of arm entries during two days of testing in the plus-maze (n=27 per genotype; two-tailed unpaired t-test: t_52_ = 1.839, P = 0.0717). **i**, Total time in which the animal’s nose was within the contact zone of a novel mouse (NM) and novel object (NO) during the three-chamber social approach test (*Atoh1-En1/2* CKOs: n=38, littermate controls: n=40). Repeated measure two-way ANOVA: main effect of location (F_1,152_ = 81.64, P < 0.0001), but not of genotype (P = 0.6191) or interaction (P = 0.4360); with post hoc two-tailed t-tests with Šídák correction for effect of location for *Atoh1-En1/2* CKOs (t_152_ = 6.854, P < 0.0001) and littermate controls (t_152_ = 5.913, P < 0.0001). **j**, Total distance travelled during habituation and test phases in the three-chamber social approach test (*Atoh1-En1/2* CKOs: n=38, littermate controls: n=40). Repeated measure two-way ANOVA: main effect of phase (F_1,76_ = 487.6, P < 0.0001), genotype (F_1,76_ = 22.58, P < 0.0001), and interaction (F_1,76_ = 10.31, P = 0.0019); with post hoc two-tailed t-tests with Šídák correction for effect of genotype for the habituation phase (t_152_ = 5.772, P < 0.0001) and test phase (t_152_ = 2.491, P = 0.0275). **k**, Total time spent in the open arms of the elevated plus maze (n=50 per genotype; Mann-Whitney *U* test: *U* = 878, P = 0.01). **l**, Total distance travelled in the elevated plus maze (n=50 per genotype; Mann-Whitney *U* test: *U* = 467, P = 0.2705). ns, not significant: P ≥ 0.05. Data are presented as mean values ± SEM.

**Supplementary Fig. 10.**
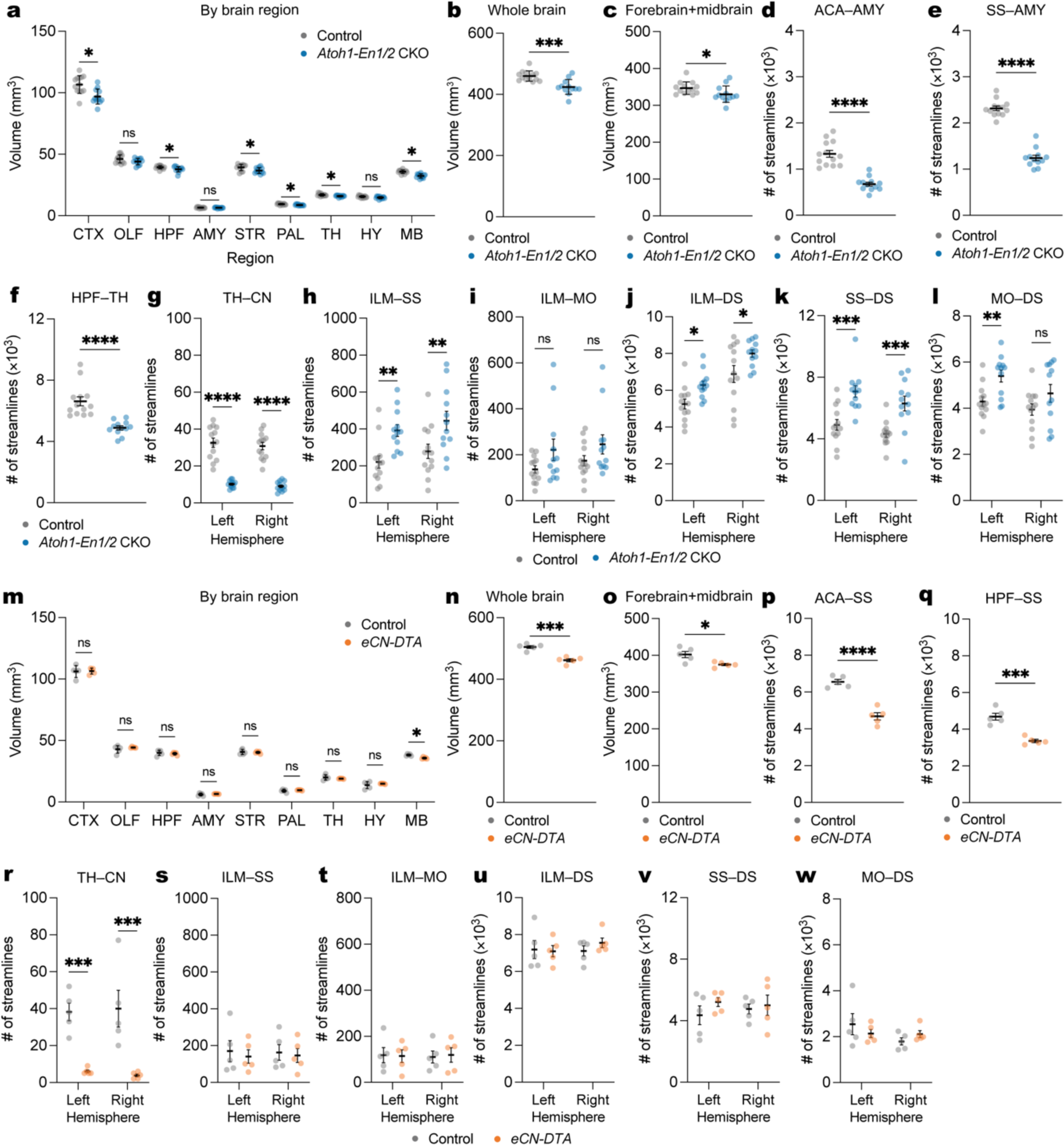
Brain volume and connectivity changes in *Atoh1-En1/2* CKO and eCN-DTA mice. **a**, Quantification of regional brain volumes in *Atoh1-En1/2* CKOs (n=12) compared to littermate controls (n=13). Two-tailed unpaired t-tests to test for effect of genotype on CTX (t_23_ = 3.633, P = 0.0014), HPF (t_23_ = 2.329, P = 0.03), STR (t_23_ = 2.612, P = 0.016), PAL (t_23_ = 3.143, P = 0.005), TH (t_23_ = 3.052, P = 0.006), MB (t_23_ = 5.665, P < 0.0001), HB (t_23_ = 11.16, P < 0.0001), CB (t_23_ = 11.15, P < 0.0001), other comparisons P ≥ 0.05. **b**, Quantification of whole brain volume in *Atoh1-En1/2* CKOs compared to littermate controls (*Atoh1-En1/2* CKOs: n=12, littermate controls: n=13; two-tailed unpaired t-test: t_23_ = 4.298, P = 0.0003). **c**, Quantification of forebrain and midbrain combined volume in *Atoh1-En1/2* CKOs compared to littermate controls (*Atoh1-En1/2* CKOs: n=12, littermate controls: n=13; two-tailed unpaired t-test: t_23_ = 2.087, P = 0.0481). **d**, Quantification of average (left and right hemispheres) ACA-AMY tractography in *Atoh1-En1/2* CKOs compared to littermate controls (*Atoh1-En1/2* CKOs: n=12, littermate controls: n=13; two-tailed unpaired t-test: t_23_ = 14.35, P < 0.0001). **e**, Quantification of average (left and right hemispheres) SS-AMY tractography in *Atoh1-En1/2* CKOs compared to littermate controls (*Atoh1-En1/2* CKOs: n=12, littermate controls: n=13; Mann-Whitney *U* test: *U* = 0, P < 0.0001). **f**, Quantification of average (left and right hemispheres) HPF-TH tractography in *Atoh1-En1/2* CKOs compared to littermate controls (*Atoh1-En1/2* CKOs: n=12, littermate controls: n=13; two-tailed unpaired t-test: t_23_ = 4.298, P = 0.0003). **g,** Quantification of TH-CN tractography in *Atoh1-En1/2* CKOs (n=12) compared to littermate controls (n=13). Ordinary two-way ANOVA: main effect of genotype (F_1,46_ = 166.5, P < 0.0001), but not of hemisphere (P = 0.3838) or interaction (P = 0.8437); with post hoc two-tailed t-tests with uncorrected Fisher’s LSD for effect of genotype for left hemisphere (t_46_ = 9.265, P < 0.0001) and right hemisphere (t_46_ = 8.985, P < 0.0001). **h**, Quantification of ILM-SS tractography in *Atoh1-En1/2* CKOs (n=12) compared to littermate controls (n=13). Ordinary two-way ANOVA: main effect of genotype (F_1,46_ = 18.16, P < 0.0001), but not of hemisphere (P = 0.1675) or interaction (P = 0.9536); with post hoc two-tailed t-tests with uncorrected Fisher’s LSD for effect of genotype for left hemisphere (t_46_ = 3.055, P = 0.0037) and right hemisphere (t_46_ = 2.972, P = 0.0047). **i**, Quantification of ILM-MO tractography in *Atoh1-En1/2* CKOs (n=12) compared to littermate controls (n=13). Ordinary two-way ANOVA: main effect of genotype (F_1,46_ = 5.585, P = 0.024), but not of hemisphere (P = 0.3553) or interaction (P = 0.8322); with post hoc two-tailed t-tests with uncorrected Fisher’s LSD for effect of genotype for left hemisphere (t_46_ = 1.822, P = 0.0750) and right hemisphere (t_46_ = 1.520, P = 0.1353). **j**, Quantification of ILM-DS tractography in *Atoh1-En1/2* CKOs (n=12) compared to littermate controls (n=13). Ordinary two-way ANOVA: main effect of genotype (F_1,46_ = 11.94, P < 0.0001) and hemisphere (F_1,46_ = 29.03, P < 0.0001), but not of interaction (P = 0.9007); with post hoc two-tailed t-tests with uncorrected Fisher’s LSD for effect of genotype for left hemisphere (t_46_ = 3.055, P = 0.0037) and right hemisphere (t_46_ = 2.972, P = 0.0047). **k**, Quantification of SS-DS tractography in *Atoh1-En1/2* CKOs (n=12) compared to littermate controls (n=13). Ordinary two-way ANOVA: main effect of genotype (F_1,46_ = 32.43, P < 0.0001), but not of hemisphere (P = 0.0662) or interaction (P = 0.7773); with post hoc two-tailed t-tests with uncorrected Fisher’s LSD for effect of genotype for left hemisphere (t_46_ = 4.228, P = 0.0001) and right hemisphere (t_46_ = 3.825, P = 0.0004). **l**, Quantification of MO-SS tractography in *Atoh1-En1/2* CKOs (n=12) compared to littermate controls (n=13). Ordinary two-way ANOVA: main effect of genotype (F_1,46_ = 10.60, P = 0.0021), but not of hemisphere (P = 0.0535) or interaction (P = 0.4680); with post hoc two-tailed t-tests with uncorrected Fisher’s LSD for effect of genotype for left hemisphere (t_46_ = 2.820, P = 0.0071) and right hemisphere (t_46_ = 1.785, P = 0.0809). **m**, Quantification of regional brain volumes in *eCN-DTA* mice (n=5) compared to littermate controls (n=5). Two-tailed unpaired t-tests to test for effect of genotype on MB (t_8_ = 4.935, P = 0.001) and CB (t_8_ = 3.130, P = 0.014), other comparisons P ≥ 0.05. **n**, Quantification of whole brain volume in *eCN-DTA* mice compared to littermate controls (n=5 per genotype; two-tailed unpaired t-test: t_8_ = 6.346, P = 0.0002). **o**, Quantification of forebrain and midbrain combined volumes in *eCN-DTA* mice compared to littermate controls (n=5 per genotype; two-tailed unpaired t-test: t_8_ = 3.055, P = 0.0157). **p**, Quantification of average (left plus right hemispheres) ACA-SS tractography in *eCN-DTA* mice compared to littermate controls (n=5 per genotype; two-tailed unpaired t-test: t_8_ = 7.743, P < 0.0001). **q**, Quantification of average (left plus right hemispheres) HPF-STR tractography in *eCN-DTA* mice compared to littermate controls (n=5 per genotype; two-tailed unpaired t-test: t_8_ = 6.324, P = 0.0002). **r**, Quantification of TH-CN tractography in *eCN-DTA* mice (n=5) compared to littermate controls (n=5). Ordinary two-way ANOVA: main effect of genotype (F_1,16_ = 37.89, P < 0.0001), but not of hemisphere (P = 0.9717) or interaction (P = 0.7233); with post hoc two-tailed t-tests with uncorrected Fisher’s LSD for effect of genotype for left hemisphere (t_16_ = 4.103, P = 0.0008) and right hemisphere (t_16_ = 4.613, P = 0.0003). **s**, Quantification of ILM-SS tractography in *eCN-DTA* mice (n=5) compared to littermate controls (n=5). Ordinary two-way ANOVA: no main effect of genotype (P = 0.6080), hemisphere (P = 0.9875) or interaction (P = 0.8883). **t**, Quantification of ILM-MO tractography in *eCN-DTA* mice (n=5) compared to littermate controls (n=5). Ordinary two-way ANOVA: no main effect of genotype (P = 0.9440), hemisphere (P = 0.9600) or interaction (P = 0.8281). **u**, Quantification of ILM-DS tractography in *eCN-DTA* mice (n=5) compared to littermate controls (n=5). Ordinary two-way ANOVA: no main effect of genotype (P = 0.6155), hemisphere (P = 0.5876) or interaction (P = 0.4505). **v**, Quantification of SS-DS tractography in *eCN-DTA* mice (n=5) compared to littermate controls (n=5). Ordinary two-way ANOVA: no main effect of genotype (P = 0.2886), hemisphere (P = 0.8391) or interaction (P = 0.5482). **w**, Quantification of MO-SS tractography in *eCN-DTA* mice (n=5) compared to littermate controls (n=5). Ordinary two-way ANOVA: no main effect of genotype (P = 0.8636), hemisphere (P = 0.1664) or interaction (P = 0.1924). Abbreviations: CTX=cerebral cortex; OLF=olfactory bulb; HPF=hippocampal formation; AMY=amygdala; STR=striatum; PAL=pallidum; TH=thalamus; HY=hypothalamus; MB=midbrain; HB=hindbrain; CB=cerebellum; ILM=intralaminar nuclei; SS=primary somatosensory cortex; MO=primary motor cortex; ACA = anterior cingulate cortex. ns, not significant: P ≥ 0.05. Data are presented as mean values ± SD for **a**,**b**,**g**,**h** and mean value ± SEM for **c**-**f**,**i-k**.

**Supplementary Fig. 11.**
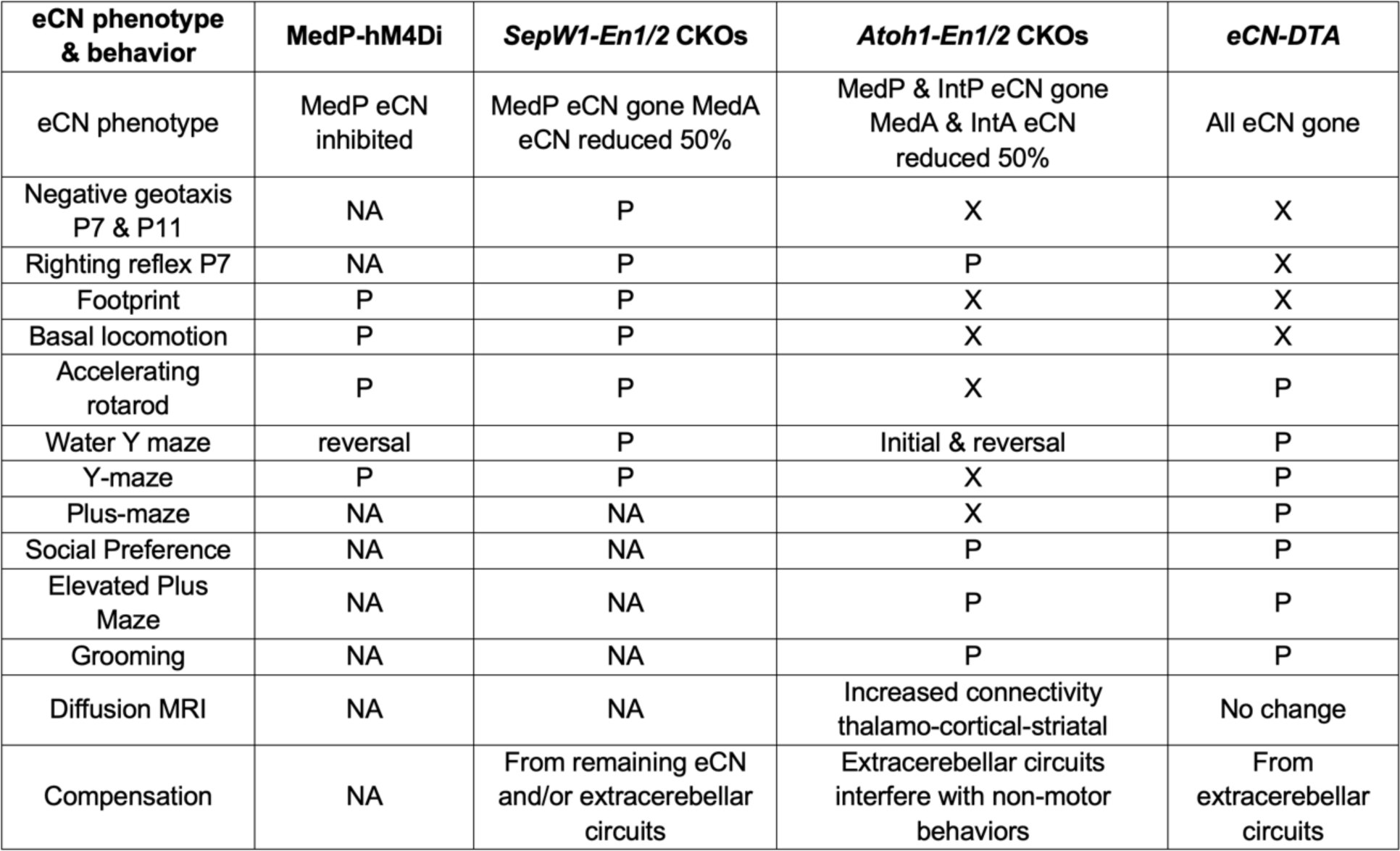
Summary of results. Legend: ✓ = no difference; NA = not applicable; X = impairment.

